# Tracing the stepwise Darwinian evolution of a plant halogenase

**DOI:** 10.1101/2024.12.09.627616

**Authors:** Colin Y. Kim, David W. Kastner, Andrew J. Mitchell, Michael A. Gutierrez, Jocelyn S. Yao, Edwin N. Neumann, Heather J. Kulik, Jing-Ke Weng

## Abstract

Halogenation chemistry is rare in plant metabolism, with the chloroalkaloid acutumine produced by Menispermaceae species being the only well characterized example, involving a specialized dechloroacutumine halogenase (DAH) from the iron(II)- and 2-oxoglutarate-dependent dioxygenase (2ODD) superfamily. While DAH is presumed to have evolved from an ancestral 2ODD enzyme, the broader question of how new enzymes arise through Darwinian processes, such as the birth of DAH in Menispermaceae, remains a fundamental challenge in understanding metabolic evolution. Here, we investigate DAH’s evolutionary trajectory using the chromosomal-level genome assembly of *Menispermum canadense*. By analyzing the genomic context of *DAH* in *M. canadense* and syntenic regions in related plants, we show that *DAH* evolved through tandem duplication of an ancestral *flavonol synthase* (*FLS*) gene, followed by a series of neofunctionalization and gene loss events. Through structural modeling, molecular dynamics simulations, and site-directed mutagenesis, we identify residue changes enabling the transition from FLS to DAH. This functional switch required traversing a complex evolutionary landscape where adaptive peaks were separated by deep fitness valleys. Our work illustrates how new enzymatic functions can arise through lineage-specific evolutionary pathways that gradually reshape the active site architecture through permissive mutations, ultimately enabling mechanism-switching mutations that establish novel catalytic activities.

## Introduction

Halogenation in nature is a valuable chemical transformation that enhances the diversity and functionality of natural products, contributing to their medicinal potency^1–3^. While plants have developed elaborate specialized metabolic pathways to produce a dazzling array of structurally diverse secondary compounds, halogenation chemistry is rarely observed. To date, dechloroacutumine halogenase (DAH), which catalyzes the terminal chlorination in acutumine biosynthesis in the Menispermaceae family, stands out as the only characterized halogenase across all land plants^4^. The limited occurrence of halogenated plant natural products presents an opportunity to expand nature’s chemical space through biocatalytic halogenation, potentially yielding new pharmaceutically relevant compounds. However, with DAH being the only characterized plant halogenase, understanding how these enzymes evolved remains a key challenge for rationally designing new halogenases. DAH thus offers a unique window into understanding how nature can evolve new catalytic functions through Darwinian processes.

DAH is an iron(II)- and 2-oxoglutarate (Fe(II)/2OG)-dependent halogenase (2ODH) of plant origin, belonging to the large superfamily of iron(II)- and 2-oxoglutarate-dependent dioxygenases (2ODDs)^4^. 2ODDs are found across all kingdoms of life that utilize a non-heme ferrous cofactor for their radical oxidative catalysis^5^. These enzymes target otherwise inactive sp^3^ and sp^2^ C-H bonds at a highly conserved iron-binding facial triad (HxD/EX*_n_*H) in their active site for a diverse range of reactivities, with hydroxylation being by far the most common^6^. 2ODHs are evolutionarily derived from 2ODDs and harbor a mechanistic active-site substitution, replacing the key acidic residue Asp/Glu of the facial triad in 2ODDs with a Gly/Ala^7^. This allows the halide ligand to occupy an iron coordination site, and in turn elicits halogenation chemistry^8^. While the key mechanistic D-to-G mutation is only observed once in plants to yield DAH, a handful of 2ODHs have been characterized in bacteria including WelO5 from welwitindolinone biosynthesis in *Hapalosiphon welwischii*^8^ and a group of amino acid halogenases from *Streptomyces cattleya*^9^. Leveraging this highly conserved mechanism along with diverse reaction outcomes, 2ODD has been recognized as a promising biocatalytic scaffold for directed evolution over the past decade to engineer regio-stereospecific halogenases and azidases^10–12^. In particular, a bacterial Fe(II)/2OG hydroxylase, MBT76 in lysine metabolism of *Streptomyces*, was converted to a lysine halogenase inspired by sequence and structural comparison between a pair of closely related hydroxylase and halogenase^13^. While activity-guided directed evolution of Fe(II)/2OG hydroxylases has been explored, the natural evolutionary pathway leading to 2ODHs has not been studied in any host organism.

Plant genomes, with their large size and frequent duplication events, serve as rich but largely untapped reservoirs of evolutionary history, preserving the genomic footprints of how biosynthetic genes acquire new functions to expand metabolic diversity^14^. For example, the metabolic diversity of acylsugars found in cultivated tomato roots is driven by gene duplication followed by neofunctionalization within a conserved biosynthetic gene cluster across various cultivated Solanaceae genomes^15^. As the only known 2ODH in plant metabolism, DAH’s unique emergence in Menispermaceae provides a valuable opportunity to investigate the molecular mechanisms underlying its evolution, with potential implications for designing biocatalysts for targeted C-H functionalization. Here, we report a chromosomal-level genome assembly of *M. canadense* and trace DAH’s evolution from its progenitor flavonol synthase (FLS), a Fe(II)/2OG desaturase, through two evolutionary intermediate pseudogenes preserved in the genome. Using structural models of DAH and FLS, we identify a set of amino acid substitutions and insertions necessary for the functional transition from FLS to DAH. Through mechanism-guided engineering of additional selected plant 2ODDs, we validate the rugged evolutionary landscape leading to DAH, which in turn highlights the role of lineage-specific evolutionary events in enabling subsequent mechanism-switching mutations in enzyme evolution.

## Results

### Chromosomal-level assembly of the *M. canadense* genome

To shed light on the evolutionary history of *DAH*, we sequenced and assembled the genome of *M. canadense*. This species is diploid (2*n* = 2*x* = 52), with an estimated genome size of approximately 958 Mb according to *k*-mer (*k* = 19) analysis (Supplementary Fig. 1) and 925 Mb according to flow cytometry (Supplementary Fig. 2). We extracted high-molecular weight genomic DNA from leaf tissues and performed highly accurate long-read sequencing (Pacific Biosciences). The initial draft assembly using HiFi circular consensus sequencing reads yielded a size of approximately 970 Mb with 1,327 contigs and an auN of 15.6 Mb (Supplementary Fig. 3, Supplementary Table 1). To improve the assembly, we employed the high-throughput chromosome conformation capture (Hi-C) technique to scaffold 890.5 Mb of the assembly onto 26 pseudochromosomes with an auN of 34.2 Mb, covering 96.28% of the expected genome size from flow cytometry (Fig. 1a, Supplementary Figs. 4 and 5, Supplementary Table 2). We mapped Illumina short read sequencing data onto this reference genome, resulting in 98.87% of the 1.821 billion total reads being mapped onto 26 pseudochromosomes (Supplementary Table 3).

**Fig. 1.**
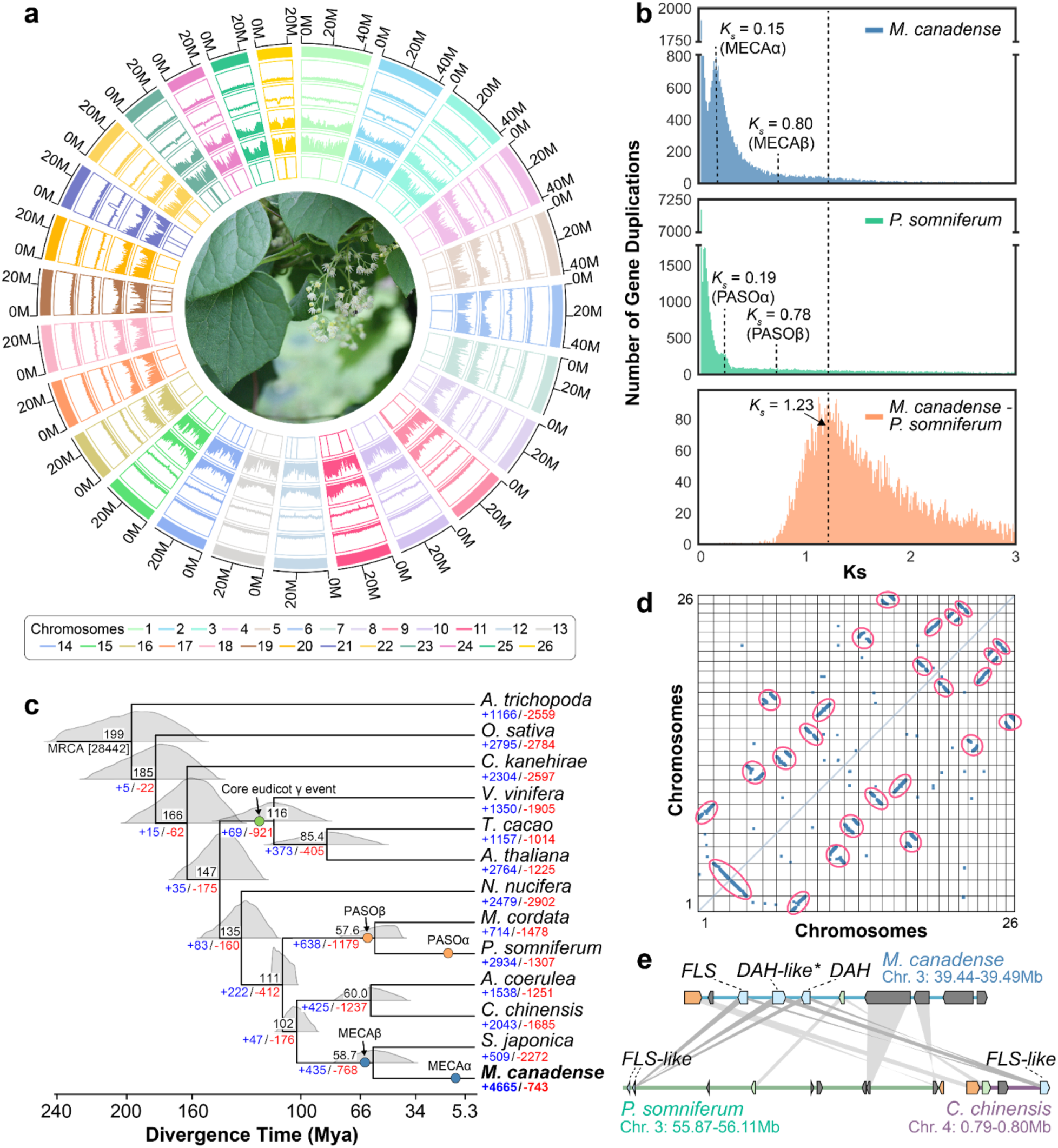
Chromosomal-level genome assembly of *M. canadense*. (**a**) Circos plot displaying genome assembly statistics for each pseudochromosome, including chromosome size, repeat element density, GC content, gene density, transcript coverage, and assembly gaps (from outer to inner tracks). An image of *M. canadense* is positioned at the center of the plot. (**b**) Histogram of *Ks* distribution based on all-to-all blast of total orthologous genes in *M. canadense* and *P. somniferum* indicating WGDs. Mixtools analysis revealed *Ks* distribution peaks at 0.15 (MECAα) and 0.80 (MECAβ) for *M. canadense*, 0.19 (PASOα) and 0.78 (PASOβ) for *P. somniferum*, and 1.23 for the divergence of *M. canadense* and *P. somniferum*. (**c**) Phylogenetic divergence time of *M. canadense* compared to 12 other angiosperm species; *Amborella trichopoda*, *Oryza sativa*, *Cinnamomum kanehirae*, *Vitis vinifera*, *Theobroma cacao*, *Arabidopsis thaliana*, *Nelumbo nucifera*, *Macleaya cordata*, *Papaver somniferum*, *Aquilegia coerulea*, *Coptis chinensis*, and *Stephania japonica*. Black number at the top of each tree node indicates the mean divergence time million years ago (Mya). The blue and red numbers at the bottom of each tree node indicate gene family expansion and contraction, respectively, with the most recent common ancestor (MRCA) having 28442 gene families represented across the species in the phylogenetic tree. Green circle indicates the relative timing of the core eudicot γ event, whereas orange circles correspond to two WGD events present in *P. somniferum*. Blue circles indicate the relative timing of two predicted WGD events in *M. canadense*. (**d**) Dot plot visualization of intragenomic syntenic regions within the *M. canadense* genome showcasing WGD. A total of 10,745 genomic pairs are shown with a C-score cutoff of 0.90. Red circles indicate WGD-pairs. (**e**) Microsynteny comparison of the genomic loci containing DAH in *M. canadense* (chromosome 3; 39.44-39.49 Mb) and *P. somniferum* (chromosome 3; 55.87-56.11 Mb) or *C. chinensis* (chromosome 4; 0.79-0.80 Mb) genomes.

Based on this assembled *M. canadense* genome, we annotated 65,843 protein-coding genes with homologous alignments, *ab initio* gene models, and transcriptome data from our previous study^4^. In total, 99.9% of the transcripts longer than 1,000 bp could be mapped onto the *M. canadense* genome with >50% sequence coverage (Supplementary Table 4). To assess its completeness, the BUSCO^16^ analysis was conducted, which demonstrated a score of 99.4% (1604 out of 1614 conserved genes) (Supplementary Fig. 6). In addition, the OMArk^17^ analysis for its proteome revealed its high completeness with a score of 96.37% (7832 out of 8127 conserved hierarchical orthologous groups) (Supplementary Fig. 7). Moreover, Merqury^18^ analysis of the Hi-C scaffolded assembly showed a completeness score of 83.97% with the quality value (QV) of 59.66 and low error rate of 1.08x10^-6^ (Supplementary Table 5). The final assembly statistics are shown in a Circos plot (Fig. 1a). The highly contiguous and complete genome of *M. canadense* enables us to leverage bioinformatic analyses to investigate the genomic distribution and evolution of enzymes involved in acutumine-type alkaloid biosynthesis.

### Phylogenomic analysis of the *M. canadense* genome

To enable comparative genomic study of *M. canadense* relative to other plants with sequenced genomes, we first used OrthoFinder^19^ to obtain orthologous groups between *M. canadense* and 12 other angiosperm species (Supplementary Table 6). Out of 55,164 curated genes from the *M. canadense* genome, 47,394 (85.9%) genes were placed in orthologous groups and 15,244 (27.6%) in *M. canadense*-specific groups (Supplementary Table 7). Using 249 single-copy orthologous sequences across all 13 species, we constructed a maximum-likelihood phylogenetic tree. The coalescence-based analysis suggests that *M. canadense* shares a recent common ancestor with *Stephania japonica*, a Menispermaceae plant that does not produce acutumine (Fig. 1c). These Menispermaceae plants share a common ancestor with two Ranunculaceae species, *Coptis chinensis* and *Aquilegia coerulea*, which is sister to the Papaveraceae family containing *Papaver somniferum* and *Macleaya cordata* (Fig. 1c).

1. *M. canadense* belongs to the Ranunculales, a basal eudicot group evolved in parallel to the core eudicots (Fig. 1c). Using MCMCtree^20^ with molecular clock calibrations, we estimated that eudicots diverged from early angiosperms ∼166 million years ago (Mya), with basal eudicots and core eudicots diverging ∼19 Mya afterwards. Within Ranunculales, Papaveraceae species diverged ∼111 Mya, Menispermaceae species diverged from Ranunculaceae ∼103 Mya, and *M. canadense* diverged from *S. japonica* ∼59 Mya (Fig. 1c). Analysis of gene family evolution revealed 5,507 expanded and 2,518 contracted families in *M. canadense*, compared to 1,351 expanded and 4,047 contracted in *S. japonica*, 2,875 expanded and 3,929 contracted in *C. chinensis*, and 3,932 expanded and 3,317 contracted in *P. somniferum* (Fig. 1c). Among 158 gene families annotated as 2ODD, we observed 47 expansions and 22 contractions in *M. canadense* (Supplementary Fig. 8).

### Whole-genome duplication events indicated by the *M. canadense* genome

Ancient whole-genome duplication (WGD) provides a primary mechanism for generating copy number variation of biosynthetic genes in plants, and can be detected through analysis of paralogous gene age distributions^21,22^. Given the large gene family expansions in *M. canadense*, we investigated whether Menispermaceae-family plants had undergone lineage-specific WGD events that might have contributed to the emergence of *DAH*. We analyzed potential WGD events by calculating the number of synonymous mutations per synonymous site (*K*_S_) for paralogous genes^22,23^, followed by peak detection using a mixture model from the mixtools R package^24^. The distribution of reciprocal best hit (RBH) paralogous gene pair *K*_S_ values exhibits peaks at 0.80 in *M. canadense*, 0.89 in *S. japonica*, 0.78 in *P. somniferum*, and 0.72 in *C. chinensis*, suggesting ancient WGD events (Fig. 1b and Supplementary Fig. 9). This WGD event, designated as MECAβ, occurred after the divergence of *M. canadense* from *P. somniferum* (*K*_S_ = 1.23) and *C. chinensis* (*K*_S_ = 1.10), but before its divergence from *S. japonica* (*K*_S_ = 0.7) (Fig. 1b and Supplementary Fig. 9). This indicates that an ancient WGD event occurred in the evolutionary history of *M. canadense* after Menispermaceae split from other Ranunculales plants, independent of other Ranunculales WGD events. This WGD event is evident in the syntenic dotplot of the *M. canadense* genome using paralogous gene pairs (Fig. 1d, Supplementary Fig. 10a).

To examine additional rounds of ancient WGD, we compared syntenic depth ratios between *M. canadense* and *Amborella trichopoda*. We observed a four-to-one syntenic depth ratio, indicating that a single *A. trichopoda* genomic region aligns to four *M. canadense* blocks (Supplementary Fig. 10b). Since *A. trichopoda* has not experienced any WGD after the ancestral angiosperm genome duplication event^25^, this ratio suggests *M. canadense* underwent two rounds of ancient WGD since their last common ancestor. Furthermore, we observed two-to-one and four-to-four syntenic depth ratios between *M. canadense* and *C. chinensis* and *P. somniferum*, respectively (Supplementary Fig. 10c-e). Previous analyses show that *C. chinensis* experienced one WGD event (*K*_S_ = 0.72), while *P. somniferum* underwent both a Papaveraceae-specific WGD (PASOβ; *K*_S_ = 0.78) and an additional *Papaver*-specific WGD^26^ (PASOα; *K*_S_ = 0.19) (Fig. 1b, Supplementary Figs. 9 and 10). Similarly, *M. canadense* experienced an additional WGD event (MECAα; *K*_S_ = 0.15) after the Menispermaceae-specific WGD (MECAβ; *K*_S_ = 0.80) (Fig. 1b, Supplementary Fig. 9). The two-to-one syntenic depth ratio between *M. canadense* and *S. japonica* and two-to-two ratio between *S. japonica* and *C. chinensis*^27^ support that MECAβ is shared within Menispermaceae while MECAα is specific to the *Menispermum* genus (Supplementary Fig. 10g, h).

### Genomic signatures of *DAH* evolution

To probe the evolutionary origin of *DAH*, we examined the genomic loci containing *DAH* and its closely related homologs in the *M. canadense* genome. We found that the *DAH* gene, located on chromosome 3, is proximal to two paralogous genes: *DAH-like*, positioned next to *DAH*, contains the halogenase sequence motif HxGX*_n_*H and shares 95.4% amino acid sequence identity with DAH; and *flavonol synthase* (*FLS*), positioned next to *DAH-like*, contains the hydroxylase sequence motif HxDX*_n_*H and shares 64.3% amino acid sequence identity with DAH (Fig. 2a-c). Syntenic analysis revealed corresponding regions in *P. somniferum* and *C. chinensis* genomes, containing tandemly duplicated *FLS-like* genes in *P. somniferum* and a single *FLS-like* gene in *C. chinensis*, but no *DAH-like* genes in either case (Fig. 1e). Additionally, the WGD event (MECAα) within the *M. canadense* genome produced a duplicated region of this locus on chromosome 2 (Fig. 2a). Unlike the tandem arrangement on chromosome 3, this duplicated region contains only one gene, named *dechloroacutumine hydroxylase-like* (*DAHy-like*), which contains the hydroxylase sequence motif HxDX*_n_*H and shares 88.5% amino acid sequence identity with DAH (Figs. 1d and 2a-c). Similarly, WGD (PASOα) within the *P. somniferum* genome duplicated the *FLS-like* gene locus, resulting in a total of four *FLS-like* genes (Supplementary Fig. 11).

**Fig. 2.**
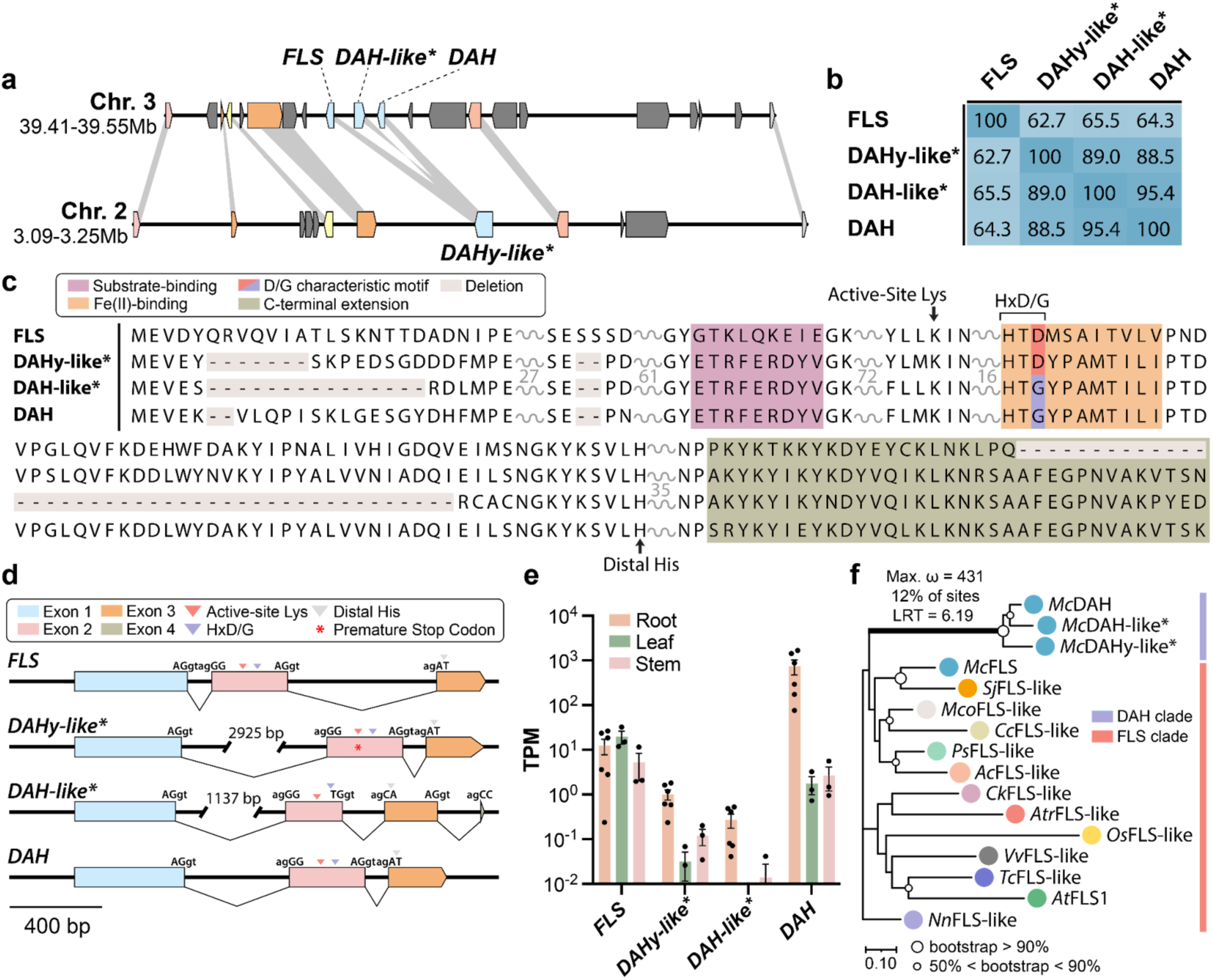
Synteny-based analysis of *DAH* locus and its evolutionary context. (**a**) WGD region on chromosome 3 and 2 containing *FLS*, *DAH*, and its paralogous genes. Asterisk denotes pseudogenes. (**b**) Amino-acid sequence identity matrix of FLS, DAHy-like, DAH-like, and DAH calculated by local pairwise alignments using EMBOSS supermatcher^43^. Asterisk denotes non-functional proteins. (**c**) Multiple sequence alignment of FLS, DAHy-like, DAH-like, and DAH found on chromosomes 2 and 3. Asterisk denotes non-functional proteins. (**d**) Intron-exon structure of *FLS*, *DAHy-like*, *DAH-like*, and *DAH*. All gene structures are shown in 5’-3’ orientation. All genes share the first two intron-exon splice junction sequences. Asterisk denotes pseudogenes. (**e**) Transcript per million (TPM) values of tissue specific RNA sequencing reads of *M. canadense* mapped onto *FLS*, *DAHy-like*, *DAH-like*, and *DAH*. TPM values are shown at log_10_ scale. Data are presented as mean values +/- standard error of the mean with 6 biological replicates of root tissues, 3 of leaf and 3 of stem. (**f**) Maximum-likelihood phylogenetic tree of DAH, its paralogs and FLS orthologs from select angiosperm species represented in Figure 1c. Bootstrap statistics (200 replicates) are indicated at the tree nodes. The scale measures evolutionary distance in substitutions per amino acid. The branch leading to the common ancestor of DAH and FLS recovered with significant (p = 0.0161) evidence of positive selection, with 12% of sites showing strong directional selection (ω or max d_N_/d_S_ = 431) according to the adaptive branch-site REL test for episodic diversification (aBSREL) method. Asterisk denotes non-functional proteins.

To investigate the evolutionary relationship among *DAH*, *DAH-like*, *DAHy-like*, and *FLS* in *M. canadense*, we examined their exon-intron structures and found that the splicing junction sequences between *FLS*, *DAHy-like*, and *DAH* are conserved (Fig. 2d). For *DAH-like*, only the splicing junction sequences for the first intron are conserved. At the protein level, multiple sequence alignment shows that DAH, DAH-like, and DAHy-like all contain a 13-amino-acid C-terminal extension compared to FLS, while DAH-like has a 30-amino-acid internal truncation compared to DAH in a region likely critical for protein folding and catalysis, suggesting a loss of function (Fig. 2c). This truncation is found in the second exon of *DAH-like*, which contains unique splicing junction sequences for its second intron and a unique third exon (Fig. 2d and Supplementary Figs. 12 and 13). Both *DAHy-like* and *DAH-like* genes have elongated first introns compared to *FLS* and *DAH*, suggesting a negative correlation with transcriptional expression or mRNA maturation process as a result of gene duplication (Fig. 2d). Moreover, analysis of tissue-specific transcriptome datasets from our previous study^4^ revealed that *DAHy-like* and *DAH-like* exhibited generally lower transcript-per-million (TPM) values than *FLS* and *DAH* across all three tissues (Fig. 2e). Interestingly, mapped RNA-sequencing reads for *DAHy-like* indicate that an independent *M. canadense* sample carries a naturally occurring allele containing two single nucleotide polymorphisms (SNPs) at the codon sequence of Lys200 (’AAA’ to ’TAG’), leading to a premature stop codon (Supplementary Fig. 14). Based on these observations, we surmise that *DAHy-like* and *DAH-like* are relics of evolutionary intermediates in the trajectory linking the *FLS* progenitor to *DAH*. The low expression levels, large truncation in *DAH-like*, and natural presence of an allele containing a premature stop codon in *DAHy-like* suggest that these two *DAH* paralogs are likely no longer functional and are undergoing pseudogenization in the *M. canadense* genome.

To further resolve the phylogenetic relationships between FLS and the three DAH paralogs, we constructed a maximum-likelihood phylogenetic tree using DAH, DAH-like, DAHy-like, and representative FLS orthologs from the orthologous groups generated in *M. canadense* and 11 relevant angiosperm species (Fig. 2f). DAH, DAH-like, and DAHy-like appear to form a distinct clade separate from the FLS clade. Moreover, the branching pattern shows that proteins with the acquired halogenase sequence motif HxGX*_n_*H (i.e., DAH and DAH-like) were likely derived from an ancestral DAHy-like protein, which harbored the hydroxylase motif HxDX*_n_*H. Overall, these phylogenetic analyses suggest that the characteristic D-to-G mechanism-switching mutation observed in DAH and DAH-like occurred in an evolutionary intermediate paralogous to DAHy-like after it had significantly diverged from FLS. Additionally, it is noteworthy that FLS homologs can be found in angiosperms ranging from early monocots to core eudicots (Fig. 2f). This indicates that flavonol and flavonoid biosynthesis are highly conserved across these species, yet the divergence towards DAH appears limited to Menispermaceae plants alone. To further test for positive selection of DAH, we performed a targeted molecular adaptation analysis using the coding nucleotide sequences of the genes from this phylogenetic tree for the adaptive branch-site REL test for episodic diversification (aBSREL). Our analysis revealed that 12% of the nucleotide coding sequence sites along the branch leading to the DAH clade containing *McDAH*, *McDAH-like*, and *McDAHy-like* exhibited a statistically significant signal of episodic positive selection, with *d*_N_/*d*_S_ > 1 (ω or max *d*_N_/*d*_S_ = 431) (Fig. 2f). This finding suggests a signal of neofunctionalization, where DAH has acquired distinct functional capabilities from its ancestral FLS role.

### Structural basis for the evolution of DAH from an FLS progenitor

As genomic signatures in *M. canadense* suggest that chlorinated alkaloids in plants evolved from flavonol biosynthesis, we next sought to examine possible mutational trajectories and the structural basis underlying the FLS-to-DAH functional transition. We first characterized the biochemical functions of FLS and DAH using purified recombinant proteins against their native substrates in *in vitro* assays (Supplementary Fig. 15). FLS catalyzes desaturation of dihydroflavonols, dihydrokaempferol, and dihydroquercetin to produce flavonols, kaempferol, and quercetin, respectively (Fig. 3a, c and Supplementary Fig. 16). Surprisingly, a trace amount of kaempferol was also detected when DAH was assayed against dihydrokaempferol, suggesting it still retains a low level of the ancestral FLS activity (Supplementary Fig. 17). Likewise, we examined whether FLS harbors any reactivity against dechloroacutumine. While DAH shows 2OG-dependent chlorinase activity on dechloroacutumine to produce acutumine, FLS does not exhibit any oxidase or halogenase activity on dechloroacutumine (Fig. 3b, d). Furthermore, the expression of pseudogenes encoding evolutionary intermediates, DAH-like and DAHy-like, resulted in insoluble protein during the purification procedure, consistent with the observation of pseudogenization.

**Fig. 3.**
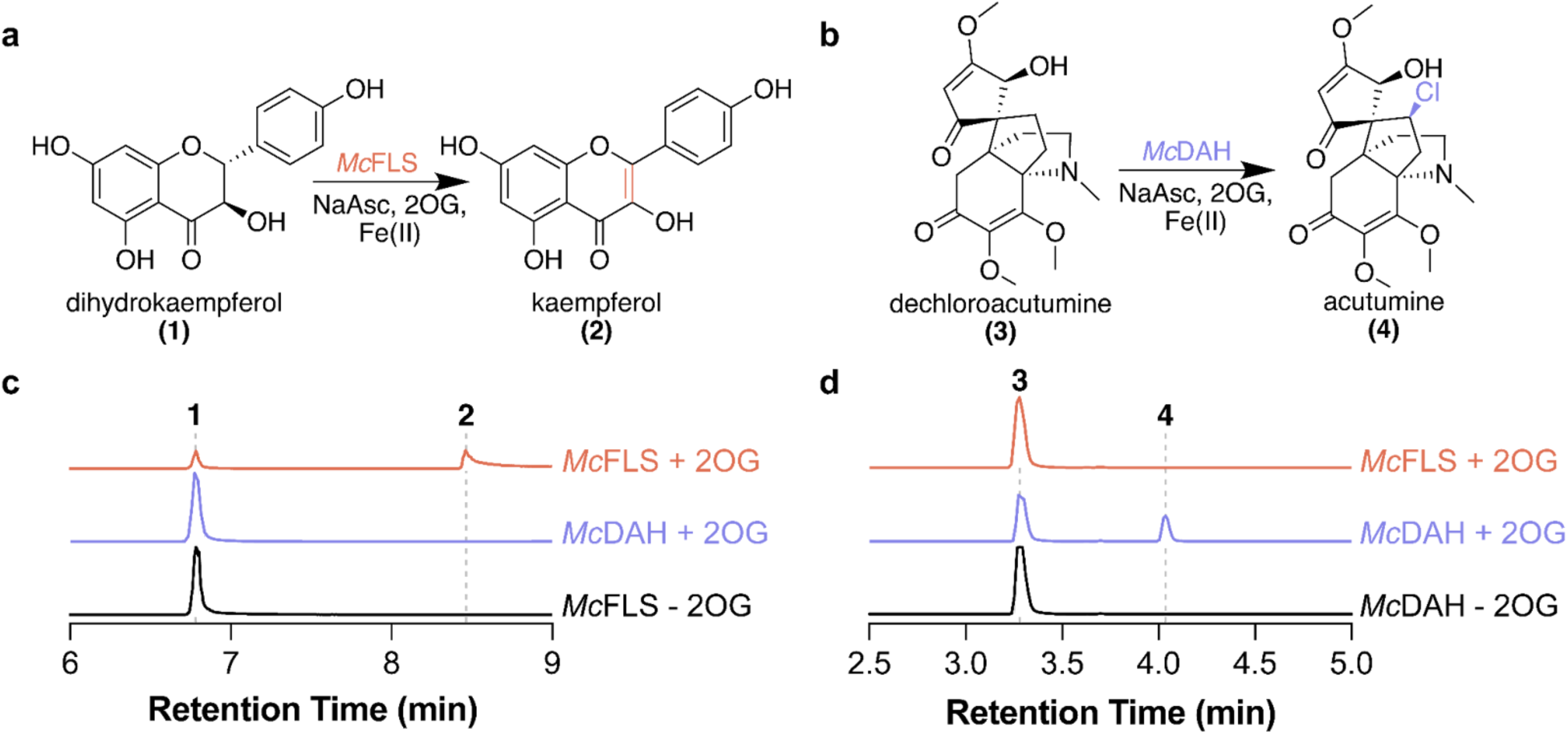
*In vitro* biochemical assay of FLS and DAH. (**a**) Reaction schematic of the conversion of dihydrokaempferol; **1** to kaempferol; **2** using flavonol synthase (*Mc*FLS). (**b**) Reaction schematic of the conversion of dechloroacutumine; **3** to acutumine; **4** using dechloroacutumine halogenase (*Mc*DAH). (**c**) Combined LC-MS extracted ion chromatograms (EICs) of 287.05594 *m/z*; **1** = [M-H]^-^ and 285.04050 *m/z*; **2** = [M-H]^-^. EICs show the *in vitro* activity of *Mc*FLS that desaturate **1** to **2** in a 2OG-dependent manner. *Mc*DAH produces trace amounts of **2** in the reaction (**Supplementary** Fig. 17). (**d**) Combined LC-MS extracted ion chromatograms (EICs) of 364.17526 *m/z*; **3** = [M+H]^+^ and 398.13629 *m/z*; **4** = [M+H]^+^. EICs show the *in vitro* activity of *Mc*DAH that chlorinate **3** to **4** in a 2OG-dependent manner, while *Mc*FLS exhibits no production of **4**.

To examine the structural features contributing to the functional divergence between FLS and DAH, we obtained structural models of DAH and FLS using AlphaFold2^28^, followed by docking of dechloroacutumine and dihydrokaempferol into their active sites, respectively (Supplementary Fig. 18). We estimated the positioning of Fe(II), Cl^-^ anion, and 2-oxoglutarate (2OG) based on a structural alignment with a previously reported crystal structure of the *Arabidopsis thaliana* anthocyanidin synthase (*At*ANS; PDB: 2BRT)^29^ (Supplementary Fig. 19). From the structural alignment, we noticed that the characteristic D-to-G mutation in DAH creates two open coordination positions, axial or equatorial to His224, for the oxo/hydroxo ligand of the Fe(IV)=O/Fe(II)-OH unit to occupy (Supplementary Fig. 20). Thus, we derived both axial-oxo and equatorial-oxo conformational models to represent the reactive state of DAH with Fe(IV)-oxo and succinate (Supplementary Fig. 20). We then ran extended molecular dynamics (MD) simulations of both DAH isomers with constraints favoring either an acute or obtuse oxo-Fe(IV)-H target angle (Supplementary Figs. 20-23).

Previous examination of available crystallographic and spectroscopic data revealed that halogenases prefer obtuse angles and hydroxylases prefer acute angles^30^. When analyzing the angle and distance preferences of the spectroscopically guided MD simulations, we noticed that FLS simulations preferred the acute conformation, while DAH preferred the obtuse conformation (Supplementary Figs. 24-25). Finally, the MD simulations were clustered and the centroid of the clustered simulations was QM/MM optimized (Fig. 4a, Supplementary Fig. 21). Upon inspection of the DAH model in the equatorial-oxo conformation, we found Thr231 and Asn262 were located proximal to the Fe(IV)-oxo and speculated about their ability to perform second-sphere interactions that favor halogenation over hydroxylation outcome, similar to Ser189 and Asn219 residues in bacterial 2ODHs WelO5^8^ and BesD^9^, respectively (Supplementary Fig. 26). However, when *in vitro* assays were performed using T231A and N262A DAH mutants against dechloroacutumine, we observed no significant change in acutumine production, suggesting that the active site of DAH does not depend on a hydrogen bond donor for the oxo to achieve its halogenation specificity (Supplementary Fig. 26).

**Fig. 4.**
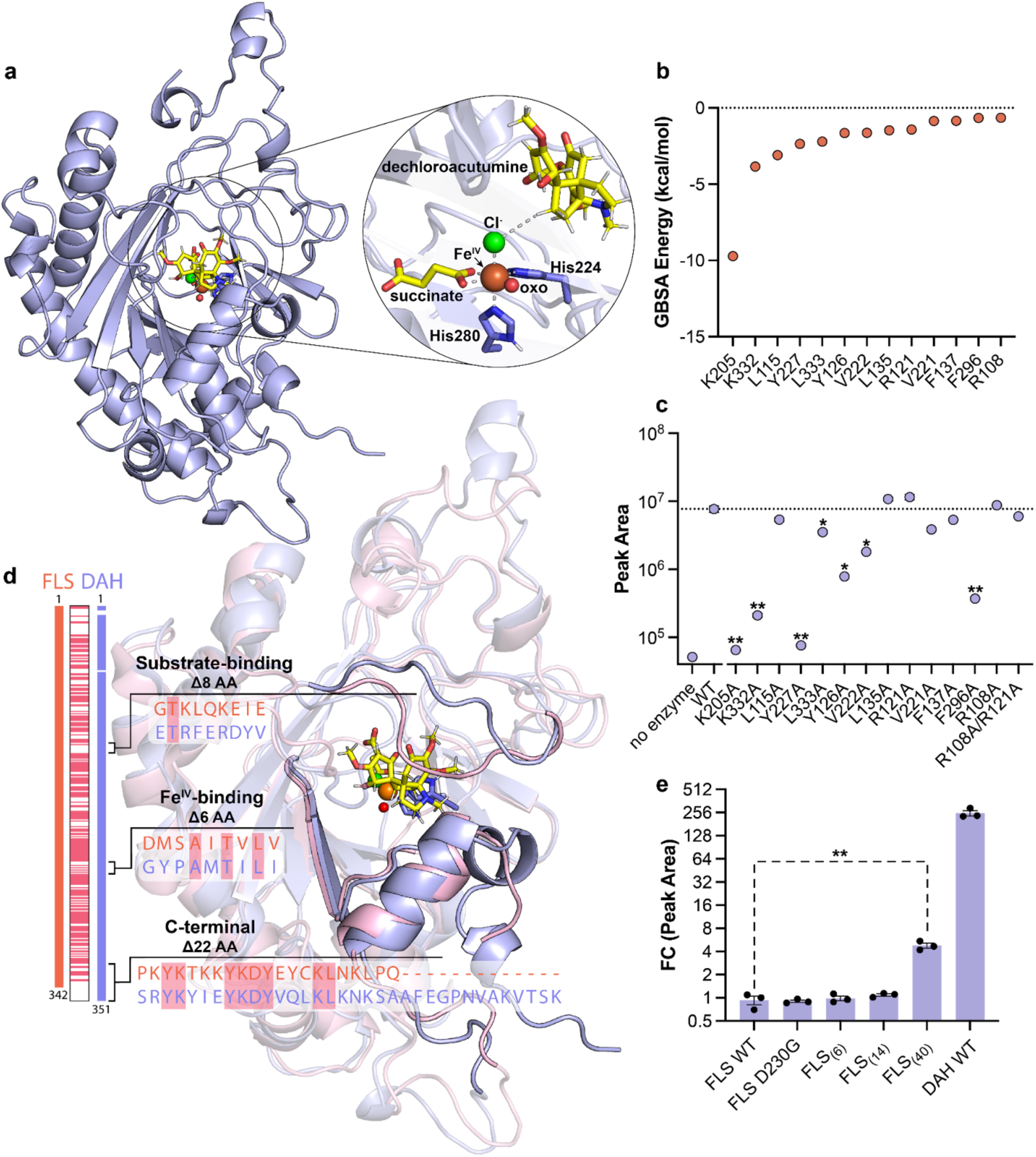
Structural model-guided characterization and molecular evolution of DAH. (**a**) QM-MM-optimized structural model of *Mc*DAH generated with constraints that favor an obtuse oxo-Fe(IV)-H target angle. Predicted active site residues are shown in purple sticks, along with optimized positioning of dechloroacutumine, succinate, Cl anion, and Fe(IV)-oxo species. (**b**) MM-GBSA hydrogen-bonding energy calculations from molecular dynamics simulations of *Mc*DAH structural model. 14 lowest energy residues with their GBSA energy values are shown. (**c**) Catalytic activity of *Mc*DAH alanine mutants. Target mutants were inferred from GBSA energy decomposition analysis and the structural model of *Mc*DAH. LC-HRAM-MS peak areas of acutumine ([M+H]^+^ = 398.13629 *m/z*) are shown for all samples. Dashed line represents the average peak area for acutumine detected using *Mc*DAH wild-type (WT). All assays were performed in triplicates and the error bars represent standard error of the mean. Single asterisk (*) indicates p < 0.01 and double asterisk (**) indicates p < 0.001 compared to the WT peak area of acutumine. (**d**) Sequence and structural alignment of *Mc*DAH and *Mc*FLS structural models. Both structures have been QM/MM optimized starting from centroids of molecular dynamics simulations. (**e**) Structural model-guided design of FLS mutants for minimal halogenase activity. Fold-change (shown in Log_2_-scale) between the LC-HRAM-MS peak areas of acutumine ([M+H]^+^ = 398.13629 *m/z*) in all samples and that of FLS WT sample are shown.

Upon further examination of both equatorial-oxo and axial-oxo DAH conformers, we found that Lys205 was near the substrate-binding pocket and speculated about its involvement in substrate positioning (Fig. 4a, Supplementary Fig. 27). When the K205A DAH mutant was generated and tested *in vitro*, it completely abolished the halogenation activity (Fig. 4c). Subsequently, we performed classical generalized Born energy decomposition analysis (GBSA) on the dominant cluster of the clustered MD simulations and found that K205 contributes -9.7 kcal/mol in energy, suggesting K205 likely plays a role in substrate positioning to maintain the obtuse oxo-Fe(IV)-H target angle (Fig. 4b, Supplementary Figs. 27-31). We also investigated several active-site-lining residues around the substrate binding pocket, notably Lys332, Leu115, Tyr227, Leu333, Arg121, Tyr126, Val222, Leu135, Val221, Phe137, Phe296, and Arg108, which belong to the top 15 most strongly interacting residues based on GBSA energy (Fig. 4a, b and Supplementary Figs. 27-31). To test the potential role of these residues in DAH’s catalytic function, we generated alanine mutants for these active-site-lining residues and tested their activity against dechloroacutumine. We found that K332A, Y227A, L333A, Y126A, V222A, and F296A exhibit significant decreases in halogenation activity, whereas R121A, R108A, L135A, L115A, V221A, F137A, and an R108A/R121A double mutant show no particular difference compared to that of WT DAH (Fig. 4c). These findings suggest that residues with high GBSA dechloroacutamine interaction energies are generally important for DAH halogenation activity, likely through key non-covalent interactions that facilitate favorable substrate positioning, as supported by the diminished activity observed in their alanine mutants (Supplementary Fig. 32).

With the structural models of DAH and FLS, we sought to test the capacity to evolve halogenase activity by incorporating structural characteristics from DAH that are pivotal for halogenation into FLS. First, we generated the FLS D230G mutant and tested its oxidase or halogenase activity against dechloroacutumine and dihydrokaempferol.

However, the FLS D230G mutant did not show any detectable activity against either dechloroacutumine or dihydrokaempferol (Fig. 4e and Supplementary Fig. 33). As evolutionary analyses revealed the importance of other residues beyond the characteristic D-to-G mutation, we identified three structural features that likely contribute to the functional divergence between DAH and FLS: substrate positioning lid-loop (residues 119 to 127), Fe(II)-coordinating β-sheet loop (residues 224-234), and C-terminal helical loop near the substrate pocket (residues 318-351) (Figs. 2c, 4d and Supplementary Fig. 12). FLS mutants containing DAH sequences in each region were produced and analyzed for halogenase activity via *in vitro* assay. The FLS_(6)_ mutant, which involves the replacement of the Fe(II)-coordinating β-sheet loop with its corresponding DAH sequence, does not exhibit any halogenase activity (Fig. 4d, e and Supplementary Fig. 33). Moreover, swapping both the β-sheet loop and substrate positioning loop regions, represented as the FLS_(14)_ mutant, showed no significant changes in halogenase activity (Fig. 4d, e and Supplementary Fig. 33). Interestingly, swapping the C-terminal helical loop region, in addition to the β-sheet loop and substrate positioning loop regions, resulted in the FLS_(40)_ mutant that exhibits detectable halogenase activity towards dechloroacutumine, although such activity is only 1.9% of the wild-type DAH activity (Fig. 4d, e and Supplementary Fig. 33). Conversely, substituting these three structural regions’ corresponding FLS sequences in DAH resulted in increased production of kaempferol when reacted with dihydrokaempferol (Supplementary Figs. 15 and 32).

### Exploration of evolutionary paths to halogenase from other plant 2ODDs

Given the challenges in evolving DAH function from FLS, we further explored this rare characteristic mutation in the widely distributed plant 2ODDs. We selected ten plant 2ODDs that catalyze various oxidation reactions including hydroxylation and *O*-demethylation on metabolites ranging from flavonoids and alkaloids to phytohormones and phytotoxins: *A. thaliana* flavonone-3-hydroxylase (*At*F3H), *Hyoscyamus niger* hyoscyamine-6-β-hydroxylase (*Hn*H6H), *P. somniferum* codeine-*O*-demethylase (*Ps*CODM), *Catharanthus roseus* desacetoxyvindoline-4-hydroxylase (*Cr*D4H), *A. thaliana* dioxygenase-for-auxin-oxidation-1 (*At*DAO1), *A. thaliana* gibberellin-2-oxidase-3 (*At*GA2ox-3), *A. thaliana* gibberellin-3-oxidase-1 (*At*GA3ox-1), *A. thaliana* jasmonate-induced-oxygenase-1 (*At*JOX1), *A. thaliana* downy-mildew-resistant-6 (*At*DMR6), and *Zea mays* DIBOA-glucoside-dioxygenase (*Zm*BX6). Upon examining their multiple sequence alignment with DAH and FLS, we generated the D-to-G or D-to-A mutations at the iron-binding acidic residue for each 2ODD sequence (Fig. 5a). Out of ten plant 2ODDs attempted in this exercise, only two showed alternative reaction outcomes.

**Fig. 5.**
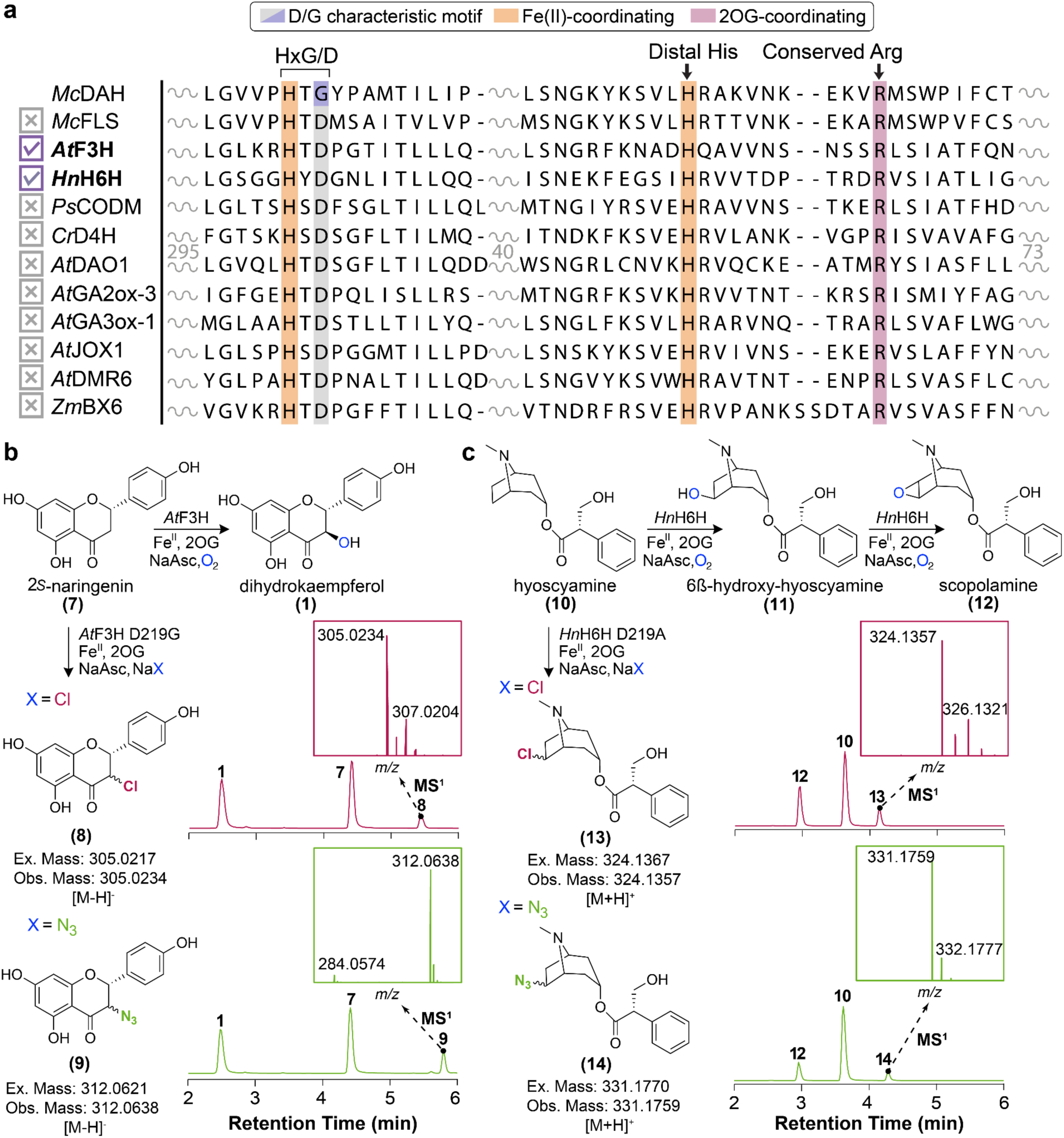
Mechanism-based engineering of plant 2ODDs toward C-H functionalization activities. (**a**) Multiple sequence alignment of select plant 2ODDs with DAH and FLS. Active-site distal histidine is highlighted in orange. Conserved arginine mutation observed in plant 2ODDs is highlighted in red. (**b**) Engineered C-H functionalization of 2*S*-naringenin using *At*F3H D219G variant. Extracted ion chromatograms (EICs) showing the *in vitro* activity of F3H D219G variant that performs alternative anion installations on 2*S*-naringenin, **7** in reaction buffers containing 1 mM NaX, where X = Cl, chlorination of **7** to 9-chloro-naringenin, **8** and X = N_3_, azidation of **7** to 9-azido-naringenin, **9**. The native hydroxylated product, dihydrokaempferol, **1** is also produced as a side product in each EICs. The EICs are scaled to their relative ion intensity. Mass windows used for displaying the EICs: **1**, 287.057 *m/z*; **7**, 271.062 *m/z*; **8**, 305.023 *m/z*; **9**, 312.063 *m/z*. (**c**) Engineered C-H functionalization of hyoscyamine using hyoscyamine**-**6β-hydroxylase D219A variant. EICs showing the *in vitro* activity of H6H D219A variant that performs alternative anion installations on **10** in reaction buffers containing 1 mM NaX, where X = Cl, chlorination of **10** to **13** and X = N_3_, azidation of **10** to **14**. The native hydroxylation product **12** is also produced as a side product in each EICs. The EICs are scaled to their relative ion intensity. Mass windows used for displaying the EICs: **12**, 304.155 *m/z*; **10**, 290.175 *m/z*; **13**, 324.137 *m/z*; **14**, 331.177 *m/z*.

First, we investigated *At*F3H that catalyzes the conversion of 2*S*-naringenin to dihydrokaempferol, a key step during flavonoid biosynthesis in land plants. The *At*F3H D219G variant was recombinantly expressed in *E. coli* and assessed for its activity on 2*S*-naringenin under NaCl conditions, with a direct comparison to assay performed using the WT *At*F3H. (Fig. 5b). We observed a new product peak that corresponds to the *m/z* value of an alternative chlorinated product, which also displayed an isotope distribution consistent with chloride incorporation (Fig. 5b). As previous characterization of DAH has extended catalytic activities to install alternative anions, we tested *At*F3H D219G’s ability to derivatize 2*S*-naringenin using alternative anions like N_3_ ^-^ and Br^-^. In addition to chlorination, we show that *At*F3H D219G is capable of installing azide and bromide when reacted under NaN_3_ or NaBr conditions, respectively (Fig. 5b). Next, we examined *Hn*H6H D219A that catalyzes the hydroxylation of hyoscyamine to scopolamine in tropane alkaloid biosynthesis. Similarly to the *At*F3H D219G case, we observed site-selective installation of chloride and azide on hyoscyamine, as indicated by the new peaks that correspond to the *m/z* and their predicted mass isotope patterns (Fig. 5c).

In both cases, the regio- and stereo-specificity of the anion-installed moieties are not directly measured, but can be reasonably postulated to be at the same -OH group that WT *At*F3H and *Hn*H6H installs onto 2*S*-naringenin and hyoscyamine, respectively. This postulation aligns with the manner in which substrate binding and C-H bond abstraction have been documented in other carrier-independent bacterial 2ODHs and DAH^4,8^. Though *At*F3H D219G and *Hn*H6H D219A exemplify successful cases of D-to-G/A point mutations that facilitate halogenation through opening an anion coordination position in the active site, the mutants still preferred hydroxylation outcome as indicated by higher existing peaks corresponding to their native products (Fig. 5b,c). Furthermore, eight other selected 2ODD D-to-G/A mutants all did not exhibit any halogenation activity against their native substrates (Fig. 5a). It was noted that all mutants were difficult to express and purify as recombinant proteins in *E. coli* (Fig. 5a). This set of experiments suggest that although the D-to-G/A point mutation is key to the mechanistic switch from a hydroxylase to a halogenase in DAH evolution, it is likely preceded by many other mutations that enabled an continuously viable Darwinian mutational trajectory.

## Discussion

The emergence of DAH in *M. canadense* is the only appearance of a halogenase in specialized metabolism across all land plant species reported to date. Our analyses based on the chromosomal-level genome assembly of *M. canadense* capture a series of tandem duplications (TDs) and WGD events that reveal the evolutionary trajectory of FLS to DAH (Fig. 6a). We suspect that the TD of *FLS* initially resulted in two tandem copies of *FLS* on *M. canadense* chromosome 3, where the TD copy is referred to as *FLS-α* (Fig. 6a). *FLS-α* likely acquired inserted sequences encoding the C-terminal extension observed in DAH, presumed to be critical for substrate switch from dihydroflavonol to dechloroacutumine. This evolutionary intermediate is represented as *FLS-β* and whether *FLS-β* harbored DAHy activity is unknown (Fig. 6a). Then, the more recent Menispermum-specific WGD event (MECAα) in *M. canadense* produced a duplicated copy of chromosome 3 as chromosome 2. The resulting *FLS* on chromosome 2 likely underwent gene loss (GL) following WGD, which is a commonly observed evolutionary process in many angiosperms to favor single-copy conservation of critical metabolic flux-controlling genes^31^ (Fig. 6a). At this point, our data are insufficient to conclude whether either *FLS-β* on chromosomes 3 or 2 encoded for DAHy activity in evolutionary history. However, it is clear that *FLS-β* on chromosome 2 eventually underwent pseudogenization to yield *DAHy-like*, which shares 88.5% amino acid sequence identity with DAH while still retaining its hydroxylase sequence motif (Fig. 2b,c, Fig. 6a). We postulate that *FLS-β* on chromosome 3 acquired the functional D-to-G mutation that yielded a functional copy of *DAH* (Fig. 6a). Given that the exon-intron structure of *DAH-like* is significantly different from other paralogous genes, we hypothesize it was produced from an inverted TD event of *DAH* on chromosome 3, which introduced significant truncation in its 2nd exon sequence (Fig. 2c,d, Fig. 6a).

**Fig. 6.**
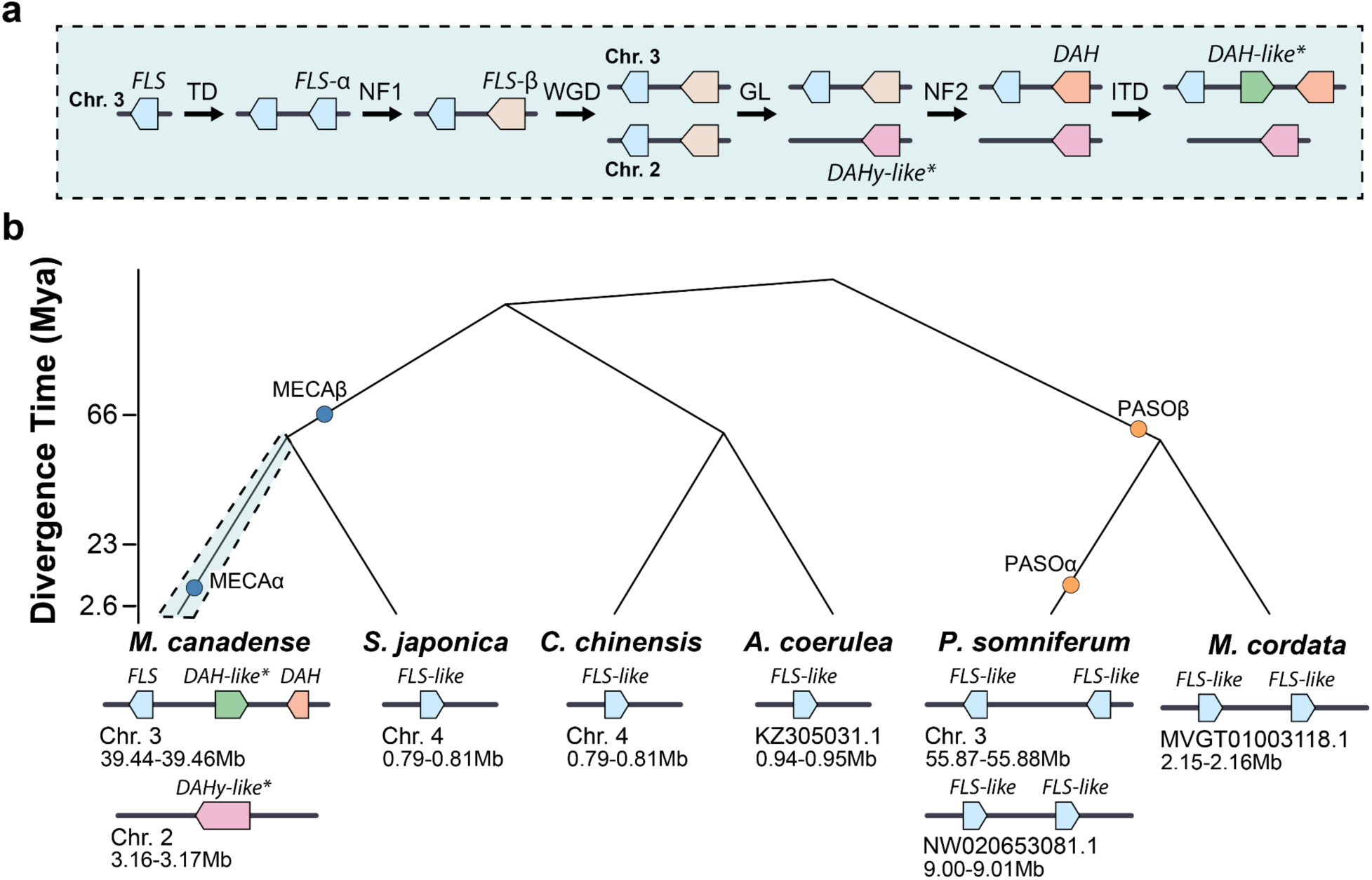
Genomic basis for duplication, neofunctionalization, and peodogenization events related to DAH evolution. (**a**) A proposed evolutionary model of the emergence of *DAH* from a *FLS* progenitor in *M. canadense*. Genomic intermediates not captured by the *M. canadense* genome assembly are colored in wheat. Asterisk denotes pseudogenized genes. Abbreviations: TD; tandem duplication, NF1; neofunctionalization event reflected as C-terminal amino acid extension, WGD; whole genome duplication, GL; gene loss, NF2; neofunctionalization event via D-to-G catalytic switch mutation, ITD; inverted tandem duplication. (**b**) Phylogenetic divergence time of Ranunculales plants as portrayed as a subtree of Fig. 1c. Gene locus containing *FLS* orthologous sequences in *M. canadense*, *S. japonica*, *C. chinensis*, *A. coerulea*, *P. somniferum* and *M. cordata* are shown. Whole genome duplication leading to second locus containing *FLS* orthologs are displayed for *M. canadense* (MECAα) and *P. somniferum* (PASOα).

Perhaps the most remarkable feature of the proposed evolutionary model is the retention of intermediate genes *DAHy-like* and *DAH-like* in the *M. canadense* genome, as it explains several hypotheses in the evolution of DAH. First, the observation of a naturally occurring allele of *DAHy-like* that contains nonsense mutations at the codon of Lys200 (’AAA’ to ’TAG’) leading to a premature stop codon demonstrates a pattern of nonsense-mediated decay (NMD)^32^, as the evolutionary purge of this intermediate gene is reflected by its low expression and mapping of RNA-sequencing reads. This process of pseudogenization is also similarly observed in *DAH-like*, as the internal truncation present in its exon 2 sequence likely contributes to improper protein folding and loss of catalytic activity, consistent with their difficult recombinant expression and purification process. Next, these relics of evolutionary intermediates also provide genomic evidence suggesting the relative order of mutational events responsible for the catalytic function switch from flavonoid to alkaloid biosynthesis. Due to the high conservation of the substrate-positioning loop, Fe(II)-coordinating β-sheet loop, and C-terminal loop regions in *DAHy-like*, *DAH*, and *DAH-like*, we postulate that latent mutations were present in *DAH* before the D-to-G mutation was introduced. We speculate that these mutations enabled the expansion of substrate preference from dihydroflavonol to dechloroacutumine, as demonstrated by DAH’s retention of ancestral FLS activity toward dihydrokaempferol (Supplementary Fig. 17).

The importance of latent mutations in facilitating ultimate mechanism-switching mutations is also underscored by our biochemical investigation of the FLS_(40)_ mutant and select D-to-G/A mutants of select plant 2ODDs. These protein engineering experiments reaffirm the notion that achieving catalytic function conversion between ancestral and derived activities often requires more than simply mutating mechanism-dictating residues. This was previously demonstrated by a study of 311 catalytic variants of wild-type serum paraoxonase (PON1), generated through random mutagenesis, which found that all variants with altered substrate specificity harbored latent mutations at residues unrelated to direct catalytic functionality^33^. A similar observation was also made in the halogenase engineering of the bacterial 2ODD MBT76 from *Streptomyces sp.*, where a DNA shuffling library of the MBT76 D144G variant and a native lysine halogenase, BesD, identified 13 additional distal residues essential for the full functional switch^13^. Although this laboratory-directed evolution exercise of halogenase began with the D-to-G mechanism-switching mutant and subsequently improved desirable activity through additional mutations, our work on the natural evolution of DAH suggests an opposite order of mutational events, where the D-to-G mutation occurred at a much later stage of the evolutionary trajectory leading to the emergence of DAH. Moreover, the difficulty in achieving even minimal halogenase activity in the FLS_(40)_ mutant suggests that the evolutionary landscape between ancestral 2ODDs and derived 2ODHs is highly rugged, with deep fitness valleys^34^, likely limiting the natural evolution of more 2ODHs from the abundant plant 2ODDs.

Previous mechanistic studies of 2ODHs have investigated the significance of specific residues and substrate positioning angles that increase the favorability of chlorination relative to hydroxylation. Our structural modeling of DAH and subsequent GBSA analysis revealed key residues involved in strong non-covalent interactions with the substrate such as Lys205, Tyr227, and Lys332 that were further validated with alanine scanning mutagenesis. Through restrained MD simulations, we also found that DAH favors the obtuse oxo-Fe(IV)-H target angle over acute angles as observed for other halogenases, further explaining its preference for halogenation over hydroxylation (Supplementary Figs. 24 and 25)^30,35,36^. Future efforts in elucidating the crystal structure and catalytic mechanism of DAH will further resolve its reactivity preference as similarly showcased in bacterial 2ODH, BesD^37,38^.

Late-stage, regio- and stereo-selective C-H functionalization of natural product scaffolds represents a valuable “on-demand” reaction but remains a significant challenge with traditional organic synthesis methods. Plant natural products are rarely halogenated, yet a substantial number of small molecule drugs contain a halogen, as halogen atoms play pivotal roles in modulating their pharmacological potency, pharmacokinetic stability, and physicochemical properties^39,40^. Understanding the catalytic and evolutionary mechanisms of halogenating enzymes like DAH and BesD will provide insights for designing enzymes capable of performing these “on-demand” reactions, thereby expanding the functional scope of plant natural products as drug candidates. With the advent of large language models (LLMs) for biocatalyst design, such as ProteinMPNN^41^ and proseLM^42^, future computationally assisted enzyme engineering will harness a wide array of plant 2OGDs to enable the installation of C-H substituents for medicinal compound derivatization. These approaches will likely also illuminate viable evolutionary trajectories that navigate the narrow paths on a rugged evolutionary landscape. The natural evolutionary trajectory from FLS to DAH as preserved in Menispermaceae, will serve as a valuable “ground-truth” resource for LLMs, guiding the design and engineering of more plant 2ODDs into 2ODHs.

## Methods

### Sample Collection, Processing, and Sequencing

1. *M. canadense* plants were purchased from Toadshade Wildflower Farm (Frenchtown, NJ, USA) and grown in a greenhouse at the Whitehead Institute for Biomedical Research (Cambridge, MA, USA). *M. canadense* leaf tissues were harvested and flash-frozen in liquid nitrogen after approximately 6-weeks of growth upon breaking dormancy. High-molecular genomic DNA was extracted using the NucleoBond TakaraBio HMW DNA Extraction kit, quality-controlled by Femtopulse (Agilent Technologies). This DNA was used for PacBio library preparation and SMRT sequencing (Sequel II) for HiFi read generation at the Genomics Core Facility at the Icahn School of Medicine, Mount Sinai, along with PhaseGenomics Hi-C library preparation and Illumina sequencing. We collected 48.57 Gb of SMRT HiFi sequencing data (∼50X coverage) from PacBio Sequel II platform (Pacific Biosciences) used for initial assembly. A total of 270.18 Gb NGS data (∼303X coverage) were generated using Illumina NovaSeq 6000 (Illumina) for quality assessment and mapping onto the final genome assembly. Moreover, 162.63 Gb Hi-C data (∼168X coverage) were generated using Illumina NovaSeq 6000 (Illumina) for scaffolding the initial assembly onto chromosomal-scaffolds. Fastp^44^ (v 0.21.0) was used to assess the quality of sequencing reads.

### Genome Size Estimation by Cell-cytometry and k-mer Analysis

Plant homogenates for analysis of *M. canadense* nuclei were prepared by excising the *M. canadense* leaf sample submerged in nuclei isolation buffer (45 mM MgCl_2_, 30 mM sodium citrate, 20 mM 3-(N-morpholino) propane sulfonic acid (MOPS), pH 7.0) using a razor blade on a petri dish. The resulting homogenates were filtered through a 50 µm disposable filter and loaded onto the flow cytometer (BD Accuri™ C6 Cytometer) to assess the relative fluorescence intensity of nuclei in suspension and estimate the genome size (in picograms of DNA). The resulting flow cytometry measurement for *M. canadense* nuclei was compared to the control sample, *Solanum lycopersicum* (tomato) nuclei to arrive at the estimation of ∼925 Mb for *M. canadense* genome size. Moreover, the genome size of *M. canadense* was computationally estimated using a *k*-mer (k = 19) frequency-based approach with the Illumina paired-end short reads. The software jellyfish (v 2.3.0)^24,45^ was used to count the *k*-mers and visualized using GenomeScope (v2.0)^46^, which was also used for estimating the genome size, resulting in a predicted genome size of ∼958 Mb.

### Genome Assembly, Quality Assessment, and Annotation

To generate the initial contig-level assembly of *M. canadense*, Hifiasm^47^ (v0.15.2) was used with HiFi reads in Hi-C mode and default settings. The initial contig sequence was used as the assembly for scaffolding with Hi-C reads using juicer^48^ (v1.6). 3D-DNA^49^ (v190716) was used to generate Hi-C contact maps with options --editor-repeat-coverage 5 and --splitter-coarse-stringency 30, and the scaffolds were assembled into chromosomes using Juicebox^50^ (v1.11.08). Both the initial contig-level assembly and post-scaffolding chromosomal-level assembly statistics were determined using quast (5.2.0)^51^.

The gene completeness of the assembly were assessed with Benchmarking Universal Single-Copy Orthologs (BUSCO) (v5.1.3)^16^ using hmmsearch (v3.1)^52^, metaeuk (v4.a0f584d)^53^, and embryophyta_odb10 database. Quality assessment of the proteome of *M. canadense* was performed using the OMArk web server^17^ (https://omark.omabrowser.org; OMAmer database: OMA / All.Jul2023 / LUCA | OMAmer version: 2.0.3 | OMArk version: 0.3.0) with default parameters. The *k*-mer (*k* = 20) based genome assessment using merqury^18^ (v1.3) was performed with the post-scaffolded chromosomal-level assembly of *M. canadense* including unscaffolded contigs and 48.57 Gb of SMRT HiFi sequencing data. For counting RNA-seq reads mapped to the 26 pseudo-chromosomes, bwa-mem (v2.2.1) and samtools (v1.16) with options view -c and -c -F 260 found 1,815,513,564 primary aligned reads out of 1,838,251,865 total reads. RNA-seq reads used for mapping are from our previous work deposited in NCBI SRA (accessions SRR10947794-SRR10947801). Kallisto (v0.48.0) was used to quantify RNA sequencing data.

The resulting 26 chromosomes of *M. canadense* were annotated using the Funannotate pipeline (v.1.8.7)^54^. First, the chromosomal-level genome assembly was soft-masked using tantan, with repeats and transposable elements soft-masked before gene model prediction using PASA (v.2.4.1) with *M. canadense* RNA-seq reads, *de novo* assembled transcripts and protein homology evidence as input. The gene models and protein homology evidence were then used to train Augustus (v.3.3.3), GeneMark-ES (v.4.61), SNAP(v.2006-07-28) and Glimmerhmm (v.3.0.4) *ab initio* gene predictors and predicted genes passed to Evidence modeller (v.1.1.1) with various weights for integration. tRNAscan-SE (v.2.0.7) was used to predict non-overlapping tRNAs. Transcript evidence from our previous *de novo* transcriptome assembly^4^ using Trinity (v.2.8.5) was leveraged to correct, improve and update the predicted gene models, in addition to refining the 5′- and 3′-untranslated regions in the final step with Funannotate (v.1.8.7). Moreover, the sequence and annotation of DAH paralogous genes that are highlighted in this study were corrected upon amplification of their genomic locus using PCR and validated with their Sanger sequencing results.

### Phylogenomic Analysis

To examine the phylogenetic relationships of *M. canadense* with other related species, we selected 12 additional species for WGD events (Supplementary Tables 6-7). The protein sequences of each species were collapsed using the 90% identity threshold with cd-hit (v4.5.4)^55^. The homologous groups among all 13 species were identified using the OrthoFinder (v2.5.4)^56^ with settings -S diamond -M msa -A muscle -T raxml-ng -I 1.5.

Following OrthoFinder analysis, the amino acid sequences of single-copy orthologous genes from 13 species were aligned using MAFFT (v7.402)^57^ with L-INS-i method and options –maxiterate 1000 –leavegappyregion. Then the protein alignments were reverse-transcribed into their coding sequences using pal2nal (v14)^58^ and trimmed using trimAl (v1.2rev59)^59^. Phylogenetic trees of each single-copy genes were constructed using RAxML (v8.2.11)^59,60^ according to the model of GTRGAMMA with 1000 bootstrap replicates and *A. trichopoda* as the outgroup species (options -# 1000 -o Atri, -x 12345 -p 25258 -f a -T 2). Following gene tree constructions, a consensus species tree was inferred using ASTRAL-III^61^ with 1000 bootstraps. To estimate species divergence times, the Bayesian relaxed molecular clock method in MCMCtree (v4.9j)^20^ was used with the F84 model and the divergence times of *M. canadense*-*S. japonica* (∼35.4-115.2 MYA), *M. cordata*-*P. somniferum* (∼44-82 MYA), *A. coerulea*-*C. chinensis* (∼25.7-79.6 MYA), Ranunculales plants (∼103-118 MYA), and monocots-dicots (∼130-240 MYA) as estimated by a previous study^62^ and TimeTree^63^. CAFE (v5.0)^64^ was used in the Base model and single lambda to infer gene family expansion and contraction in the species tree.

### Phylogenetic Tree Construction and Positive Selection Analysis

An adaptive branch-site REL test for episodic diversification was performed on the gene-tree containing FLS orthologous genes from select angiosperm species as highlighted in Figure 2f and their nucleotide multiple sequence alignment (MSA) using the adaptive branch-site REL test for episodic diversification (aBSREL) method^65^ within the HyPhy program^66^ (v2.3.11). The input MSA contained 14 sequences with 381 sites (codons). One branch of the gene phylogeny leading to the clade containing *DAH*, *DAH-like*, and *DAHy-like* was formally tested for diversifying selection. The aBSREL analysis found evidence of episodic diversifying selection on this node in the phylogeny with significance at p-value of 0.016 after the Holm-Bonferroni correction for multiple hypothesis testing. The intermediate files and results of this analysis, including the nucleotide MSA, GTR based gene-tree, and aBSREL produced adaptive rate class model gene tree are available as Source Data. The corresponding positive selection aBSREL result was mapped onto the protein-based maximum-likelihood phylogenetic tree constructed in Figure 2f. Protein sequences were aligned using the MUSCLE^67^ algorithm in MEGAX^68^. Evolutionary histories were inferred by using the maximum-likelihood method on the basis of the JTT matrix-based model. Bootstrap statistics were calculated using 200 replicates. All phylogenetic analyses were conducted in MEGAX^68^. All alignment files can be found under Source Data.

### Synteny Analysis and Assessment for WGD

To perform macro- and microsynteny analyses between plant genomes explored in this study, BLASTP (v. 2.15.0+) was used to calculate pairwise similarities (*e*-value < 1x10^5^) between CDS DNA of the plant species, and MCScanX^69^ was used with default parameters to identify synthetic gene pairs. MCScanX was further used to visualize the syntenic blocks and assess syntenic depths between plant genomes. Moreover, the DupPipe pipeline from EvoPipes^22^ was used to calculate the *K*_S_ values. The distribution of *K*_S_ values were analyzed using Mixtools^24^ with *k* = 2 and 200 bootstraps at an epsilon value of 0.001 for most analyses. To examine the divergence between plant species, the OrthoPipe pipeline from EvoPipes^22^ was used to calculate the *K*_S_ values between two genomes in comparison. The distribution of these *K*_S_ values were analyzed using Mixtools with *K*_S_ max value increased to 3 and the epsilon value changed to 0.01.

### Protein Expression and Purification

All genes encoding for relevant *Mc*DAH and *Mc*FLS enzymes in this study were codon-optimized for heterologous overexpression in *Escherichia coli*, and purchased as a synthetic gene (Integrated DNA Technologies). Then they were cloned into the pHis8-4b expression vector containing an *N*-terminal 8xHis tag followed by a tobacco etch virus (TEV) cleavage site (Supplementary Table 8). The sequence-verified constructs were transformed into *E. coli* BL21 (DE3) for recombinant protein expression (Supplementary Tables 9-10). A 1 L culture of terrific broth (TB) medium containing 50 µg/mL kanamycin was inoculated with 30 mL of an overnight starter culture and allowed to grow with shaking at 200 rpm at 37 °C to an OD600 of 0.6–0.8. Then protein expression was induced by addition of 0.5 mM isopropyl β-d-1-thiogalactopyranoside (IPTG) followed by cold shock of the medium and subsequent growth with shaking at 200 rpm (18 °C for 18 h).

Cultures were harvested by centrifugation and the resulting cell paste (∼10 g/L) was resuspended in lysis buffer (100 mM Tris pH 8.0, 200 mM NaCl, 20 mM imidazole, 10% (vol/vol) glycerol, 1 mM dithiothreitol) containing 1 mg/mL lysozyme and 1 mM phenylmethylsulfonyl fluoride. Cells were lysed by sonication at 60% amplitude; 30 seconds on, 30 seconds off for 10 min using the flat tip for 1/2″ diameter disruptor horn (Branson Ultrasonics Corporation Sonifier SFX550 Cell). The resulting crude protein lysate was clarified by centrifugation (19,000 × *g*, 45 min) before QIAGEN nickel-nitrilotriacetic acid (Ni-NTA) gravity flow chromatographic purification. After loading the clarified lysate, the Ni-NTA resin was washed with 20 column volumes of lysis buffer and eluted with 1 column volume of elution buffer (100 mM Tris pH 8.0, 200 mM NaCl, 250 mM imidazole, 10% (vol/vol) glycerol, 1 mM dithiothreitol). Then 1 mg of His-tagged TEV protease was added to the eluted protein, followed by dialysis at 4 °C for 16 h in dialysis buffer (25 mM Tris pH 8.0, 200 mM NaCl, 5% (vol/vol) glycerol, 5 mM EDTA, 0.5 mM dithiothreitol). After dialysis, protein solution was passed through Ni-NTA resin to remove uncleaved protein and His-tagged TEV. The recombinant proteins were further purified by gel filtration on an ÄKTA Pure fast protein liquid chromatography (FPLC) system (GE Healthcare Life Sciences). The principal peaks were collected, verified by SDS polyacrylamide gel electrophoresis and dialyzed into a storage buffer (25 mM Tris pH 8.0, 5%(vol/vol) glycerol). Finally, proteins were concentrated to appropriate concentrations using Amicon Ultra-15 Centrifugal Filters (Millipore).

For recombinant expression of 2ODD mutants– *At*F3H D219A, *Ps*CODM D237A, *Cr*D4H H270A, *At*DAO1 H178A, *At*GA2ox-3 H204A, *At*GA3ox-1 H234A, *At*JOX1 H275A, *At*DMR6 H214A, *Zm*BX6 H245A– codon optimized gene fragments were purchased as synthetic genes then cloned into the pHis8-4b vector. For *At*F3H, the D219G mutant was also generated and cloned into the pHis8-4b vector. For recombinant expression of *Hn*H6H D219A, codon optimized gene fragment was cloned into the plasmid pBA0221-0141^70^ containing a *C*-terminal 8xHis following a TEV cleavage site (Supplementary Table 8). After obtaining each of these plasmid constructs, the same protocol as above was initially carried out for all 2ODD mutants.

However, most 2ODD mutants– *At*F3H D219A, *Ps*CODM D237A, *Cr*D4H H270A, *At*DAO1 H178A, *At*GA2ox-3 H204A, *At*GA3ox-1 H234A, *At*JOX1 H275A, *At*DMR6 H214A, *Zm*BX6 H245A– were either found in the insoluble fraction during the protein purification process or exhibit no detectable enzyme activity. For BL21 strains expressing *At*F3H D219G and *Hn*H6H D219A, 12 L culture of TB medium was used to increase the yield of these relatively unstable mutants.

For recombinant expression of *Mc*DAH alanine mutants, site-directed mutagenesis was performed according to the protocol described in the QuickChange II Site-Directed Mutagenesis Kit (Agilent Technologies) using plasmid pHis8-4b::*Mc*DAH as the template and the primer sequences in Supplementary Table 9. The resulting mutant plasmid constructs were verified by sequencing. Recombinant mutant protein production and purification were carried out following the same procedure as described above.

### In vitro Enzyme Assays

Each enzyme assay for *Mc*DAH-WT and relevant mutants reported in this study with (−)-dechloroacutumine was carried out in 50 mM Tris buffer, pH = 8.0, on a 20 µL reaction containing the following components: enzyme (5 µM), (−)-dechloroacutumine (1 mM), 2OG (500 µM), NaCl (1 mM), sodium ascorbate (5 mM), and (NH_4_)_2_Fe(So_4_)_2_ (2 mM). In a typical assay, the components were added in the following order: (1) Tris, (2) NaCl, (3) enzyme, (4) sodium ascorbate, (5) 2OG, (6) (−)-dechloroacutumine, and (7) (NH_4_)_2_Fe(So_4_)_2_. The samples were incubated under aerobic conditions at room temperature for 20 mins. The assays were quenched by adding methanol to 50% final concentration and centrifugation to spin down enzymes and debris. The supernatants were filtered using 0.2 µm filter vials (Thomson Instrument Company) prior to injection (2 µL) into the LC-HRAM-MS system for analysis. Relevant enzyme assays for activity against dihydroflavonols were performed similarly, but without the addition of (2) and substitution of (6) with 1 mM dihydrokaempferol or 1 mM dihydroquercetin, then quenched by adding acetonitrile with 0.1% formic acid to 50% final concentration. All 2ODD mutant assays were performed similarly using NaCl or NaN_3_ as the salt in the reaction for chlorination or azidation reactions, respectively. For *At*F3H D219G assays, 2*S*-naringenin was used as the substrate, whereas hyoscyamine was used as the substrate for *Hn*H6H D219A assays.

### LC-HRAM-MS Analysis

LC was conducted on a Vanquish Flex Binary UHPLC system (Thermo Fisher Scientific) using water with 0.1% formic acid as solvent A and acetonitrile with 0.1% formic acid as solvent B. Reverse phase separation of analytes was performed on a Kinetex C18 column, 150 × 3 mm^2^, 2.6 µm particle size (Phenomenex). The column oven was held at 35 °C. Most injections were eluted with 5% B for 0.5 min, a gradient of 5–95% B for 14.5 min, 95% B for 2 min, and 5% B for 3.0 min, with a flow rate of 0.5 mL/min. Most MS analyses were performed on a high-resolution Orbitrap Exploris 120 benchtop mass spectrometer (Thermo Fisher Scientific) operated in positive ionization mode with full scan range of 100–450 *m*/*z* and top four data-dependent MS/MS scans. The orbitrap resolution is 120,000 with RF lens of 70% and static spray voltage of 3500 V. For detecting dihydroflavonols and flavonols from relevant enzyme assays, injections were eluted with 5% B for 0.5 min, a gradient of 5–95% B for 14.5 min, 95% B for 2 min, and 5% B for 3.0 min, with a flow rate of 0.5 mL/min. Moreover, MS analysis was operated in negative ionization mode with full scan range of 100-450 *m/z* and top four data-dependent MS/MS scans using static spray voltage of 2500 V. For detecting products from *At*F3H D219G assay, injections were eluted with 20% B for 0.5 min, a gradient of 20–50% B for 6.5 min, 50-80% B for 0.5 min, 80% B for 0.5 min, and 20% B for 2 min with a flow rate of 0.5 mL/min using a Kinetex C18 column, 50 × 3 mm^2^, 2.6 µm particle size (Phenomenex). MS analysis was operated in negative ionization mode with full scan range of 150-400 *m/z* and top four data-dependent MS/MS scans using static spray voltage of 2500 V. For detecting products from *Hn*H6H D219A assay, injections were eluted with 5% B for 0.5 min, a gradient of 5-30% B for 6.5 min, 30-75% B for 0.5 min, 75% B for 0.5 min, and 5% B for 2 min with a flow rate of 0.5 mL/min using the Kinetex C18 column, 50 × 3 mm^2^, 2.6 µm particle size. MS analysis was operated in positive ionization mode with full scan range of 150-450 *m/z* and top four data-dependent MS/MS scans using static spray voltage of 3500 V. Raw LC-MS data were collected and analyzed using Chromeleon 7.2.10 ES, TSQ Tune 3.1.279.9, and XCalibur 4.5 (Thermo Fisher Scientific).

### Protein Structure and Preparation

AlphaFold2 was used to generate folded structures for DAH and FLS using the default parameters of the AlphaFold Singularity container for version 2.0.0^28^. All subsequent structural models discussed in this study are derived from AlphaFold2 folded structures and we demonstrate that AlphaFold3 folded structures have no significant structural differences (Supplementary Fig. 34). Protonation states were assigned using the H++ webserver with a pH of 7.0 and an internal dielectric of 10 while retaining all other system defaults (Supplementary Tables 11-12)^71–73^. Core active site residues were reviewed and assigned such that histidines were neutral and the metal-coordinating carboxylate in FLS was negatively charged. Molecular docking runs were performed for enzyme-substrate pairs FLS and dihydrokaempferol, and DAH and dechloroacutumine using AutoDock Vina 1.1.2 (Supplementary Fig. 19)^74,75^. We found that dihydrokaempferol had a binding energy of -8.2 kcal/mol, and dechloroacutumine had a binding energy of -7.5 kcal/mol. The lowest energy substrate conformations of dihydrokaempferol and dechloroacutumine were selected as the initial substrate binding poses for FLS and DAH, respectively. Structures of the initial binding poses are provided in the Source Data as a .zip file. Based on the dihydrokaempferol and dechloroacutumine docked complexes, we generated models for both FLS and DAH with either 2OG or succinate and oxo bound. We also generated DAH models with chlorine and the oxo in both the equatorial and axial positions. The ligands 2OG, succinate, Fe, Cl, and oxo were modeled manually into their corresponding structures. The resulting DAH and FLS holoenzymes had an atom count and a net charge of 5536 and -8 for FLS with 2OG, 5535 and -8 for FLS with succinate, 5697 and -9 for DAH with 2OG, 5696 and -9 for DAH with succinate and an equatorial-oxo, and 5696 and -9 for DAH with succinate and an axial-oxo.

Utilizing the tleap utility in the AMBER software suite, topology and coordinate files for the final structures were generated for the AMBER ff14SB force field^76^. The ligands and substrates were parameterized using the generalized AMBER force field (GAFF) and restrained electrostatic potential (RESP) charges^77^. These charges were calculated with Gaussian16 at the HF/6-31G* level of theory^78^. The core active site parameters were obtained using AMBER’s Metal Center Parameter Builder (MCPB)^79^. The MCPB.py v3.0 script was employed to compute charges with the ChgModB method at the UB3LYP/LACVP*^80,81^ level of theory for iron-coordinated residues. The Seminario method^82^ was employed to derive additional force field parameters, with details available in the Supplementary Figures 20 and 22. Subsequently, each protein system was solvated with a 15Å TIP3P water box with periodic boundary conditions and neutralized with Na+ counterions^83^. The resulting final atom counts for the various systems were: 64,767 for FLS with 2OG, 64,766 for FLS with succinate, 72,247 for DAH with 2OG, 72,246 for DAH with succinate and an equatorial-oxo, and 72,246 for DAH with succinate and an axial-oxo. The topology and inpcrd files for all MD simulations are provided in the Source Data as a .zip file.

### Classical Molecular Dynamics Simulations and Analysis

We performed molecular dynamics (MD) simulations for DAH and FLS using AMBER18 and the GPU-accelerated particle mesh Ewald (PME)^84^ molecular dynamics (PMEMD) code^85,86^. The equilibration protocol involved three steps: i) 1,000 steps of hydrogen atom minimization, 2,000 steps of sidechain minimization with a fixed backbone, and 2,000 steps of protein minimization with the core active site restrained; ii) controlled NVT heating from 0 to 300 K over 10 ps using the Langevin thermostat and a collision frequency of 5.0 ps-1; and iii) a 1 ns NPT simulation with the Berendsen barostat and a 2 ps relaxation time. Following equilibration, we collected 250 ns of production dynamics for each enzyme-substrate complex, employing 2 fs time steps, the SHAKE algorithm to fix hydrogen-heavy atom distances^87^, and electrostatics were treated with the PME method with a real-space cutoff of 10 Å^84^. Restrained MD simulations were also carried out for 250ns production runs for each system with flat-bottom harmonic restraints applied to the oxo, iron, and HAT target angle and the iron-HAT target distance to increase sampling of halogenase and hydroxylase expected angles (Supplementary Table 13). For all systems, at least four 250 ns replicates were performed.

We also performed 250 ns production runs for restrained MD simulations in each system, using harmonic restraints to enhance the sampling of target angles and distances typically observed for halogenases and hydroxylases (Supplementary Tables 14-15)^30,36,88,89^. Such experimentally informed MD simulations have been used in guiding similar computational studies of non-heme iron halogenases and hydroxylases^30,35,90^.

The harmonic restraints are based on experimental hyperfine sublevel correlation (HYSCORE) spectroscopy data, which extracts relative spatial information (e.g., distances and angles) about the metallocofactor and nearby ^2^H labels on the substrate^36,91,92^. To select representative frames for analysis, we clustered trajectories with the DBSCAN algorithm in CPPTRAJ^93^, using the substrates as masks and following a previously described method (Supplementary Tables 14-15)^90^. Following clustering, the centroid of the largest cluster was used for QM/MM optimization (Supplementary Figures 21 and 23). We used MMPBSA.py in AMBER18^90,94,95^ for interaction analysis of the selected snapshots, employing the generalized Born (GB)^35,90^ approximation and the OBC1 model. This method computes the contributions to pairwise residue interactions, electrostatics, and van der Waals’ binding^96,97^. We used 1000 snapshots from the end of the simulations, spaced 50 ps apart for GBSA analysis. Classical geometric hydrogen bonding analysis was conducted with the CPPTRAJ utility, applying a hydrogen bonding distance cutoff of 3.2 Å based on previously quantified hydrogen bonding strengths^97^. or restrained simulations, we used harmonic restraints of 100 kcal/(mol · Å^2^) as previously described^30,35^.

### Quantum Mechanical / Molecular Mechanical Simulations

We ran QM/MM geometry optimizations starting from the centroids of the largest clusters of the MD production runs for all FLS and DAH isomers. Spherical droplets with a radius of 35-42 Å, centered around the protein’s center of mass, were next created using PyMOL^98^ from the periodic box extracted from the MD simulations. Counterions were added using tleap to neutralize the droplets. The QM/MM simulations were conducted using a developer version of TeraChem v1.9^99–101^ for the QM component and AMBER18 for the MM component^85,86^. In the QM/MM simulations, no atoms were held fixed. A weak restraining spherical cap (force constant of 1.5 kcal/mol.Å^2^) was applied to maintain the MM water droplet’s shape and prevent significant movement during optimization. Unrestricted density functional theory (DFT) with the range-separated hybrid ωPBEh (ω = 0.2 Bohr^−1^)^102^ and a basis set consisting of the LANL2DZ effective core potential^103^ was used for Fe while 6-31G* was used for all other atoms in the QM region^80,81^. FLS and DAH were modeled in the high-spin state (i.e., quintet multiplicity, 2*S*+1 = 5). The QM regions of DAH and FLS included their respective substrates, the Fe center, and all ligands and residues coordinating Fe (Supplementary Tables 16-19). The only residue from the second coordination sphere included in the QM region was Lys205 in DAH and its hydroxylase corollary Lys209 in FLS, which is catalytically essential. The QM regions incorporated the following atom counts and total charges: 116 and 0 for FLS with 2OG, 115 and 0 for FLS with succinate, 123 and 0 for DAH with 2OG, 122 and 0 for DAH with succinate and an equatorial-oxo, and 122 and 0 for DAH with succinate and an axial-oxo.

## Data availability

The raw data of genome sequencing of *M. canadense* has been deposited to the NCBI SRA under BioProject No. PRJCA######. The genome assembly and annotation of *M. canadense* assembled are available at figshare platform (XXXXXX). Relevant LC-HRAM-MS raw data files generated in this study have been deposited in Zenodo [https://doi.org/#########]. AlphaFold2.0 structural models along with QM-MM optimized structures are provided in the Source data file. Relevant genome assembly statistics, phylogenomic analyses, and comparative genomic resources are found in the Source data file. Additional data are also available from the corresponding author upon request. Source data are provided with this paper.

## Acknowledgements

We would like to thank the staff at PhaseGenomics and Mount Sinai Genomics Technology Facility for their assistance with *M. canadense* genome sequencing and assembly. We would like to thank the Bollinger/Krebs group at Pennsylvania State University for providing the pBA0221-0141-*Hn*H6H expression plasmid. This work was supported by the W. M. Keck Foundation (J.-K.W.), the Family Larsson-Rosenquist Foundation (J.-K.W.), the Beckman Young Investigator Program (J.-K.W.), the Schooner Foundation (J.-K.W.), the National Institute of General Medical Sciences of the National Institutes of Health under award number R35GM152027 (H.J.K., D.W.K), and a National Science Foundation Graduate Research Fellowship under Grant #1745302 (D.W.K.).

## Author Contributions

C.Y.K. and J.-K.W. conceived and designed the research. C.Y.K. led the sequencing, assembly, and analysis of the *M. canadense* genome. C.Y.K. performed all biochemical assays and evolutionary analyses. D.W.K. and H.J.K. conducted and analyzed the MD simulations and QM/MM calculations. A.J.M., M.A.G., and J.S.Y. helped with the cloning, protein purification, and biochemical assays of plant 2ODD mutants. E.N.N. helped with the cloning, protein purification, and biochemical assays of FLS-DAH mutants. C.Y.K., D.W.K., H.J.K., and J.-K.W. interpreted the results and wrote the manuscript. All authors reviewed the manuscript.

## Supplementary information

### Supplementary Tables

**Supplementary Table 1.** Summary of sequencing reads used for assembling and polishing the *Menispermum canadense* genome.

|  | Data Type | Raw Reads | Raw Bases (G) | Clean Reads | Clean Bases (G) | Q20 (%) | Q30 (%) | GC content (%) |
| --- | --- | --- | --- | --- | --- | --- | --- | --- |
| **Long Read (Primary Assembly)** | PacBio Sequel II (HIFI CCS) | 3.079435 M | 48.571952 | 3.079435 M | 48.571952 | 98.06 | 95.41 | 35.42 |
| **Short Read (QC)** | Illumina NovaSeq (Paired End 150bp : 150bp) | 1.801201 G | 270.180090 | 1.785581 G | 266.380866 | 97.64 | 92.95 | 35.12 |
| **Hi-C (Scaffolding)** | Illumina NovaSeq (Paired End 150bp : 150bp) | 1.084218 G | 162.632754 | 1.034976 G | 154.857483 | 95.81 | 90.30 | 35.86 |

**Supplementary Table 2.** Length and scaffolds for chromosome-level assembly of the *Menispermum canadense* genome.

| **Pseudo-chromosome** | **# Contigs** | **Size (bp)** |
| --- | --- | --- |
| Chr1 | 1 | 45,646,067 |
| Chr2 | 1 | 44,085,362 |
| Chr3 | 1 | 42,833,333 |
| Chr4 | 1 | 42,297,172 |
| Chr5 | 1 | 42,242,317 |
| Chr6 | 1 | 40,690,050 |
| Chr7 | 1 | 37,974,986 |
| Chr8 | 1 | 36,980,421 |
| Chr9 | 1 | 36,537,451 |
| Chr10 | 1 | 36,006,012 |
| Chr11 | 1 | 35,609,015 |
| Chr12 | 1 | 34,818,747 |
| Chr13 | 1 | 34,191,516 |
| Chr14 | 1 | 33,879,814 |
| Chr15 | 1 | 33,347,802 |
| Chr16 | 1 | 32,511,916 |
| Chr17 | 1 | 31,451,445 |
| Chr18 | 1 | 31,054,424 |
| Chr19 | 1 | 30,121,998 |
| Chr20 | 1 | 29,630,568 |
| Chr21 | 1 | 29,598,887 |
| Chr22 | 1 | 29,497,318 |
| Chr23 | 1 | 27,578,300 |
| Chr24 | 1 | 24,848,486 |
| Chr25 | 1 | 24,327,722 |
| Chr26 | 1 | 22,787,794 |
| unmapped | 1249 | 79,359,561 |

**Supplementary Table 3.** Sequence reads mapped onto the genome using bwa-mem (v0.7.17-r1188)^1^ (Illumina) and minimap (v2.24-r1122)^2^ (Transcripts).

| Property | Number |
| --- | --- |
| Primary aligned reads (Illumina) | 1,800,058,803 |
| Total number of reads (Illumina) | 1,820,675,006 |
| Primary aligned reads (Transcripts) | 733,893 |
| Total number of reads (Transcripts) | 734,023 |

**Supplementary Table 4.** Genome annotation statistics.

| Property | Number |
| --- | --- |
| Total number of protein-coding genes | 65,843 |
| Total number of exons | 284,345 |
| Average exon length (bp) | 173.15 |
| Average protein length (aa) | 290.9 |
| Average gene length (bp) | 3088.44 |

**Supplementary Table 5.** Statistics from genome quality assessment by Merqury^3^.

| Property | Number |
| --- | --- |
| *k*-mer used | 20 |
| Solid *k*-mers in assembly | 403,045,961 |
| Total solid *k*-mers in the read set | 479,987,586 |
| Unique *k*-mers in assembly | 20,962 |
| *k*-mers in both assembly and read set | 969,874,501 |
| Assembly consensus quality value (QV) | 59.6631 |
| Error rate | 1.08067 x 10<sup>-6</sup> |

**Supplementary Table 6.** Source of plant genomes for phylogenomic and comparative genomics analyses.

| **Plant Taxon** | **Short/Common name** | **Reference** |
| --- | --- | --- |
| *Aquilegia coerulea* | Rocky Mountain Columbine | NCBI (GCA002738505) |
| *Arabidopsis thaliana* | *Arabidopsis* | Araport11 |
| *Amborella trichopoda* | *Amborella* | NCBI (GCA000471905) |
| *Coptis chinensis* | Chinese Goldthread | NCBI (GCA015680905) |
| *Cinnamomum kanehirae* | Stout Camphor Tree | NCBI (GCA003546025) |
| *Menispermum canadense* | Common Moonseed | This manuscript |
| *Macleaya cordata* | Plume Poppy | NCBI (GCA002174775) |
| *Nelumbo nucifera* | Sacred Lotus | nelumbo.cngb.org |
| *Oryza sativa* | Rice | rice.uga.edu |
| *Papaver somniferum* | Opium Poppy | NCBI (GCA003573695) |
| *Stephania japonica* | Snake Vine | NCBI (GCA039657345) |
| *Theobroma cacao* | Cacao | cocoa-genome-hub.southgreen.fr |
| *Vitis vinifera* | Grape | genoscope.cns.fr/externe/GenomeBrowser/Vitis/ |

**Supplementary Table 7.** Prediction of orthologous groups by OrthoFinder^4^.

| Plant Taxon | Total curated genes | Number of genes in orthogroups | Number of unassigned genes | Number of orthogroups containing species | Percentage of orthogroups containing species | Number of species-specific orthogroups | Number of genes in species-specific orthogroups |
| --- | --- | --- | --- | --- | --- | --- | --- |
| *Aquilegia coerulea* | 24102 | 218897 | 2205 | 12719 | 44.7% | 303 | 977 |
| *Arabidopsis thaliana* | 28258 | 26225 | 2033 | 12764 | 44.9% | 976 | 4340 |
| *Amborella trichopoda* | 25477 | 21296 | 4181 | 12968 | 45.6% | 795 | 3852 |
| *Coptis chinensis* | 34241 | 32667 | 1574 | 13947 | 49.0% | 1148 | 5206 |
| *Cinnamomum kanehirae* | 24115 | 22763 | 1352 | 12292 | 43.2% | 326 | 1472 |
| *Menispermum canadense* | 55164 | 47394 | 7770 | 15361 | 54.0% | 1752 | 15244 |
| *Macleaya cordata* | 20525 | 19724 | 801 | 12504 | 43.9% | 105 | 332 |
| *Nelumbo nucifera* | 28619 | 24084 | 4535 | 13065 | 45.9% | 648 | 1991 |
| *Oryza sativa* | 51870 | 43687 | 8183 | 14141 | 49.7% | 2525 | 18052 |
| *Papaver somniferum* | 37269 | 32491 | 4778 | 14071 | 49.5% | 1786 | 6893 |
| *Stephania japonica* | 25157 | 22357 | 2800 | 12984 | 45.6% | 599 | 3600 |
| *Theobroma cacao* | 21590 | 21132 | 458 | 12620 | 44.4% | 247 | 1022 |
| *Vitis vinifera* | 23743 | 20878 | 2865 | 12575 | 44.2% | 380 | 1140 |

**Supplementary Table 8.** Strains and plasmids used in this study.

| Strain | Description | Source |
| --- | --- | --- |
| BL21 (DE3) | *fhuA2 [lon] ompT gal (λ DE3) [dcm] ΔhsdS* | New England BioLabs |

| Plasmid | Description | Source |
| --- | --- | --- |
| pHis8-4b-His<sub>8</sub>-McDAH | His<sub>8</sub>-McDAH (T7), *lacI*, *lacO*, Kan<sup>R</sup>, F1-ori | This study |
| pHis8-4b-His<sub>8</sub>-McFLS | His<sub>8</sub>-McFLS (T7), *lacI*, *lacO*, Kan<sup>R</sup>, F1-ori | This study |
| pHis8-4b-His<sub>8</sub>-McFLS D230G | His<sub>8</sub>-McFLS-D230G (T7), *lacI*, *lacO*, Kan<sup>R</sup>, F1-ori | This study |
| pHis8-4b-His<sub>8</sub>-McFLS<sub>(6)</sub> | His<sub>8</sub>-McFLS<sub>(6)</sub> (T7), *lacI*, *lacO*, Kan<sup>R</sup>, F1-ori | This study |
| pHis8-4b-His<sub>8</sub>-McFLS<sub>(14)</sub> | His<sub>8</sub>-McFLS<sub>(14)</sub> (T7), *lacI*, *lacO*, Kan<sup>R</sup>, F1-ori | This study |
| pHis8-4b-His<sub>8</sub>-McFLS<sub>(40)</sub> | His<sub>8</sub>-McFLS<sub>(40)</sub> (T7), *lacI*, *lacO*, Kan<sup>R</sup>, F1-ori | This study |
| pHis8-4b-His<sub>8</sub>-McDAH<sub>(14)</sub> | His<sub>8</sub>-McDAH<sub>(14)</sub> (T7), *lacI*, *lacO*, Kan<sup>R</sup>, F1-ori | This study |
| pHis8-4b-His<sub>8</sub>-McDAH<sub>(40)</sub> | His<sub>8</sub>-McDAH<sub>(40)</sub> (T7), *lacI*, *lacO*, Kan<sup>R</sup>, F1-ori | This study |
| pHis8-4b-His<sub>8</sub>-AtF3H D219G | His<sub>8</sub>-AtF3H D219G (T7), *lacI*, *lacO*, Kan<sup>R</sup>, F1-ori | This study |
| pBA0221-0141-HnH6H<br>D219A-His<sub>8</sub> | HnH6H D219A-His<sub>8</sub> (T7), *lacI*, *lacO*, Kan<sup>R</sup>, F1-ori,<br>pBR322-ori, Tet<sup>R</sup> | This study |
| pHis8-4b-His<sub>8</sub>-McDAH T231A | His<sub>8</sub>-McDAH T231A (T7), *lacI*, *lacO*, Kan<sup>R</sup>, F1-ori | This study |
| pHis8-4b-His<sub>8</sub>-McDAH N262A | His<sub>8</sub>-McDAH N262A (T7), *lacI*, *lacO*, Kan<sup>R</sup>, F1-ori | This study |
| pHis8-4b-His<sub>8</sub>-McDAH K205A | His<sub>8</sub>-McDAH K205A (T7), *lacI*, *lacO*, Kan<sup>R</sup>, F1-ori | This study |
| pHis8-4b-His<sub>8</sub>-McDAH K332A | His<sub>8</sub>-McDAH K332A (T7), *lacI*, *lacO*, Kan<sup>R</sup>, F1-ori | This study |
| pHis8-4b-His<sub>8</sub>-McDAH L115A | His<sub>8</sub>-McDAH L115A (T7), *lacI*, *lacO*, Kan<sup>R</sup>, F1-ori | This study |
| pHis8-4b-His<sub>8</sub>-McDAH Y227A | His<sub>8</sub>-McDAH Y227A (T7), *lacI*, *lacO*, Kan<sup>R</sup>, F1-ori | This study |
| pHis8-4b-His<sub>8</sub>-McDAH L333A | His<sub>8</sub>-McDAH L333A (T7), *lacI*, *lacO*, Kan<sup>R</sup>, F1-ori | This study |
| pHis8-4b-His<sub>8</sub>-McDAH Y126A | His<sub>8</sub>-McDAH Y126A (T7), *lacI*, *lacO*, Kan<sup>R</sup>, F1-ori | This study |
| pHis8-4b-His<sub>8</sub>-McDAH V222A | His<sub>8</sub>-McDAH V222A (T7), *lacI*, *lacO*, Kan<sup>R</sup>, F1-ori | This study |
| pHis8-4b-His<sub>8</sub>-McDAH L135A | His<sub>8</sub>-McDAH L135A (T7), *lacI*, *lacO*, Kan<sup>R</sup>, F1-ori | This study |
| pHis8-4b-His<sub>8</sub>-McDAH R121A | His<sub>8</sub>-McDAH R121A (T7), *lacI*, *lacO*, Kan<sup>R</sup>, F1-ori | This study |
| pHis8-4b-His<sub>8</sub>-McDAH V221A | His<sub>8</sub>-McDAH V221A (T7), *lacI*, *lacO*, Kan<sup>R</sup>, F1-ori | This study |
| pHis8-4b-His<sub>8</sub>-McDAH F137A | His<sub>8</sub>-McDAH F137A (T7), *lacI*, *lacO*, Kan<sup>R</sup>, F1-ori | This study |
| pHis8-4b-His<sub>8</sub>-McDAH F296A | His<sub>8</sub>-McDAH F296A (T7), *lacI*, *lacO*, Kan<sup>R</sup>, F1-ori | This study |
| pHis8-4b-His<sub>8</sub>-McDAH R108A | His<sub>8</sub>-McDAH R108A (T7), *lacI*, *lacO*, Kan<sup>R</sup>, F1-ori | This study |
| pHis8-4b-His<sub>8</sub>-McDAH<br>R108A/R121A | His<sub>8</sub>-McDAH R108A/R121A (T7), *lacI*, *lacO*, Kan<sup>R</sup>,<br>F1-ori | This study |

**Supplementary Table 9.**
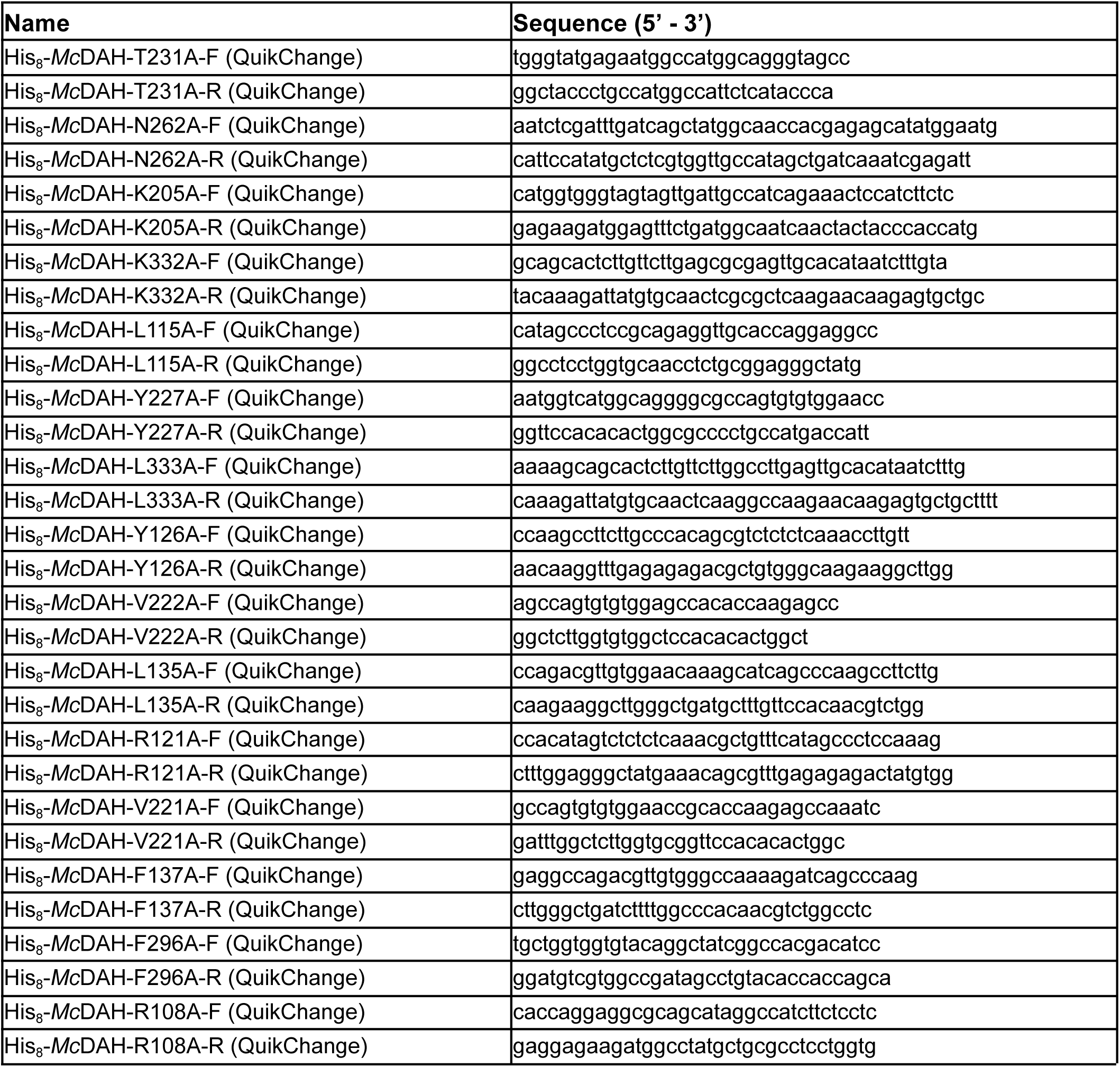
Oligonucleotide sequences reported in this study.

**Supplementary Table 10.**
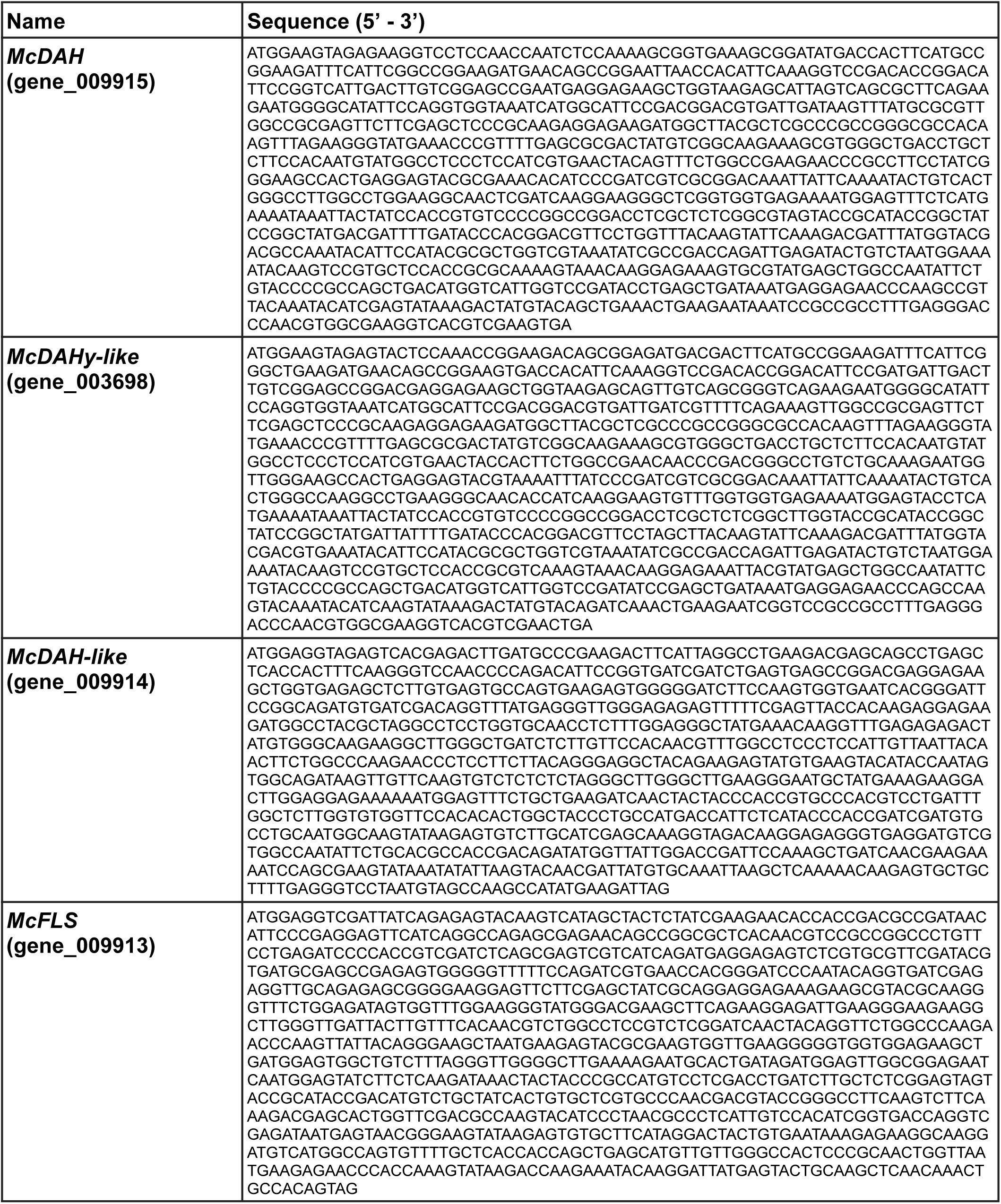
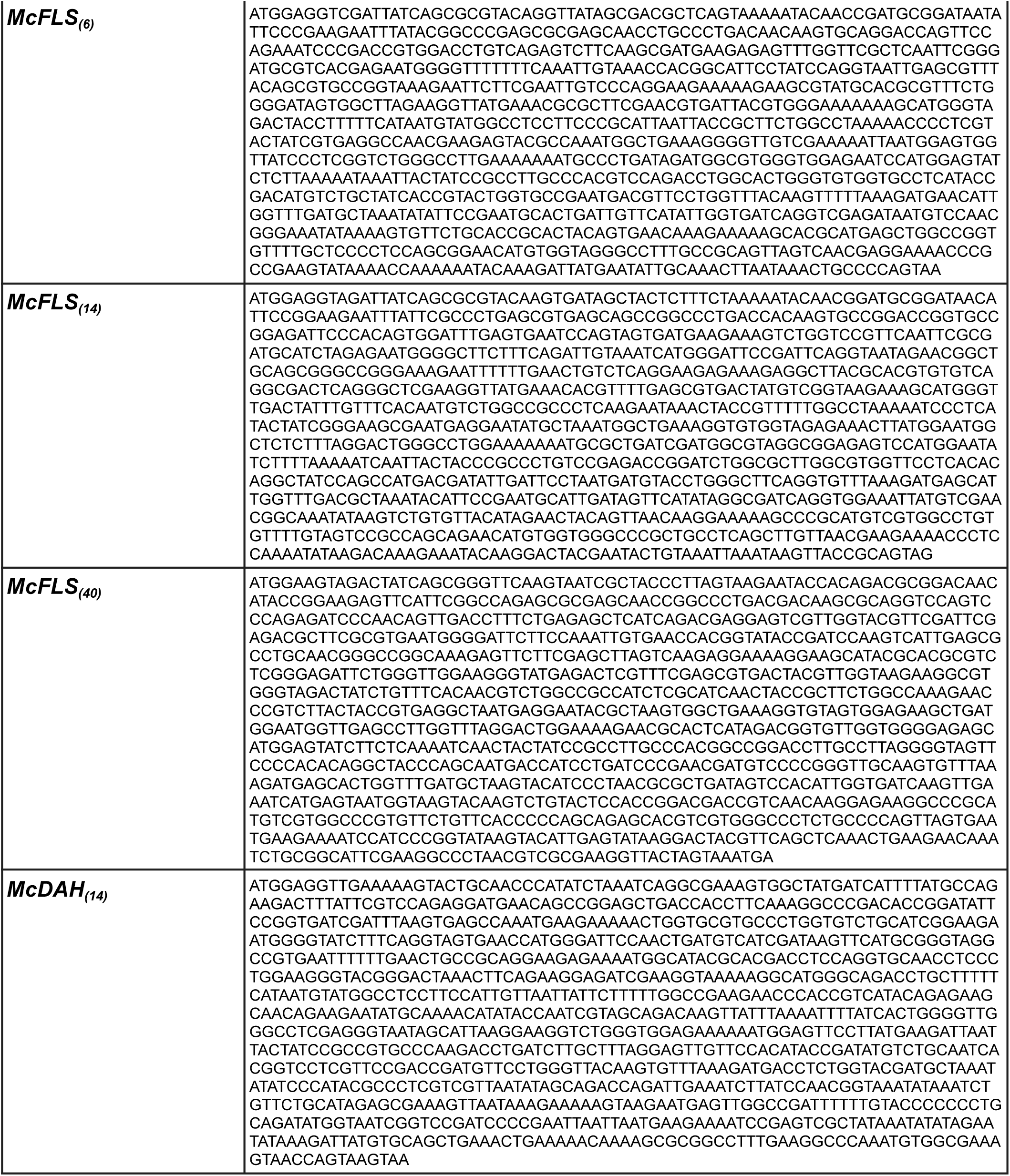
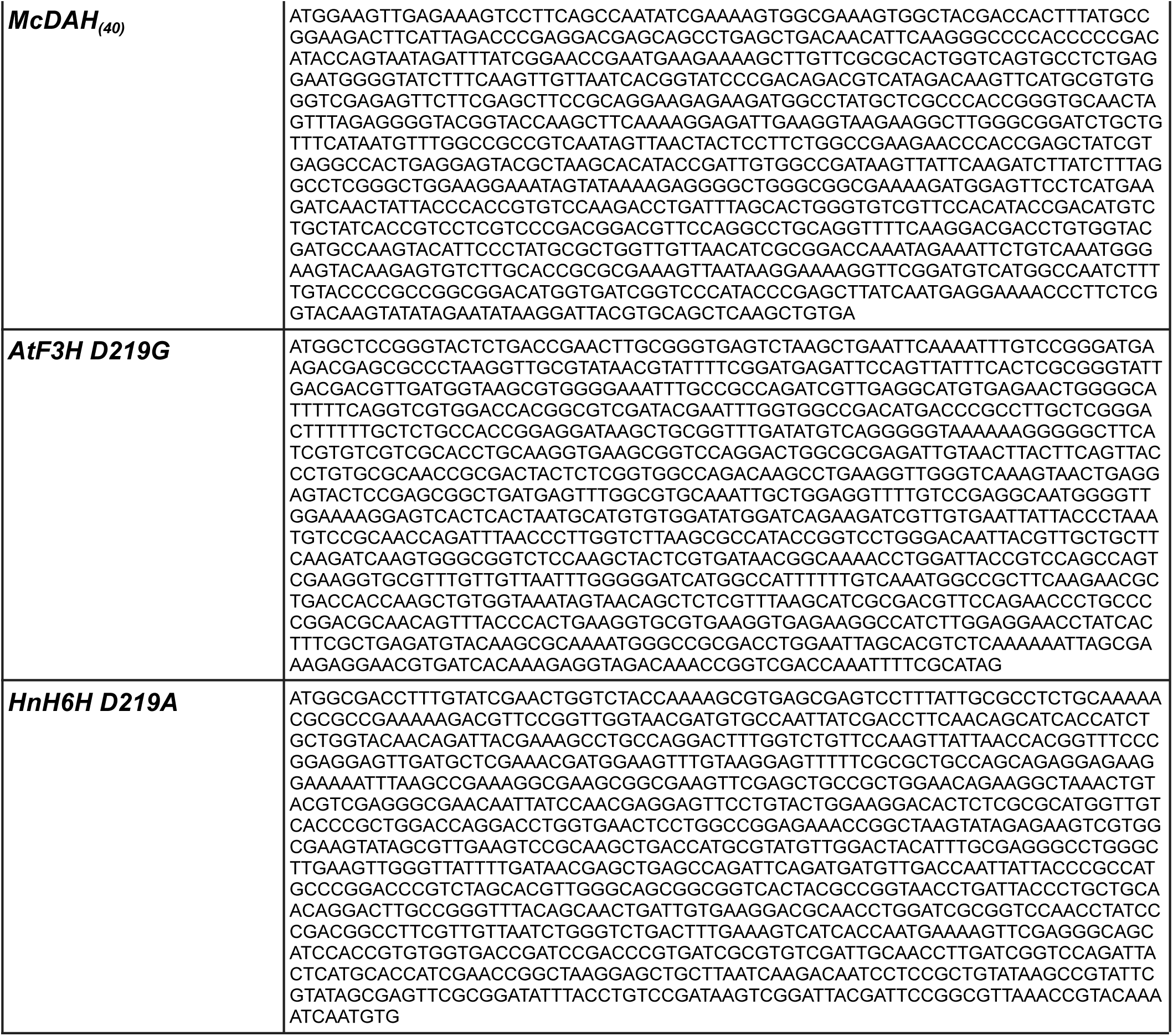
Gene sequences reported in this study.

**Supplementary Table 11.** FLS protonation states and charges (*q*) with neutral charges left blank.

| Residue | *q* | Residue | *q* | Residue | *q* | Residue | *q* | Residue | *q* | Residue | *q* | Residue | *q* |
| --- | --- | --- | --- | --- | --- | --- | --- | --- | --- | --- | --- | --- | --- |
| MET1 |  | THR50 |  | PHE99 |  | SER148 |  | ASP197 | -1 | GLN246 |  | MET295 |  |
| GLU2 | -1 | VAL51 |  | PHE100 |  | ARG149 | +1 | GLY198 |  | VAL247 |  | SER296 |  |
| VAL3 |  | ASP52 | -1 | GLU101 | -1 | ILE150 |  | VAL199 |  | PHE248 |  | TRP297 |  |
| ASP4 | -1 | LEU53 |  | LEU102 |  | ASN151 |  | GLY200 |  | LYS249 | +1 | PRO298 |  |
| TYR5 |  | SER54 |  | SER103 |  | TYR152 |  | GLY201 |  | ASP250 | -1 | VAL299 |  |
| GLN6 |  | GLU55 | -1 | GLN104 |  | ARG153 | +1 | GLU202 | -1 | GLU251 | -1 | PHE300 |  |
| ARG7 | +1 | SER56 |  | GLU105 | -1 | PHE154 |  | SER203 |  | HIP252 | +1 | CYS301 |  |
| VAL8 |  | SER57 |  | GLU106 | -1 | TRP155 |  | MET204 |  | TRP253 |  | SER302 |  |
| GLN9 |  | SER58 |  | LYS107 | +1 | PRO156 |  | GLU205 | -1 | PHE254 |  | PRO303 |  |
| VAL10 |  | ASP59 | -1 | GLU108 | -1 | LYS157 | +1 | TYR206 |  | ASP255 | -1 | PRO304 |  |
| ILE11 |  | GLU60 | -1 | ALA109 |  | ASN158 |  | LEU207 |  | ALA256 |  | ALA305 |  |
| ALA12 |  | GLU61 | -1 | TYR110 |  | PRO159 |  | LEU208 |  | LYS257 | +1 | GLU306 | -1 |
| THR13 |  | SER62 |  | ALA111 |  | SER160 |  | LYS209 | +1 | TYR258 |  | HIE307 |  |
| LEU14 |  | LEU63 |  | ARG112 | +1 | TYR161 |  | ILE210 |  | ILE259 |  | VAL308 |  |
| SER15 |  | VAL64 |  | VAL113 |  | TYR162 |  | ASN211 |  | PRO260 |  | VAL309 |  |
| LYS16 | +1 | ARG65 | +1 | SER114 |  | ARG163 | +1 | TYR212 |  | ASN261 |  | GLY310 |  |
| ASN17 |  | SER66 |  | GLY115 |  | GLU164 | -1 | TYR213 |  | ALA262 |  | PRO311 |  |
| THR18 |  | ILE67 |  | ASP116 | -1 | ALA165 |  | PRO214 |  | LEU263 |  | LEU312 |  |
| THR19 |  | ARG68 | +1 | SER117 |  | ASN166 |  | PRO215 |  | ILE264 |  | PRO313 |  |
| ASP20 | -1 | ASP69 | -1 | GLY118 |  | GLU167 | -1 | CYS216 |  | VAL265 |  | GLN314 |  |
| ALA21 |  | ALA70 |  | LEU119 |  | GLU168 | -1 | PRO217 |  | HIE266 |  | LEU315 |  |
| ASP22 | -1 | SER71 |  | GLU120 | -1 | TYR169 |  | ARG218 | +1 | ILE267 |  | VAL316 |  |
| ASN23 |  | ARG72 | +1 | GLY121 |  | ALA170 |  | PRO219 |  | GLY268 |  | ASN317 |  |
| ILE24 |  | GLU73 | -1 | TYR122 |  | LYS171 | +1 | ASP220 | -1 | ASP269 | -1 | GLU318 | -1 |
| PRO25 |  | TRP74 |  | GLY123 |  | TRP172 |  | LEU221 |  | GLN270 |  | GLU319 | -1 |
| GLU26 | -1 | GLY75 |  | THR124 |  | LEU173 |  | ALA222 |  | VAL271 |  | ASN320 |  |
| GLU27 | -1 | PHE76 |  | LYS125 | +1 | LYS174 | +1 | LEU223 |  | GLU272 | -1 | PRO321 |  |
| PHE28 |  | PHE77 |  | LEU126 |  | GLY175 |  | GLY224 |  | ILE273 |  | PRO322 |  |
| ILE29 |  | GLN78 |  | GLN127 |  | VAL176 |  | VAL225 |  | MET274 |  | LYS323 | +1 |
| ARG30 | +1 | ILE79 |  | LYS128 | +1 | VAL177 |  | VAL226 |  | SER275 |  | TYR324 |  |
| PRO31 |  | VAL80 |  | GLU129 | -1 | GLU178 | -1 | PRO227 |  | ASN276 |  | LYS325 | +1 |
| GLU32 | -1 | ASN81 |  | ILE130 |  | LYS179 | +1 | HID228 |  | GLY277 |  | THR326 |  |
| ARG33 | +1 | HIE82 |  | GLU131 | -1 | LEU180 |  | THR229 |  | LYS278 | +1 | LYS327 | +1 |
| GLU34 | -1 | GLY83 |  | GLY132 |  | MET181 |  | ASP230 | -1 | TYR279 |  | LYS328 | +1 |
| GLN35 |  | ILE84 |  | LYS133 | +1 | GLU182 | -1 | MET231 |  | LYS280 | +1 | TYR329 |  |
| PRO36 |  | PRO85 |  | LYS134 | +1 | TRP183 |  | SER232 |  | SER281 |  | LYS330 | +1 |
| ALA37 |  | ILE86 |  | ALA135 |  | LEU184 |  | ALA233 |  | VAL282 |  | ASP331 | -1 |
| LEU38 |  | GLN87 |  | TRP136 |  | SER185 |  | ILE234 |  | LEU283 |  | TYR332 |  |
| THR39 |  | VAL88 |  | VAL137 |  | LEU186 |  | THR235 |  | HID284 |  | GLU333 | -1 |
| THR40 |  | ILE89 |  | ASP138 | -1 | GLY187 |  | VAL236 |  | ARG285 | +1 | TYR334 |  |
| SER41 |  | GLU90 | -1 | TYR139 |  | LEU188 |  | LEU237 |  | THR286 |  | CYS335 |  |
| ALA42 |  | ARG91 | +1 | LEU140 |  | GLY189 |  | VAL238 |  | THR287 |  | LYS336 | +1 |
| GLY43 |  | LEU92 |  | PHE141 |  | LEU190 |  | PRO239 |  | VAL288 |  | LEU337 |  |
| PRO44 |  | GLN93 |  | HID142 |  | GLU191 | -1 | ASN240 |  | ASN289 |  | ASN338 |  |
| VAL45 |  | ARG94 | +1 | ASN143 |  | LYS192 | +1 | ASP241 | -1 | LYS290 | +1 | LYS339 | +1 |
| PRO46 |  | ALA95 |  | VAL144 |  | ASN193 |  | VAL242 |  | GLU291 | -1 | LEU340 |  |
| GLU47 | -1 | GLY96 |  | TRP145 |  | ALA194 |  | PRO243 |  | LYS292 | +1 | PRO341 |  |
| ILE48 |  | LYS97 | +1 | PRO146 |  | LEU195 |  | GLY244 |  | ALA293 |  | GLN342 |  |
| PRO49 |  | GLU98 | -1 | PRO147 |  | ILE196 |  | LEU245 |  | ARG294 | +1 |  |  |

**Supplementary Table 12.**
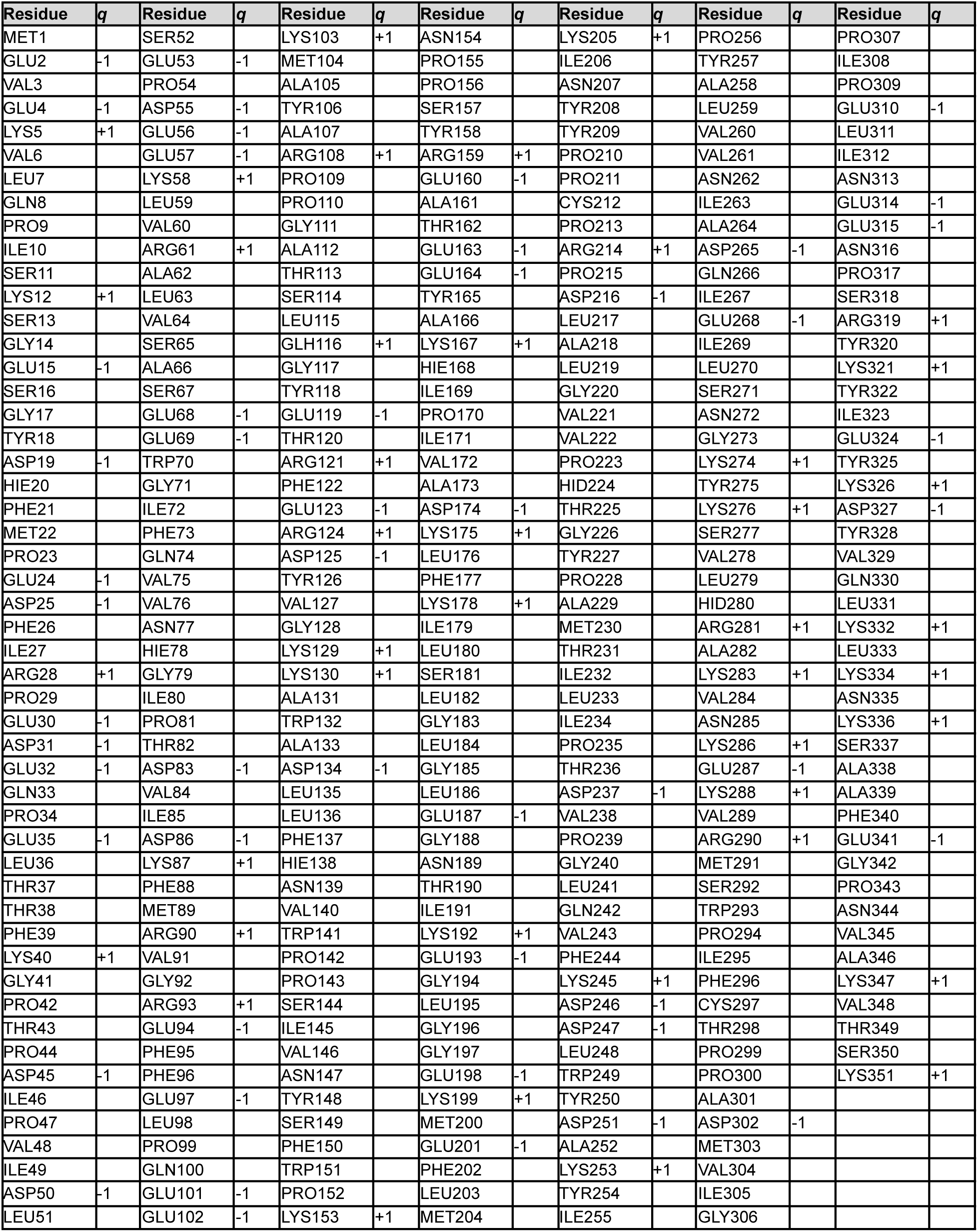
DAH protonation states and charges (*q*) with neutral charges left blank.

**Supplementary Table 13.**
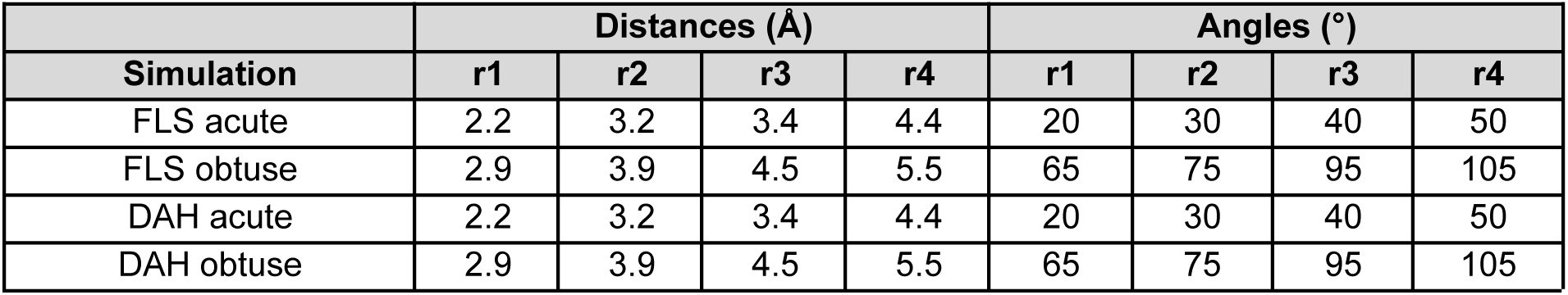
Flat-bottom harmonic restraints derived from experimentally motivated distance (Å) and angle (°) values were applied to each system. In all cases, the two restraints were the distance between the hydrogen atom target and the iron center (Fe H), and the angle between the hydrogen atom target, the iron center, and the oxo ligand (∠H-Fe-oxo). The flat bottom of the restraint spans r2 and r3, and the force is applied with a harmonic force constant between r1 and r2 as well as between r3 and r4. The restraint is centered on the experimental value with a width of the experimental uncertainty.

| Simulation | Distances (Å) |  |  |  | Angles (°) |  |  |  |
| --- | --- | --- | --- | --- | --- | --- | --- | --- |
|  | r1 | r2 | r3 | r4 | r1 | r2 | r3 | r4 |
| FLS acute | 2.2 | 3.2 | 3.4 | 4.4 | 20 | 30 | 40 | 50 |
| FLS obtuse | 2.9 | 3.9 | 4.5 | 5.5 | 65 | 75 | 95 | 105 |
| DAH acute | 2.2 | 3.2 | 3.4 | 4.4 | 20 | 30 | 40 | 50 |
| DAH obtuse | 2.9 | 3.9 | 4.5 | 5.5 | 65 | 75 | 95 | 105 |

**Supplementary Table 14.** DBSCAN clustering statistics for all FLS MD simulations. The CPPTraj implementation was used and e was tuned incrementally until there were fewer than six clusters with the top three clusters shown. The clustering selection was chosen with the goal of identifying a representative frame for QM/MM. The centroid of the largest cluster was chosen. All other clusters were discarded. The clustering mask was selected to include the first and second coordination sphere, the substrate, the iron center, and residues in contact with the substrate. A large mask was used to identify a centroid that accurately captured the equilibrium conformation. The FLS mask was: L119, N127, F141, K209, H228, D230, S232, T235, H266, H284, R294, F300, S302, Y329, E333.

| FLS 2OG unrestrained |  |  |  |  |  |
| --- | --- | --- | --- | --- | --- |
| #Cluster | Frames | Frac | AvgDist | Stdev | Centroid |
| 0 | 115228 | 0.922 | 0.975 | 0.155 | 33670 |
| 1 | 4077 | 0.033 | 0.833 | 0.169 | 903 |
| FLS succinate unrestrained |  |  |  |  |  |
| #Cluster | Frames | Frac | AvgDist | Stdev | Centroid |
| 0 | 122172 | 0.977 | 1.127 | 0.222 | 94744 |
| 1 | 428 | 0.003 | 0.657 | 0.098 | 116362 |
| FLS succinate acute |  |  |  |  |  |
| #Cluster | Frames | Frac | AvgDist | Stdev | Centroid |
| 0 | 117864 | 0.943 | 0.942 | 0.172 | 76269 |
| 1 | 476 | 0.004 | 0.615 | 0.075 | 6916 |
| 2 | 369 | 0.003 | 0.597 | 0.065 | 6392 |

**Supplementary Table 15.** DBSCAN clustering statistics for all DAH MD simulations. The CPPTraj implementation was used and e was tuned incrementally until there were fewer than six clusters with the top three shown. The clustering selection was chosen with the goal of identifying a representative frame for QM/MM. The centroid of the largest cluster was chosen. All other clusters were discarded. The clustering mask was selected to include the first and second coordination sphere, the substrate, the iron center, and residues in contact with the substrate. A large mask was used to identify a centroid that accurately captured the equilibrium conformation. The DAH mask was: L115, R121, L135, F137, K205, N207, H224, Y227, P228, T231, F296, H280, L333.

| DAH 2OG unrestrained |  |  |  |  |  |
| --- | --- | --- | --- | --- | --- |
| #Cluster | Frames | Frac | AvgDist | Stdev | Centroid |
| 0 | 66508 | 0.532 | 1.273 | 0.322 | 61940 |
| 1 | 33415 | 0.267 | 0.989 | 0.233 | 29207 |
| 2 | 9578 | 0.077 | 0.960 | 0.175 | 115829 |
| 3 | 5693 | 0.046 | 1.077 | 0.202 | 109300 |
| DAH succinate eq-oxo unrestrained |  |  |  |  |  |
| #Cluster | Frames | Frac | AvgDist | Stdev | Centroid |
| 0 | 75121 | 0.601 | 0.856 | 0.147 | 80588 |
| 1 | 19803 | 0.158 | 1.703 | 0.546 | 46050 |
| 2 | 18945 | 0.152 | 1.212 | 0.311 | 6133 |
| DAH succinate eq-oxo obtuse |  |  |  |  |  |
| #Cluster | Frames | Frac | AvgDist | Stdev | Centroid |
| 0 | 57887 | 0.528 | 0.974 | 0.213 | 64903 |
| 1 | 50243 | 0.458 | 1.362 | 0.275 | 21716 |
| DAH succinate ax-oxo unrestrained |  |  |  |  |  |
| #Cluster | Frames | Frac | AvgDist | Stdev | Centroid |
| 0 | 107202 | 0.858 | 1.146 | 0.325 | 41101 |
| 1 | 550 | 0.004 | 0.703 | 0.096 | 105067 |
| 2 | 494 | 0.004 | 0.654 | 0.102 | 326 |
| DAH succinate ax-oxo obtuse |  |  |  |  |  |
| #Cluster | Frames | Frac | AvgDist | Stdev | Centroid |
| 0 | 53171 | 0.425 | 0.782 | 0.123 | 114031 |
| 1 | 47607 | 0.381 | 1.101 | 0.342 | 66374 |
| 2 | 15578 | 0.125 | 0.952 | 0.233 | 14977 |

**Supplementary Table 16.** The residues in the QM region for the QM/MM FLS simulations with 2OG. The table contains the total number of atoms prior to adding link atoms along with their corresponding net charge.

| **Residue** | **# of atoms** | **Charge** |
| --- | --- | --- |
| Lys209 | 22 | 1 |
| His228 | 17 | 0 |
| Asp230 | 12 | -1 |
| His284 | 17 | 0 |
| Fe | 1 | 2 |
| 2-oxoglutarate (2OG) | 14 | -2 |
| Dihydrokaempferol | 33 | 0 |
| **Total** | **116** | **0** |

**Supplementary Table 17.** The residues in the QM region for the QM/MM FLS simulations with succinate. The table contains the total number of atoms prior to adding link atoms along with their corresponding net charge.

| **Residue** | **# of atoms** | **Charge** |
| --- | --- | --- |
| Lys209 | 22 | 1 |
| His228 | 17 | 0 |
| Asp230 | 12 | -1 |
| His284 | 17 | 0 |
| Fe | 1 | 4 |
| Oxo | 1 | -2 |
| Succinate | 12 | -2 |
| Dihydrokaempferol | 33 | 0 |
| **Total** | **115** | **0** |

**Supplementary Table 18.** The residues in the QM region for the QM/MM DAH simulations with 2OG. The table contains the total number of atoms prior to adding link atoms along with their corresponding net charge.

| **Residue** | **# of atoms** | **Charge** |
| --- | --- | --- |
| Lys205 | 22 | 1 |
| His224 | 17 | 0 |
| His280 | 16 | 0 |
| Fe | 1 | 2 |
| Cl | 1 | -1 |
| 2-oxoglutarate (2OG) | 14 | -2 |
| Dechloroacutumine | 54 | 0 |
| **Total** | **125** | **0** |

**Supplementary Table 19.** The residues in the QM region for the QM/MM DAH simulations with succinate. The table contains the total number of atoms prior to adding link atoms along with their corresponding net charge.

| **Residue** | **# of atoms** | **Charge** |
| --- | --- | --- |
| Lys205 | 22 | 1 |
| His224 | 17 | 0 |
| His280 | 16 | 0 |
| Fe | 1 | 4 |
| Oxo | 1 | -2 |
| Cl | 1 | -1 |
| Succinate | 12 | -2 |
| Dechloroacutumine | 54 | 0 |
| **Total** | **124** | **0** |

### Supplementary Figures

**Supplementary Fig. 1.**
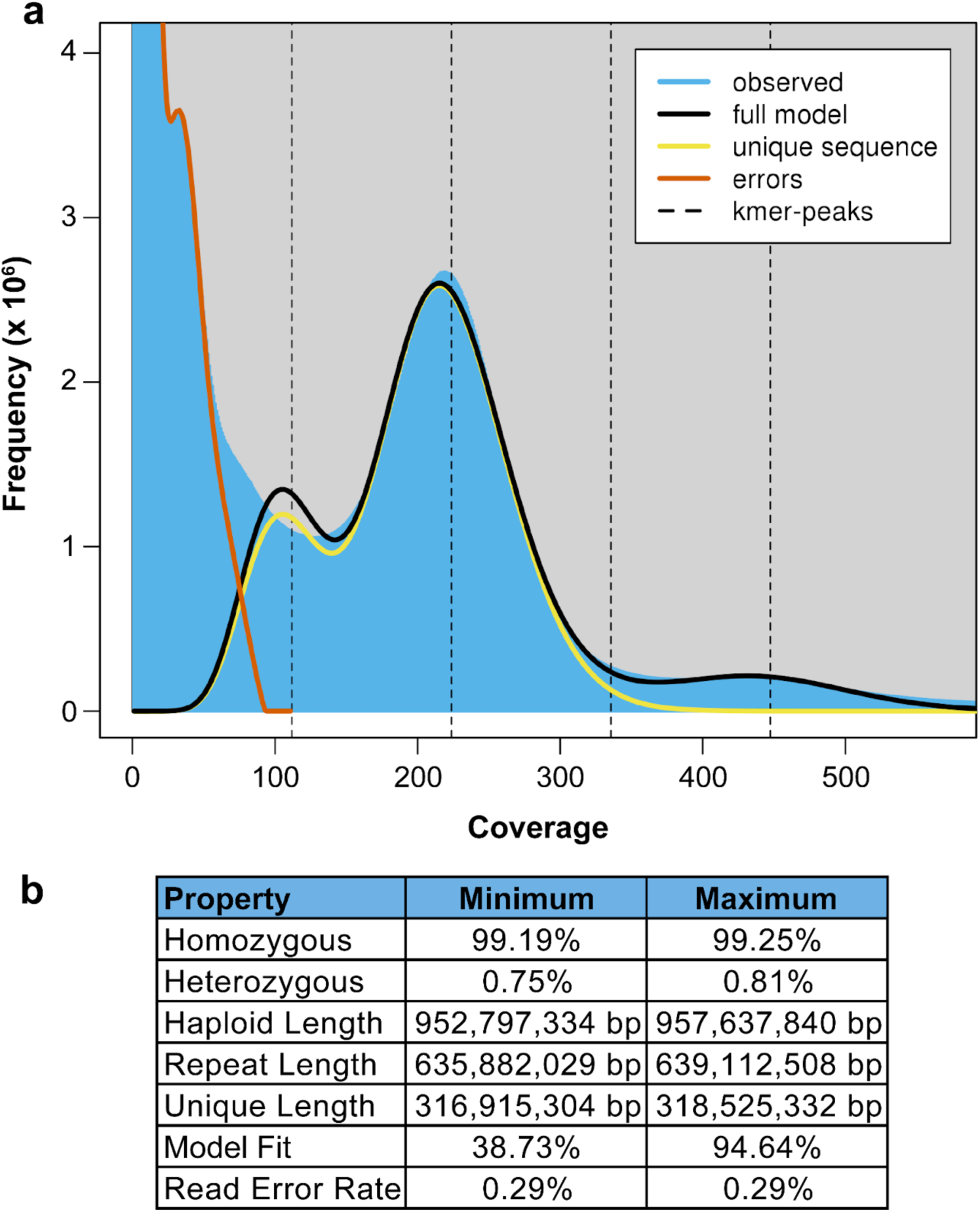
Genome size estimation by *k*-mer analysis (*k* = 19). (**a**) Linear plot of model fit output from GenomeScope^5^. (**b**) Estimations and results collected from GenomeScope analysis.

**Supplementary Fig. 2.**
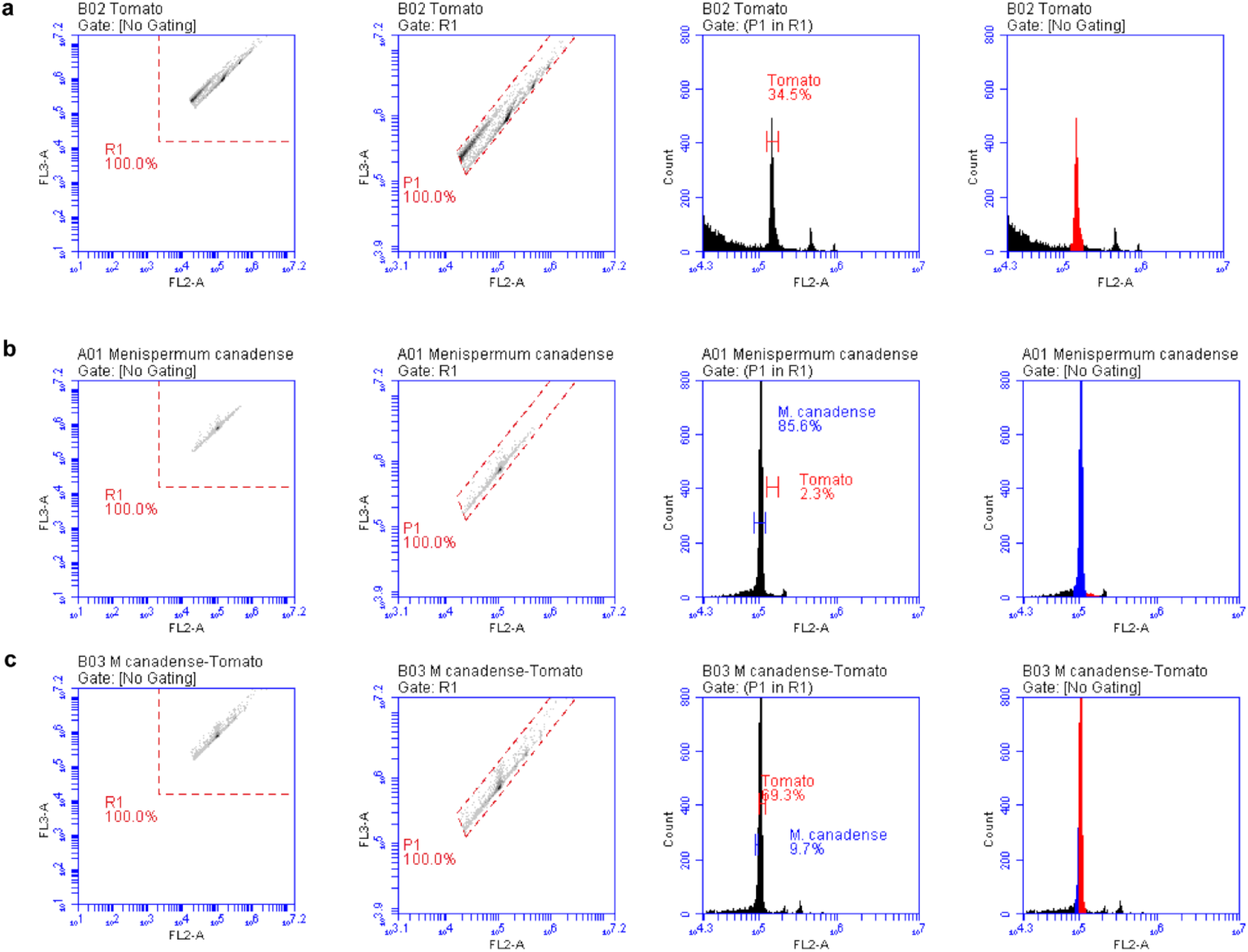
Genome size estimation by cell flow-cytometry (BD Accuri™ C6 Cytometer). Flow cytometry measurements for (**a**) *Solanum lycopersicum* (tomato), (**b**) *M. canadense*, and (**c**) *M. canadense* and tomato. The estimated genome size of tomato is 2.05 Gb (2n DNA content)^6^.

**Supplementary Fig. 3.**
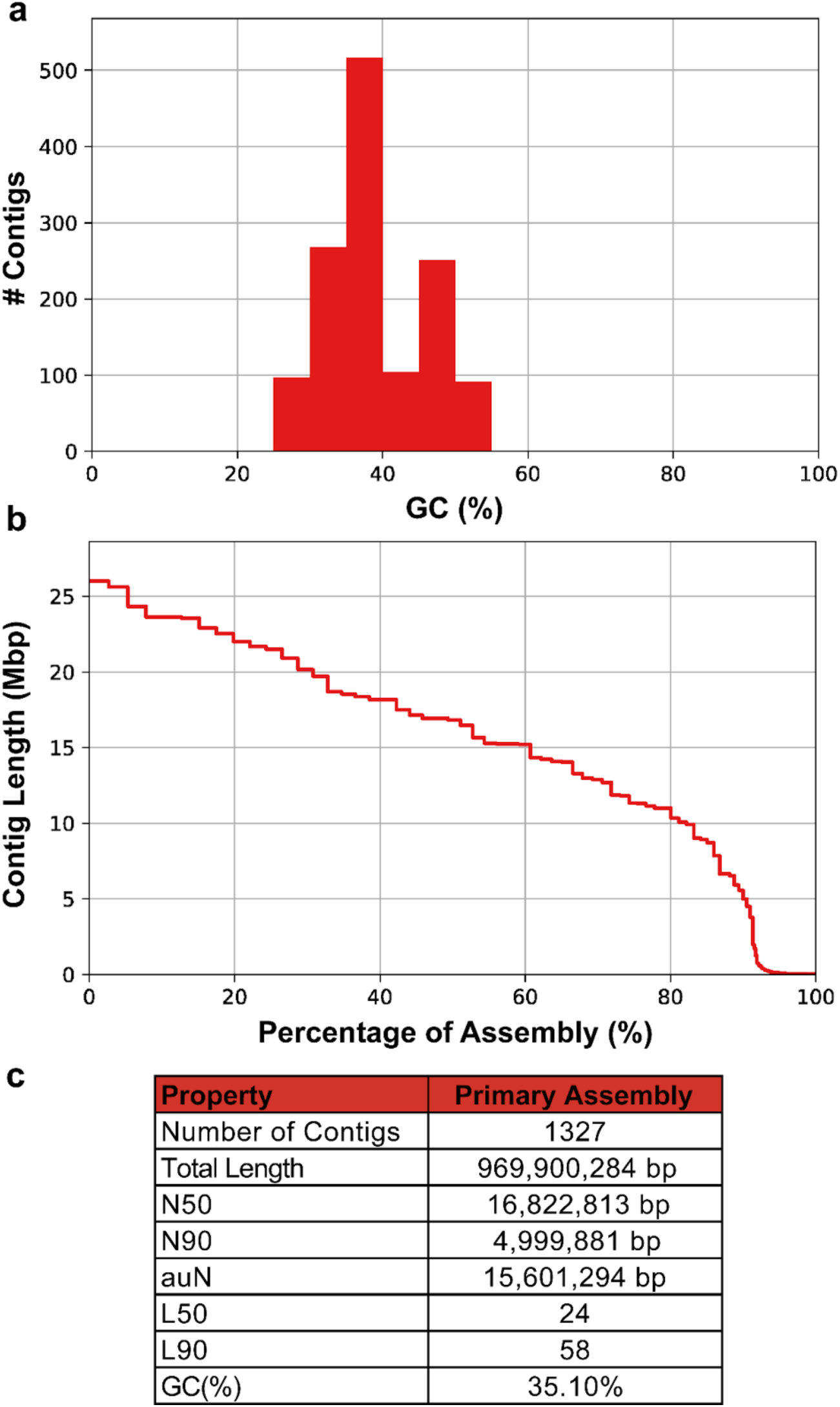
Genome assembly statistics for initial contig-level assembly. (**a**) GC% of initial assembly (**b**) Contig length (N_x_) of genome assembly (**c**) Genome assembly statistics.

**Supplementary Fig. 4.**
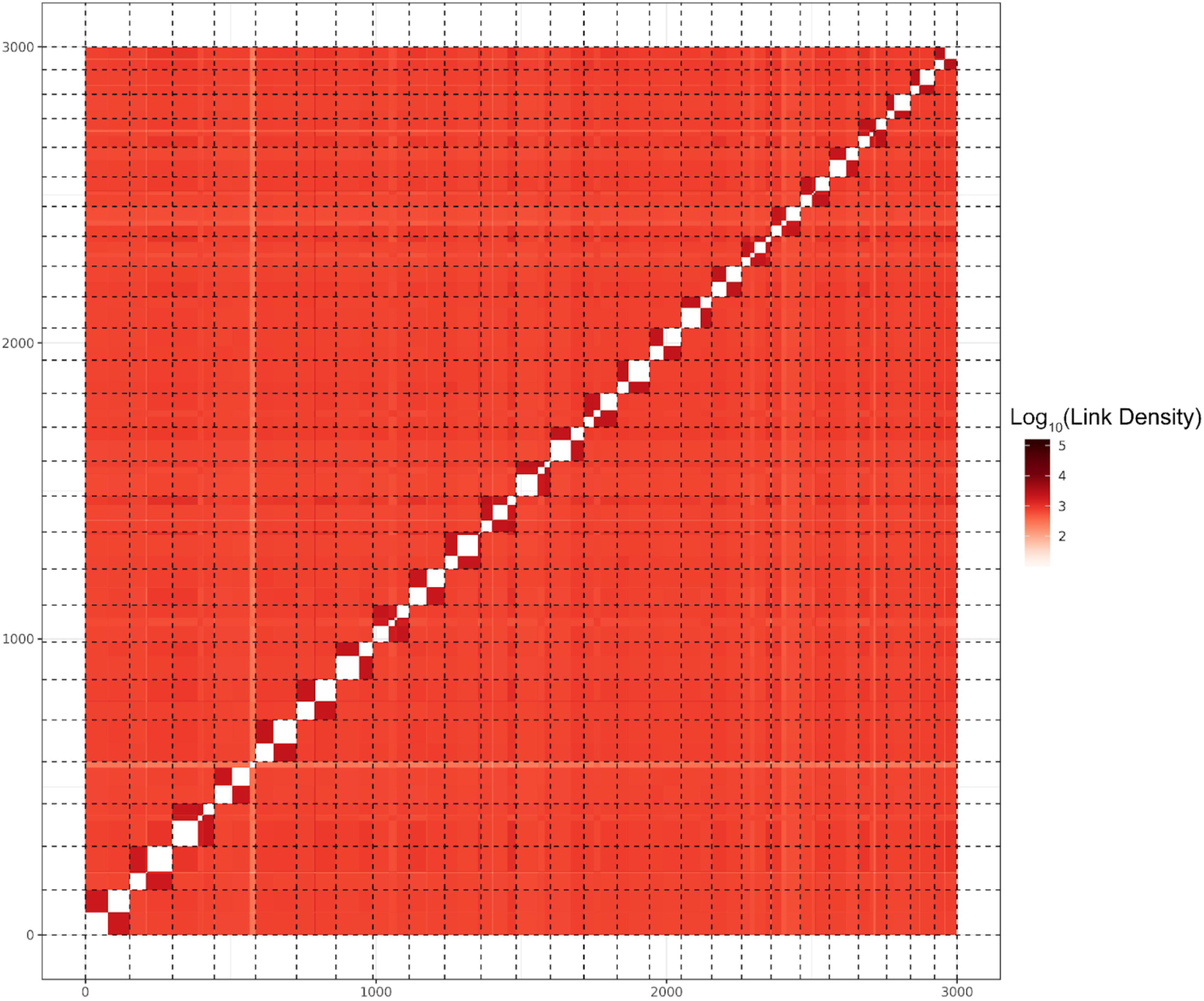
Scaffolding of *M. canadense* genome using Hi-C information. Contact map showcasing the log_10_ of link density between *M. canadense* genome to itself with chunk size of 301,424 bp.

**Supplementary Fig. 5.**
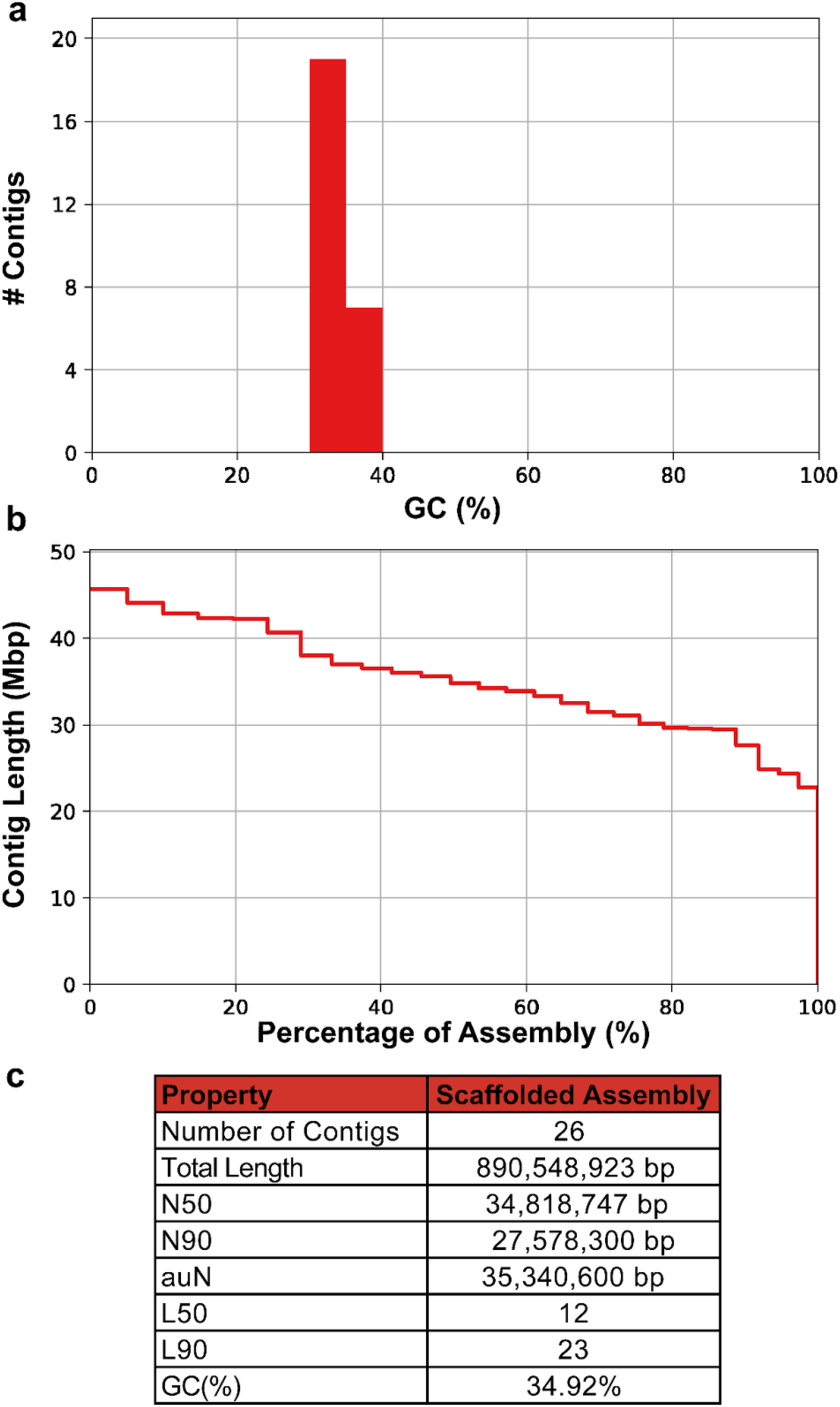
Genome assembly statistics for post-scaffolded chromosomal-level assembly of *M. canadense*. (**a**) GC% of scaffolded assembly (**b**) Contig length (N_x_) of genome assembly (**c**) Genome assembly statistics.

**Supplementary Fig. 6.**
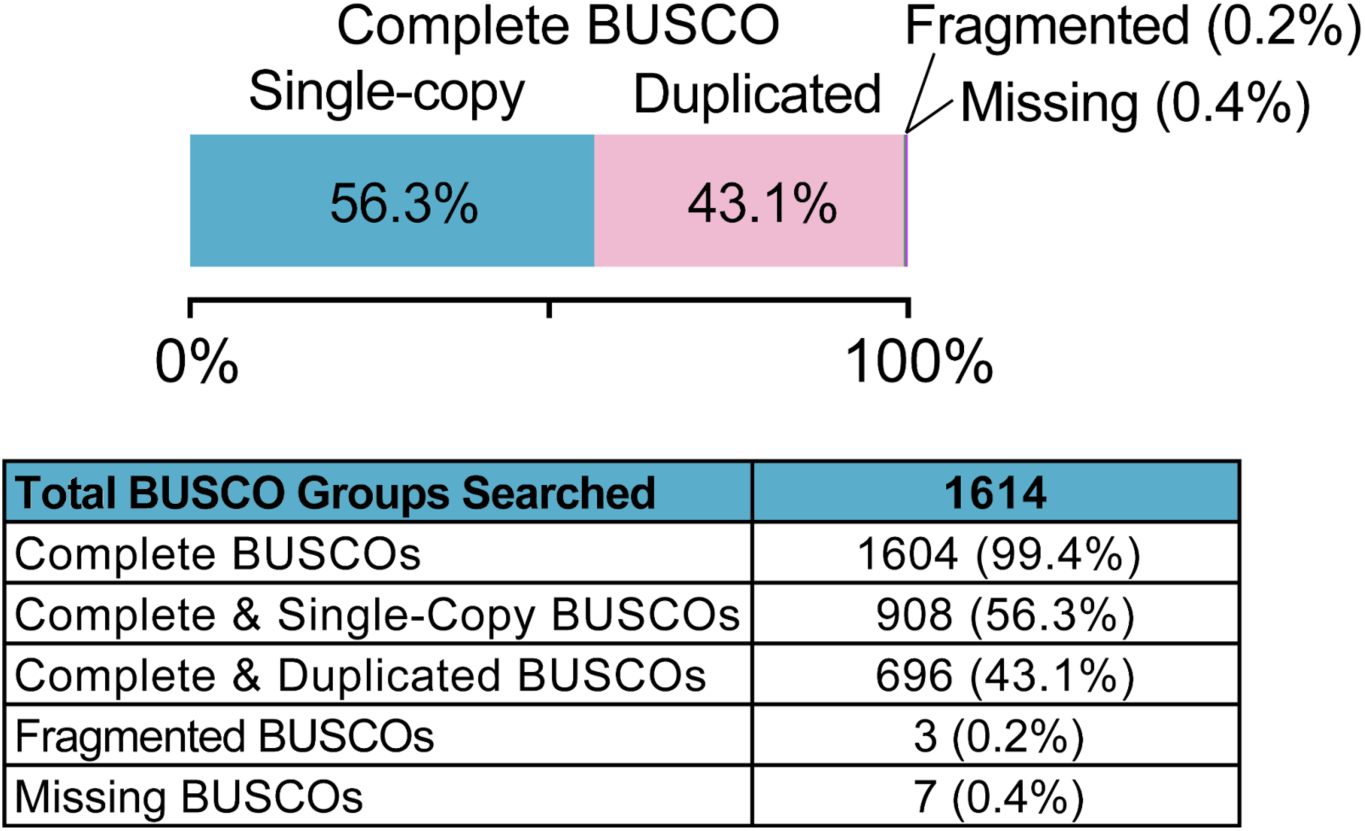
Benchmarking Universal Single-Copy Orthologs (BUSCO) assessment of *M. canadense* chromosomal-level assembly. The lineage dataset used for BUSCO assessment consists of 1614 genes in 50 embryophyta genomes (embryophyta_odb_10). 43.1% of the completed BUSCO genes exist as duplicates, which is a similar observation found in the *P. somniferum* genome (62%) that went under a relatively recent WGD^7^.

**Supplementary Fig. 7.**
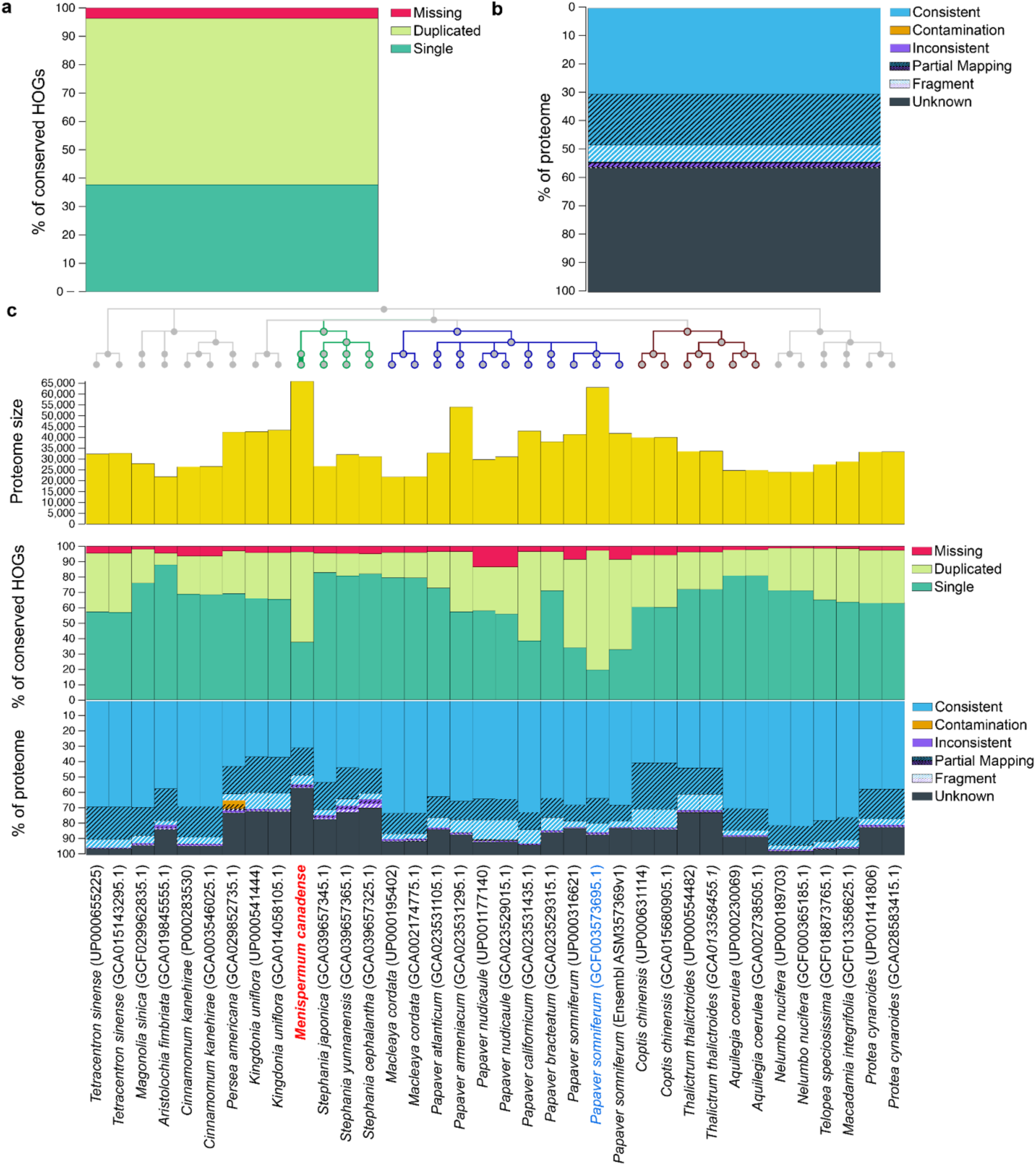
OMArk comparison of *M. canadense* proteome to the expected gene repertoire of its common ancestor. (**a**) Completeness statistic showing the percent of conserved Hierarchical Orthologous Groups (HOGs) that are represented as single (light green) or duplicated genes (teal), or entirely missing (red) in *M. canadense* proteome. (**b**) Whole proteome assessment of gene model consistency with known homologs present in comparison to gene contents of all extant species of the same lineage to *M. canadense*. Proportion of genes whose closest gene families are from the selected lineage (consistent; blue), from another lineage as contamination (contamination; orange), from another lineage as noise (inconsistent; purple), and with no closest homologs found (unknown; grey). Partial mapping refers to genes that have less than 80% of the sequence with shared k-mer content from its closest gene family. Fragments indicate genes with a length less than half the median gene content of its closest gene family. There is 58.67% duplication (4768 out of 8127) of HOGs in *M. canadense* proteome, similar to its duplicated BUSCOs in Supplementary Fig. 6. (**c**) Comparison of *M. canadense* proteome statistics with other publicly available Mesangiospermae (core angiosperm) species. *M. canadense* and *P. somniferum* proteomes have relatively high duplicated HOGs. Green branches: Menispermaceae, blue branches: Papaveraceae, brown: Ranunculaceae.

**Supplementary Fig. 8.**
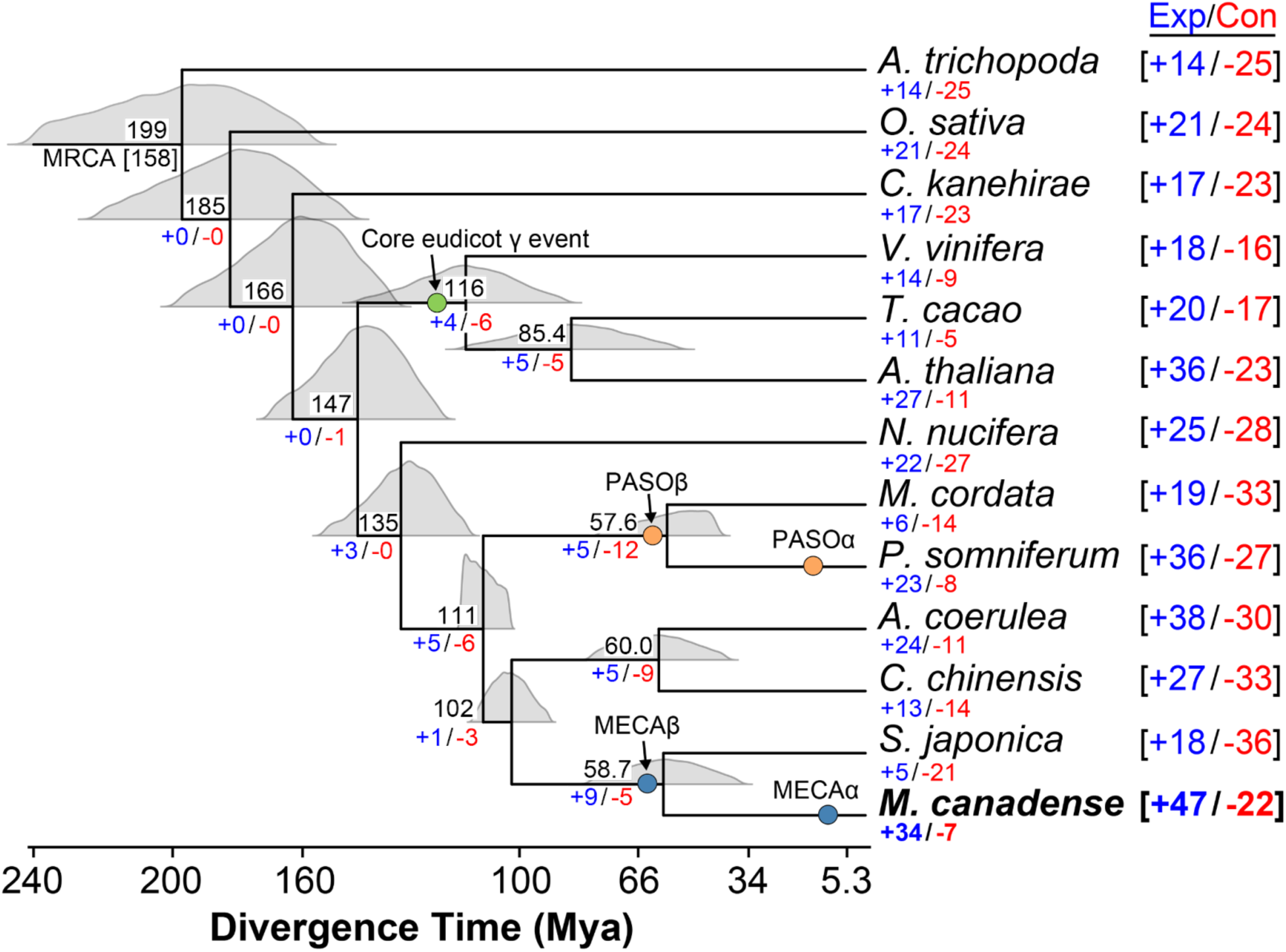
2ODD-specific gene family expansion and contraction. Phylogenetic divergence time of *M. canadense* compared to 12 other angiosperm species as shown in Fig. 1c. Black number at the top of each tree node indicates the mean divergence time million years ago (Mya). The blue and red numbers at the bottom of each tree node indicate expansion and contraction, respectively, of 158 gene families with functional annotations to 2ODD. Green circle indicates the relative timing of the core eudicot γ event, whereas orange circles correspond to two WGD events present in *P. somniferum*. Blue circles indicate the relative timing of two predicted WGD events in *M. canadense*.

**Supplementary Fig. 9.**
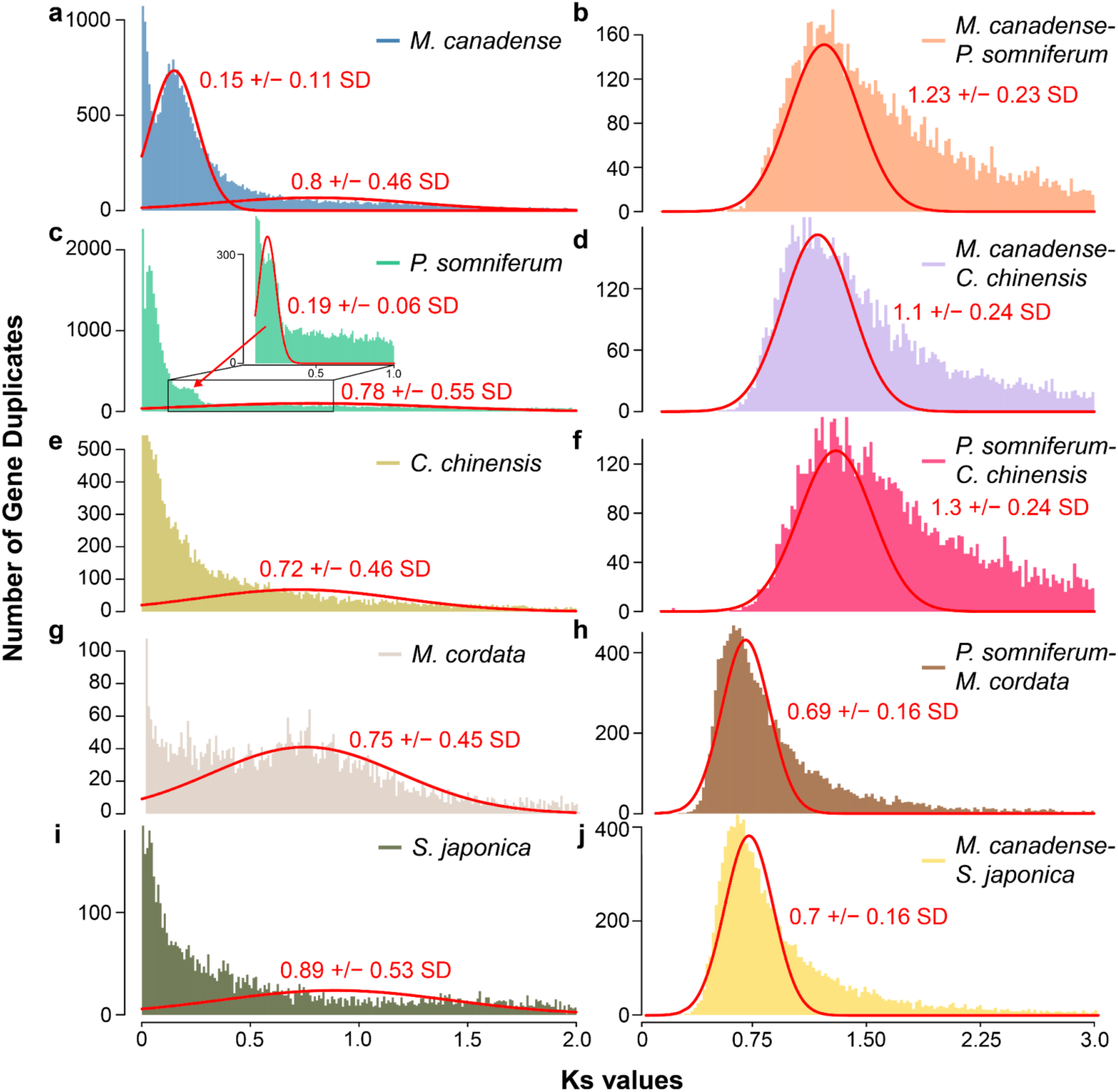
Mixtools analyses on distribution of the RBH paralogous gene pair *K*_S_ values for *M. canadense*, *P. somniferum*, *C. chinensis, M. cordata*, and *S. japonica*. *K*_S_ distribution peaks determined by Mixtools analyses using 1000 bootstraps for (**a**) *M. canadense* paralogs, (**b**) *M. canadense* and *P. somniferum* orthologs (**c**) *P. somniferum* paralogs, (**d**) *M. canadense* and *C. chinensis* orthologs, (**e**) *C. chinensis* paralogs, (**f**) *P. somniferum* and *C. chinensis* orthologs, (**g**) *M. cordata* paralogs, (**h**) *P. somniferum* and *M. cordata* orthologs, (**i**) *S. japonica* paralogs, and (**j**) *M. canadense* and *S. japonica* orthologs.

**Supplementary Fig. 10.**
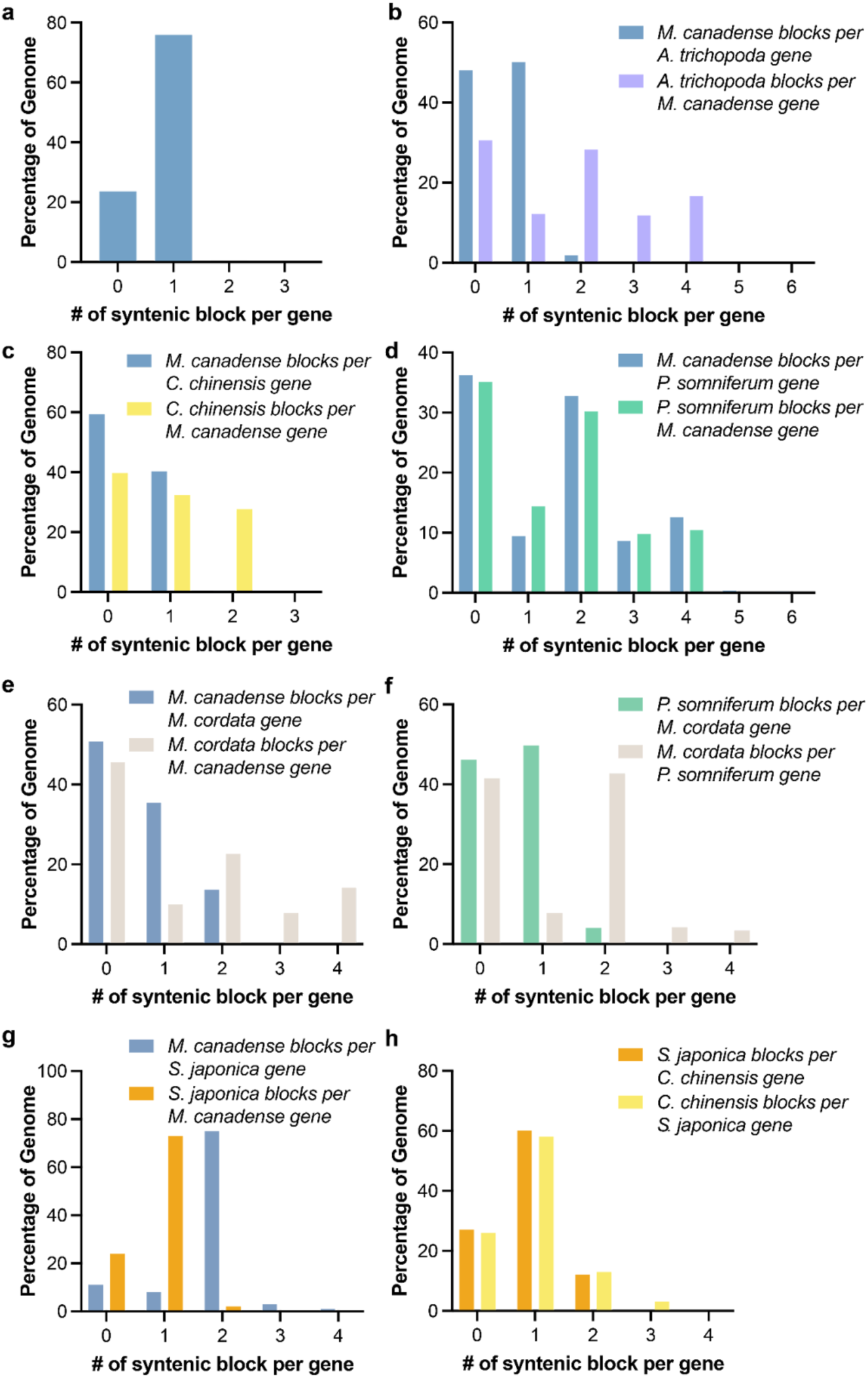
The syntenic depth between *M. canadense* and select plant genomes. (**a**) *M. canadense* vs *M. canadense* syntenic depths show 1:1 pattern. (**b**) *M. canadense* vs *A. trichopoda* syntenic depths show 4:1 pattern. (**c**) *M. canadense* vs *C. chinensis* syntenic depths show 2:1 pattern. (**d**) *M. canadense* vs *P. somniferum* syntenic depths show 4:4 pattern. (**e**) *M. canadense* vs *M. cordata* syntenic depths show 4:2 pattern. (**f**) *P. somniferum* vs *M. cordata* syntenic depths show 2:1 pattern. (**g**) *M. canadense* vs *S. japonica* syntenic depths show 2:1 pattern. (**h**) *S. japonica* vs *C. chinensis* syntenic depths show 2:2 pattern.

**Supplementary Fig. 11.**
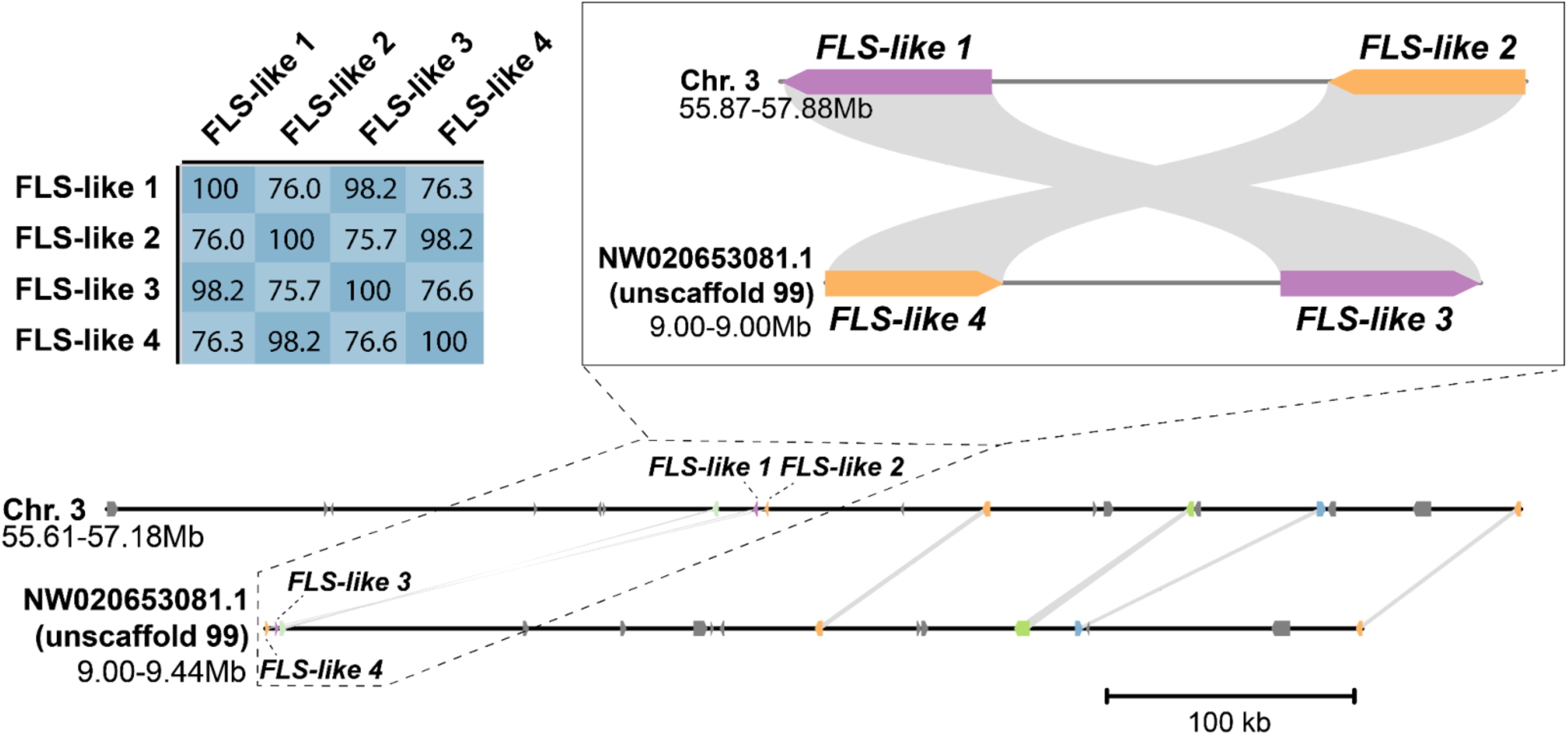
Microsynteny analysis of *FLS-like* gene locus in *P. somniferum*. WGD region in chromosome 3 and unscaffold 99 in *P. somniferum* genome containing paralogous tandem duplicated *FLS-like* genes. Amino-acid sequence identity matrix of *FLS-like 1*, *FLS-like 2*, *FLS-like 3*, and *FLS-like 4*, calculated by local pairwise alignments using EMBOSS supermatcher^8^.

**Supplementary Fig. 12.**
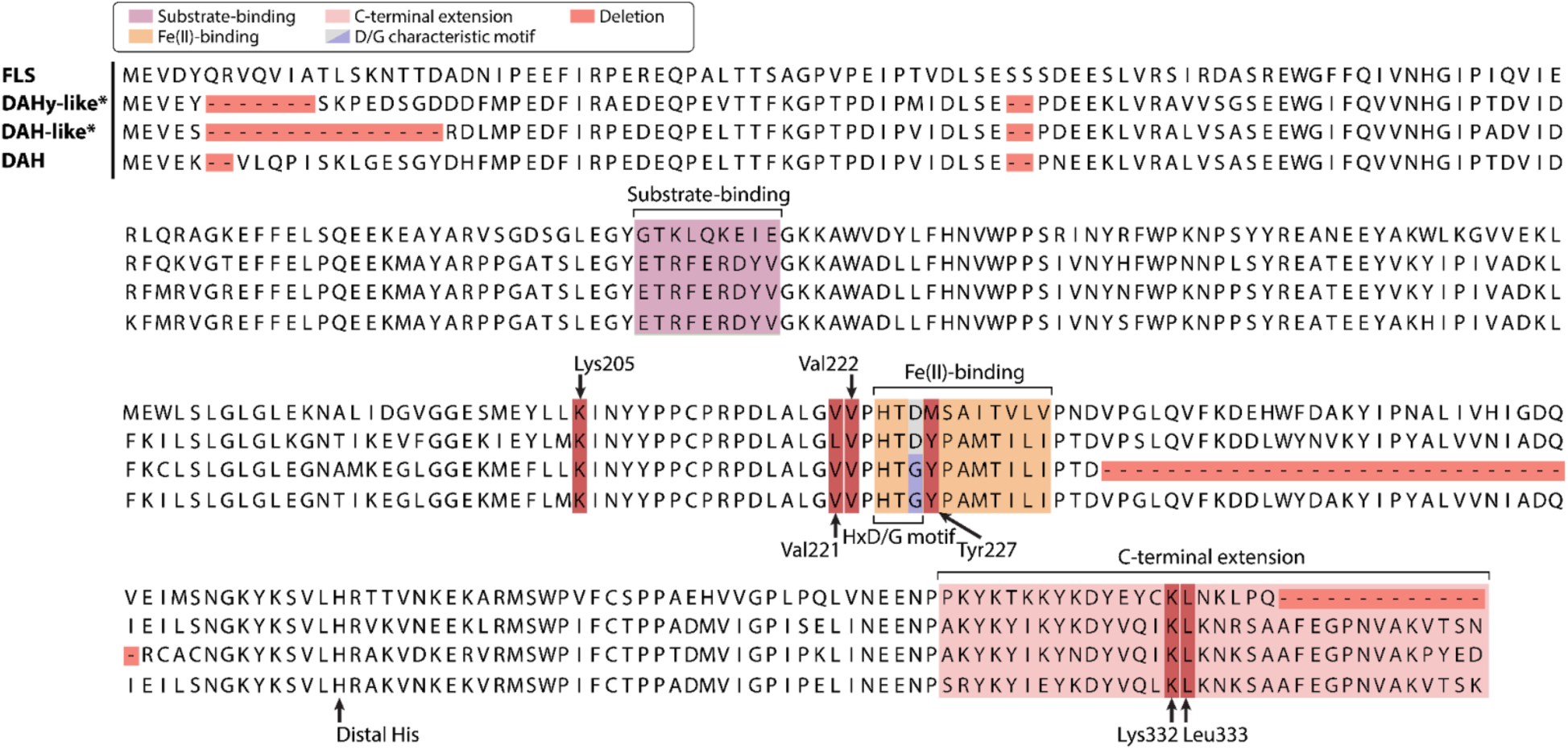
Multiple sequence alignment of FLS and DAH paralogs found in chromosomes 2 and 3. Relevant residues and motifs mentioned in this manuscript are highlighted according to the color legend and labels.

**Supplementary Fig. 13.**
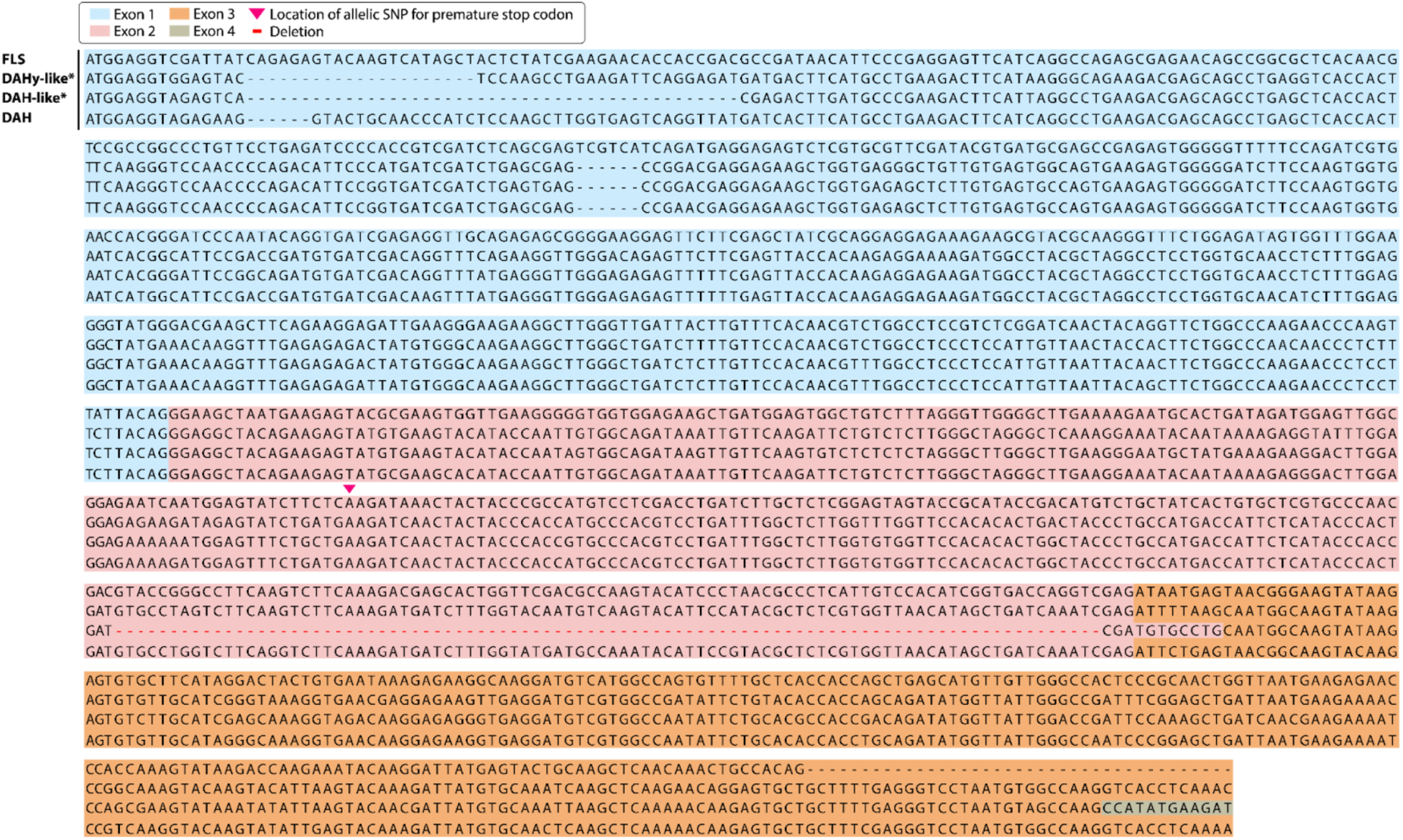
Multiple sequence codon alignment of *FLS* and *DAH* paralogous genes found in chromosomes 2 and 3. Relevant residues and motifs mentioned in this manuscript are highlighted according to the color legend and labels.

**Supplementary Fig. 14.**
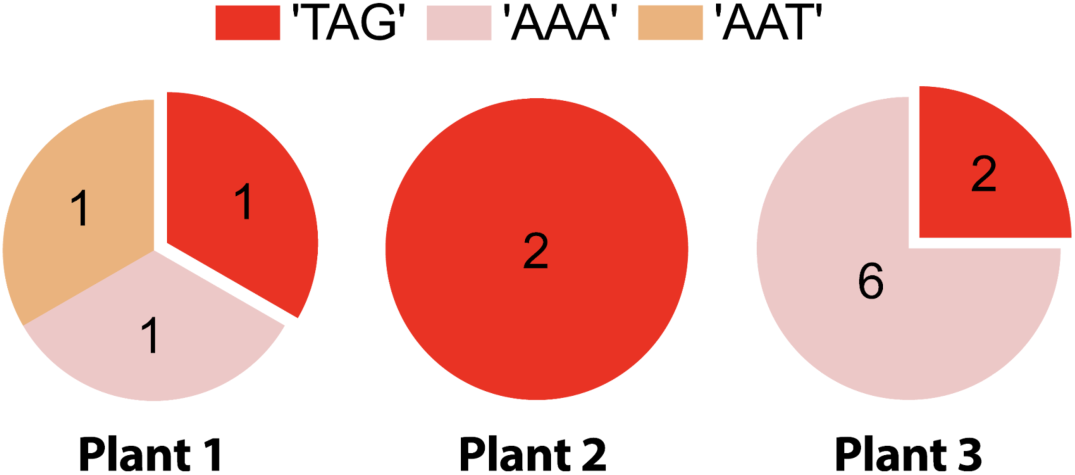
Total count of RNA-seq mapped reads in *DAHy-like* Lys200 codon. Quantification of mapped sequences for independent *M. canadense* plant RNA-sequencing samples.

**Supplementary Fig. 15.**
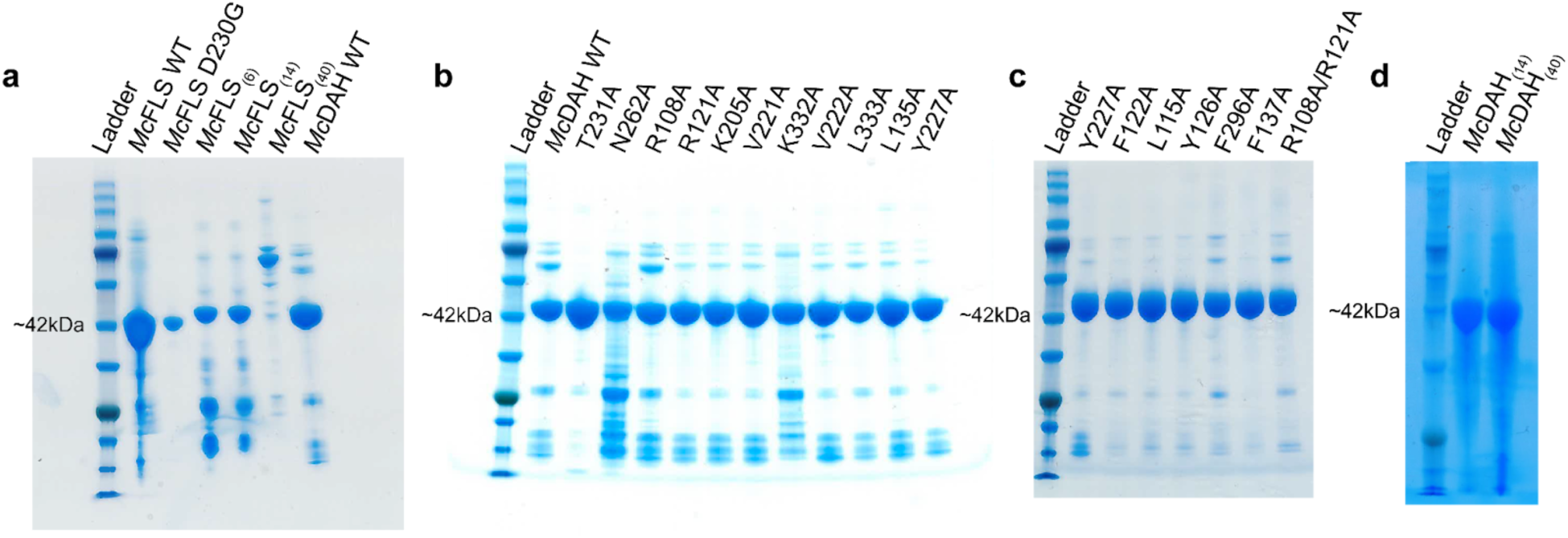
SDS-PAGE of recombinant proteins discussed in this study. (**a**) FLS-to-DAH swap mutant proteins. (**b** and **c**) DAH site-directed alanine mutants. (**d**) DAH-to-FLS swap mutant proteins.

**Supplementary Fig. 16.**
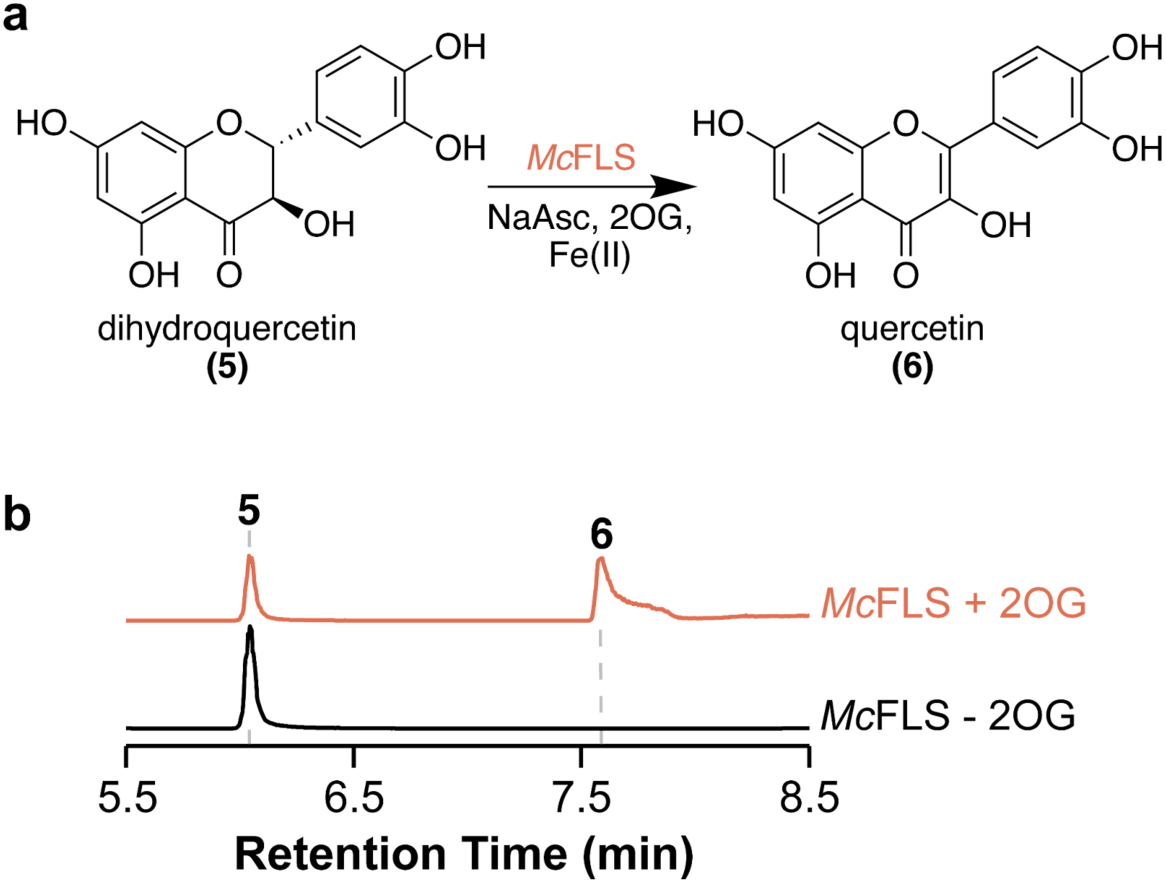
*In vitro* biochemical assay of FLS against dihydroquercetin. (**a**) Reaction schematic of the conversion of dihydroquercetin; **5** to quercetin; **6** using flavonol synthase (*Mc*FLS). (**b**) Combined LC-MS extracted ion chromatograms (EICs) of 303.05081 *m/z*; **5** = [M-H]^-^ and 301.03528 *m/z;* **6** = [M-H]^-^. XICs show the in vitro activity of *Mc*FLS that desaturate **5** to **6** in a 2OG-dependent manner.

**Supplementary Fig. 17.**
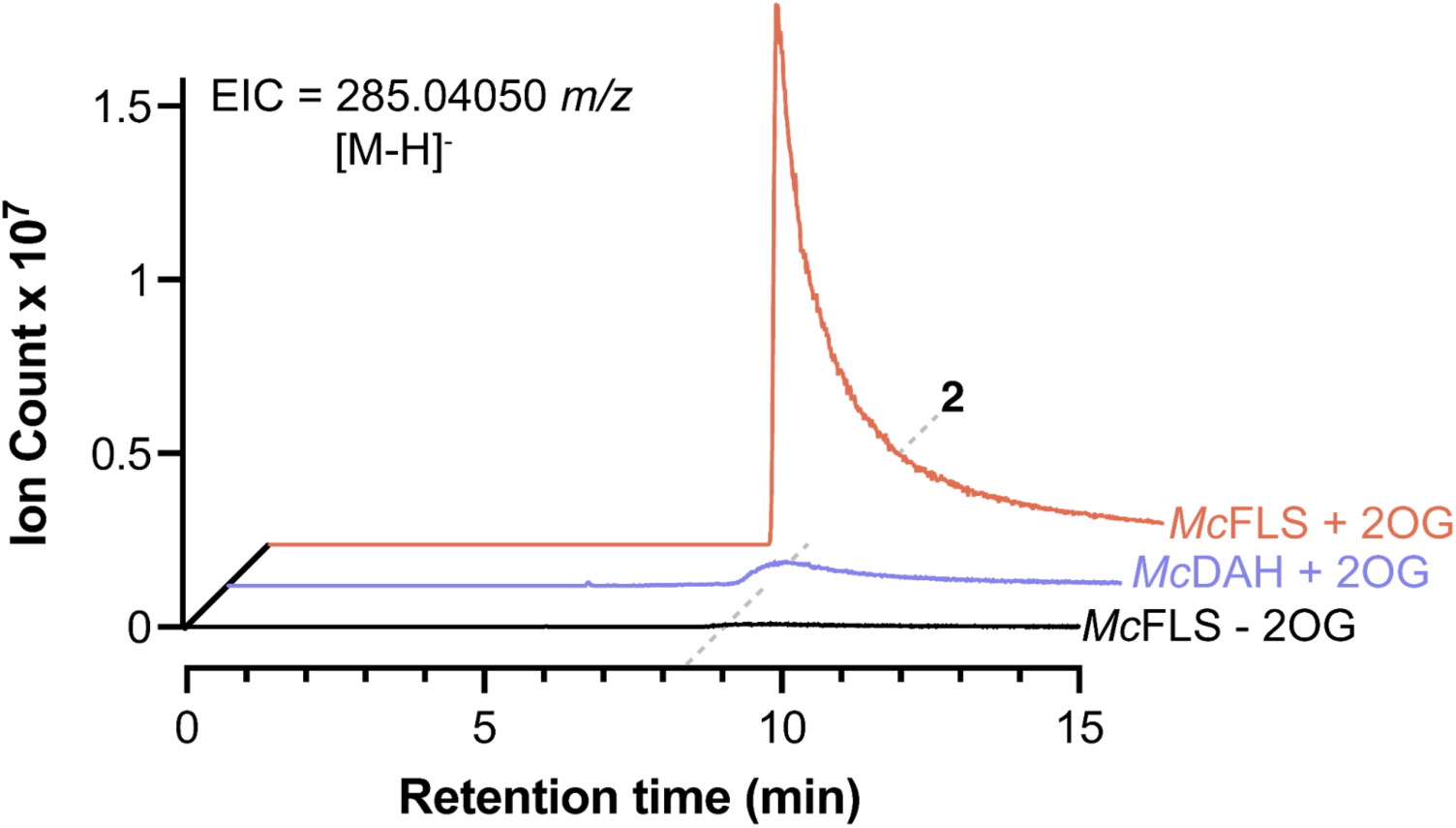
Production of trace amounts of kaempferol in *Mc*DAH assay against dihydrokaempferol. LC-HRAM-MS extracted ion chromatogram (EIC) of 285.04050 *m/z*; **2** = [M-H]^-^. *Mc*DAH produces trace amounts of **2** in the reaction as indicated by a small peak in *Mc*DAH + 2OG sample trace compared to *Mc*FLS - 2OG sample.

**Supplementary Fig. 18.**
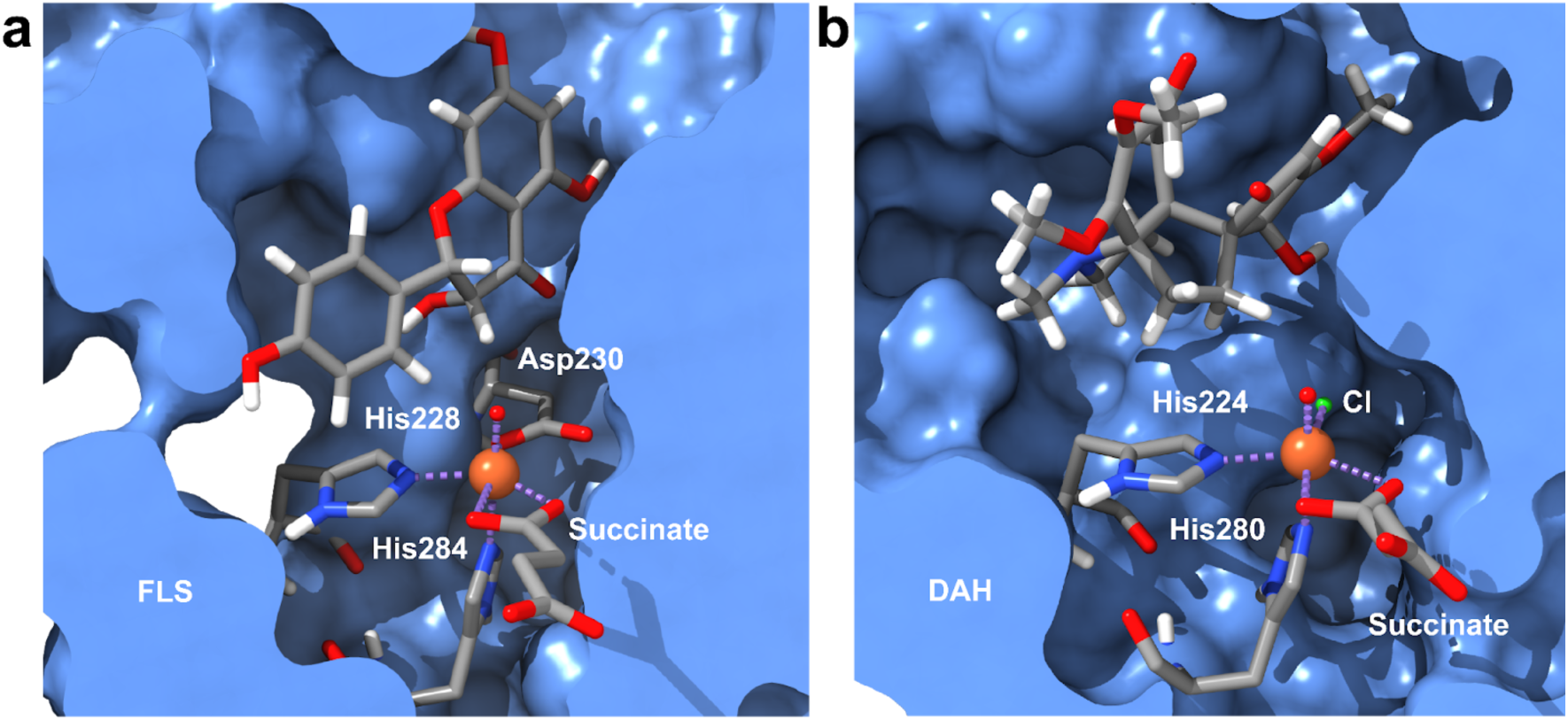
Docked AlphaFold structures of the enzyme-substrate complexes using AutoDock Vina. (**a**) Surface representation of the proposed FLS active site with dihydrokaempferol in its docked conformation. (**b**) Surface representation of the proposed DAH active site with dechloroacutumine in its docked conformation. The modeled succinate and oxo bound structure is included to illustrate the substrate position relative to the active site. The protein surface is shown in light blue, carbons in gray, nitrogens in blue, oxygens in red, hydrogens in white, and iron in orange. Dative bonds are shown as purple dashed lines.

**Supplementary Fig. 19.**
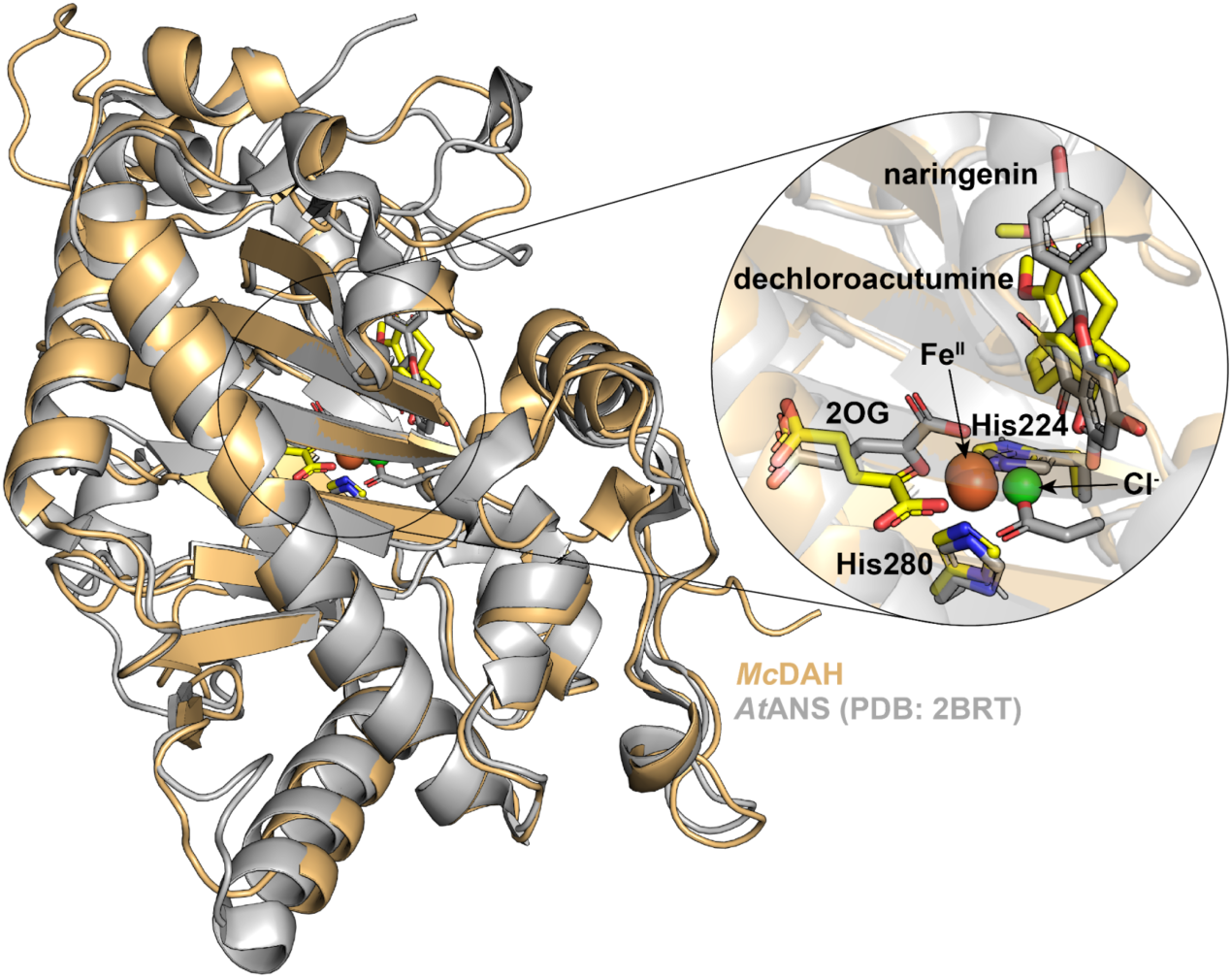
Structural alignment of *Mc*DAH AlphaFold structure and *At*ANS. The *McDAH* structure highlighted in gold is aligned to *At*ANS highlighted in gray (PDB: 2BRT). Naringenin, 2OG and Fe(II)-coordinating histidine triad with aspartate from *At*ANS structure are colored in gray, whereas the docked dechloroacutumine and Fe(II)-coordinating histidines from *Mc*DAH are colored in yellow. Fe(II) (orange) and Cl^-^ anion (green) positions are derived based on this structural alignment.

**Supplementary Fig. 20.**
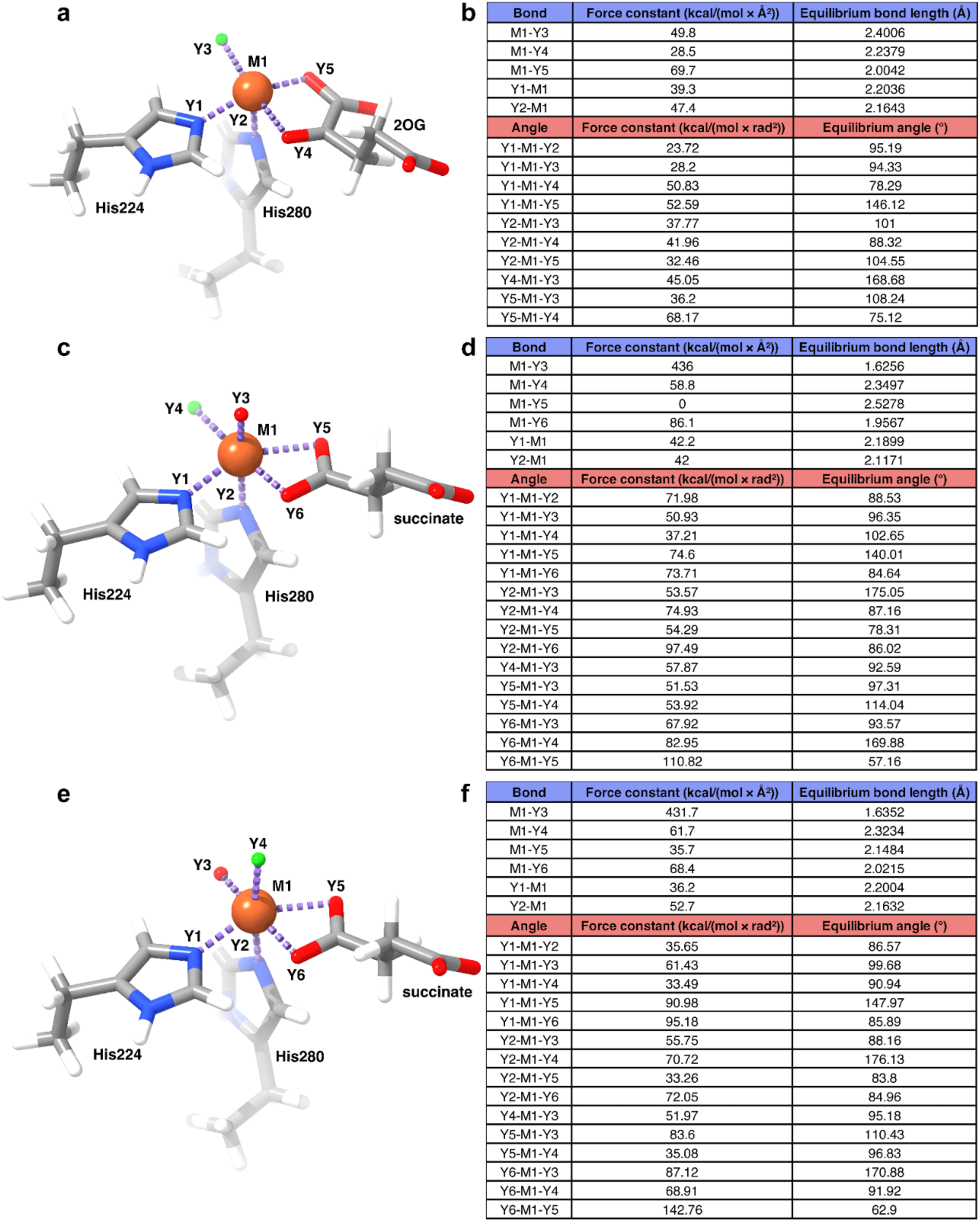
The core active site of *Mc*DAH structural models from AMBER’s Metal Center Parameter Builder (MCPB). DAH with (**a**) 2OG, (**c**) succinate and axial-oxo, or (**e**) succinate and equatorial-oxo in the MCPB small model with atom assignments labeled on the active site for coordinating atoms. Dative bonds are illustrated with purple dashed lines. Atoms are colored as follows: carbon in gray, nitrogen in blue, oxygen in red, hydrogen in white, and iron in orange. Parameters generated from DAH with (**b**) 2OG, (**d**) succinate and axial-oxo, or (**e**) succinate and equatorial-oxo force field parameters assigned using the metal center parameter builder (MCPB.py) from AmberTools22. Only the equilibrium bond lengths, angles, and their force constants that are directly bonded to the metal center are reported. Parameters were generated using the Seminario method.

**Supplementary Fig. 21.**
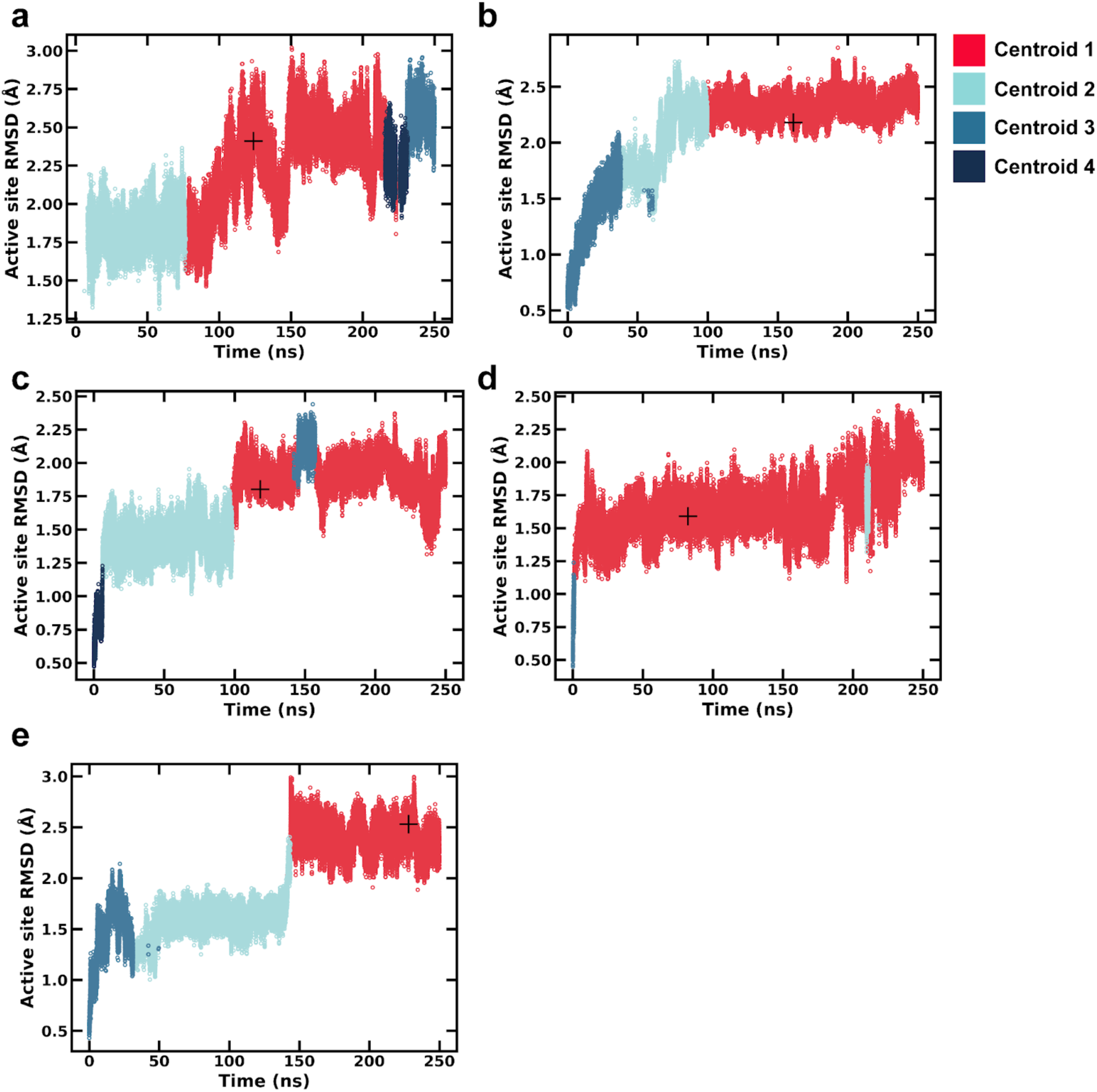
|. Clustered restrained MD simulations for DAH. The results of 250 ns MD simulations represented as time (ns) vs root mean squared deviation (Å) to the first frame. The simulations were clustered using CPPTraj and the DBSCAN method. (**a**) DAH 2OG simulation. Default clustering parameters were used with minpoints = 25 and = 0.75. (**b**) DAH succinate equatorial-oxo unrestrained simulation. Default clustering parameters were used with minpoints = 25 and = 0.76. (**c**) DAH succinate equatorial-oxo obtuse simulation. Default clustering parameters were used with minpoints = 25 and = 0.64. (**d**) DAH succinate axial-oxo unrestrained simulation. Default clustering parameters were used with minpoints = 25 and = 0.68. (**e**) DAH succinate axial-oxo obtuse simulation. Default clustering parameters were used with minpoints = 25 and = 0.66. The centroid of the largest cluster is marked with a crosshair.

**Supplementary Fig. 22.**
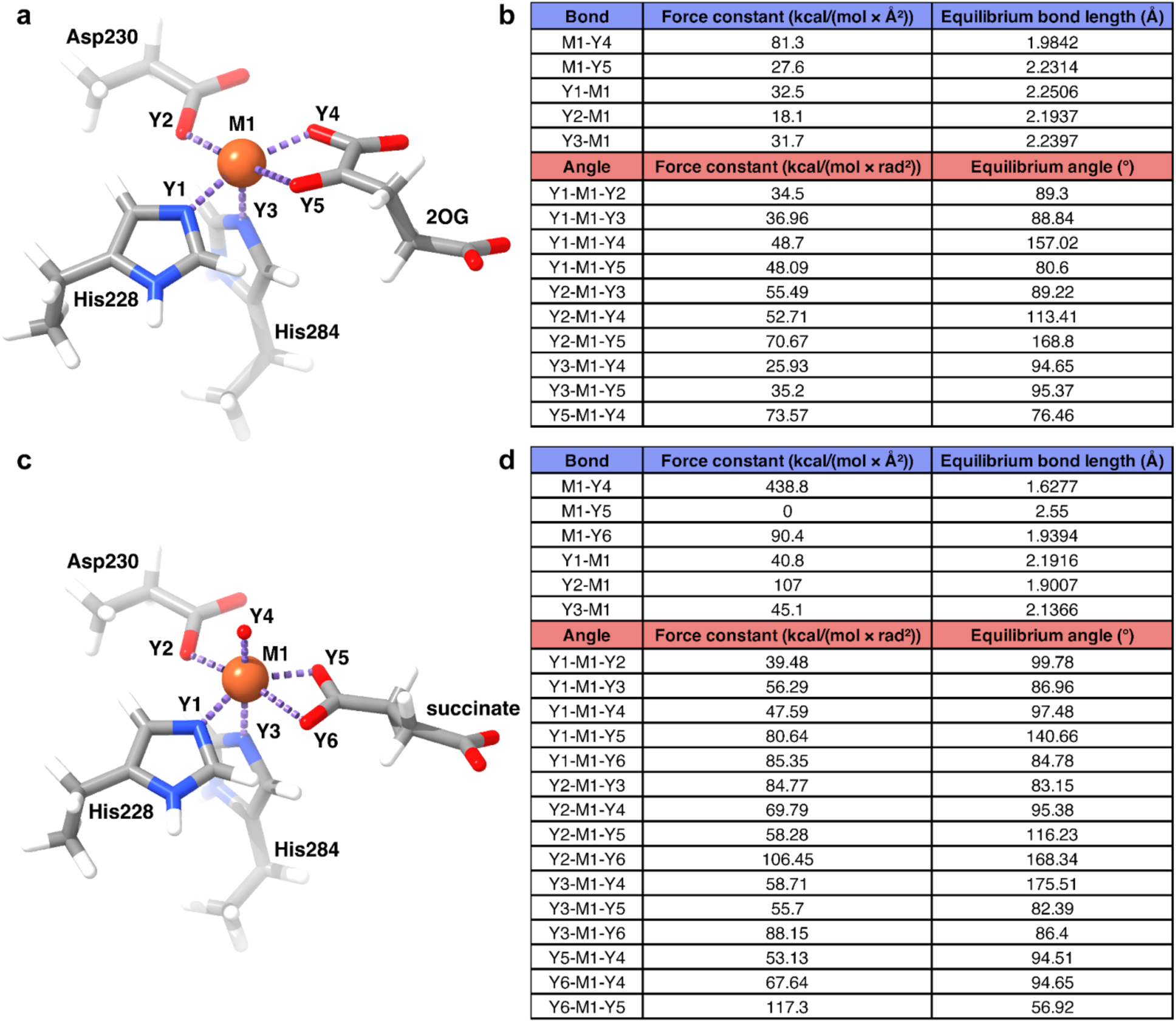
The core active site of *Mc*FLS structural models from AMBER’s Metal Center Parameter Builder (MCPB). FLS with (**a**) 2OG or (**c**) succinate in the MCPB small model with atom assignments labeled on the active site for coordinating atoms. Dative bonds are illustrated with purple dashed lines. Atoms are colored as follows: carbon in gray, nitrogen in blue, oxygen in red, hydrogen in white, and iron in orange. Parameters generated from FLS with (**b**) 2OG or (**d**) succinate force field parameters assigned using the metal center parameter builder (MCPB.py) from AmberTools22. Only the equilibrium bond lengths, angles, and their force constants that are directly bonded to the metal center are reported. Parameters were generated using the Seminario method.

**Supplementary Fig. 23.**
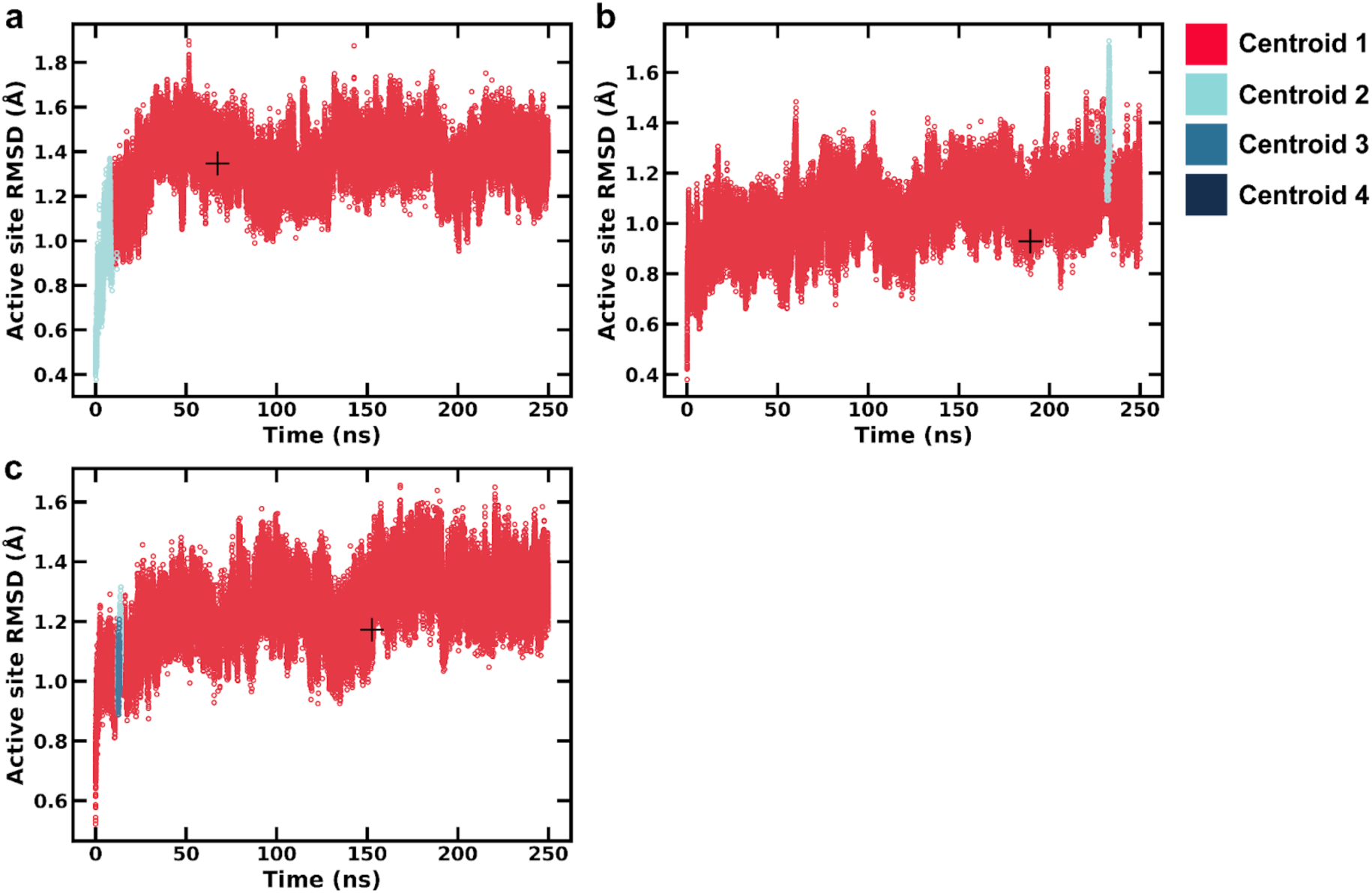
|. Clustered restrained MD simulations for FLS. The results of 250 ns MD simulations represented as time (ns) vs root mean squared deviation (Å) to the first frame. The simulations were clustered using CPPTraj and the DBSCAN method. (**a**) FLS 2OG simulation. Default clustering parameters were used with minpoints = 25 and = 0.62. (**b**) FLS succinate unrestrained simulation. Default clustering parameters were used with minpoints = 25 and = 0.66. (**c**) FLS succinate acute simulation. Default clustering parameters were used with minpoints = 25 and = 0.70. The centroid of the largest cluster is marked with a crosshair.

**Supplementary Fig. 24.**
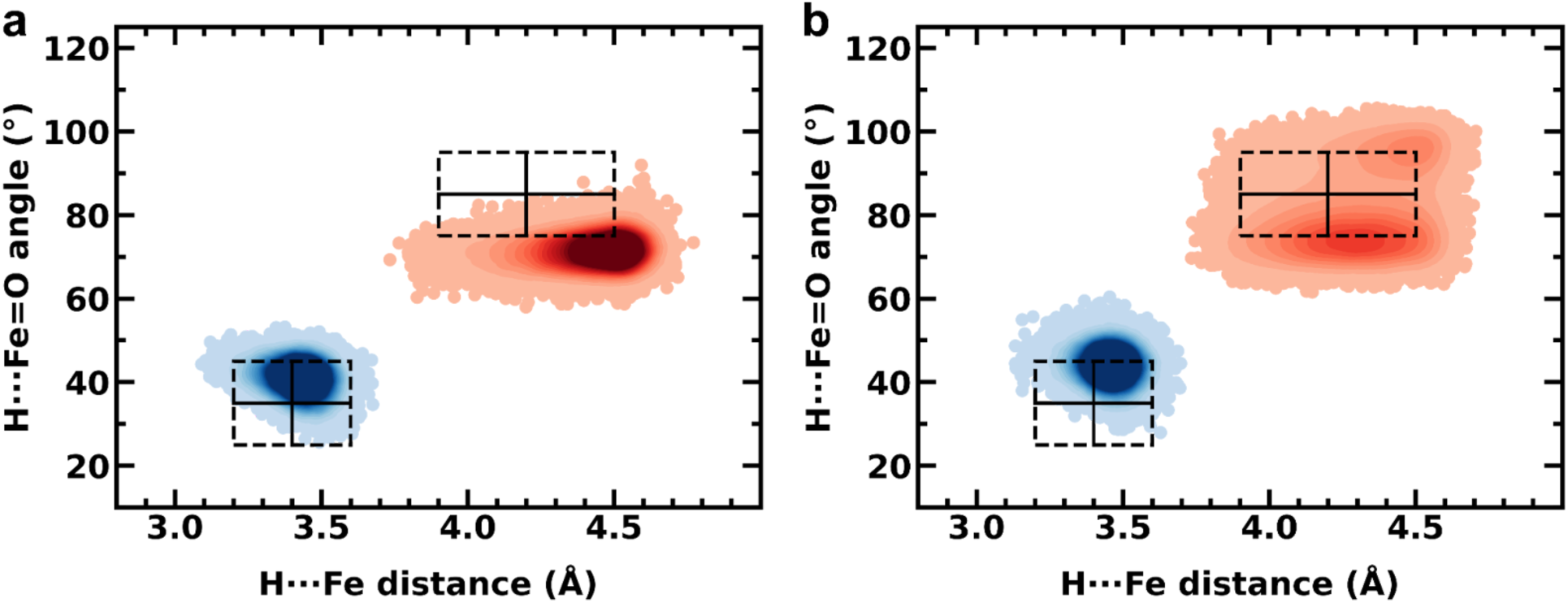
Comparison of angle and distance preferences for DAH using spectroscopically guided MD simulations. 250 ns of MD simulations were performed with the oxo in either the (**a**) axial position or (**b**) the equatorial position. The MD simulations were ran with either hydroxylase or halogenase restraints applied with restraints of 100 kcal/(mol · rad^2^) for the H···Fe=O angle (°) and 100 kcal/(mol · Å^2^) for the H···Fe distance (Å). The H···Fe=O angle and the H···Fe distance were measured with CPPTraj. Experimental HYSCORE data^3^ was used for hydroxylase-inspired restraints (blue KDE) or halogenase-inspired restraints (red KDE). The experimental target angles for the hydroxylases (OH) and halogenases (Cl) are indicated with a dotted box and crosshairs.

**Supplementary Fig. 25.**
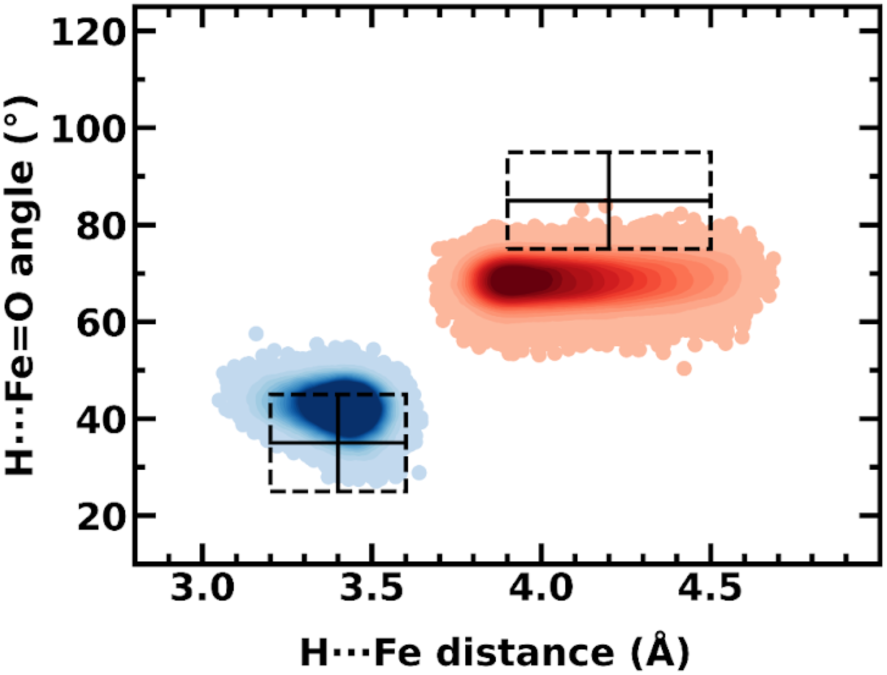
Comparison of angle and distance preferences for FLS using spectroscopically guided MD simulations. 250 ns of MD simulations were performed with either hydroxylase or halogenase restraints applied with restraints of 100 kcal/(mol · rad^2^) for the H···Fe=O angle (°) and 100 kcal/(mol · Å^2^) for the H···Fe distance (Å). The H···Fe=O angle and the H···Fe distance were measured with CPPTraj. Experimental HYSCORE data^4^ was used for hydroxylase-inspired restraints (blue KDE) or halogenase-inspired restraints (red KDE). The experimental target angles for the hydroxylases (OH) and halogenases (Cl) are indicated with a dotted box and crosshairs.

**Supplementary Fig. 26.**
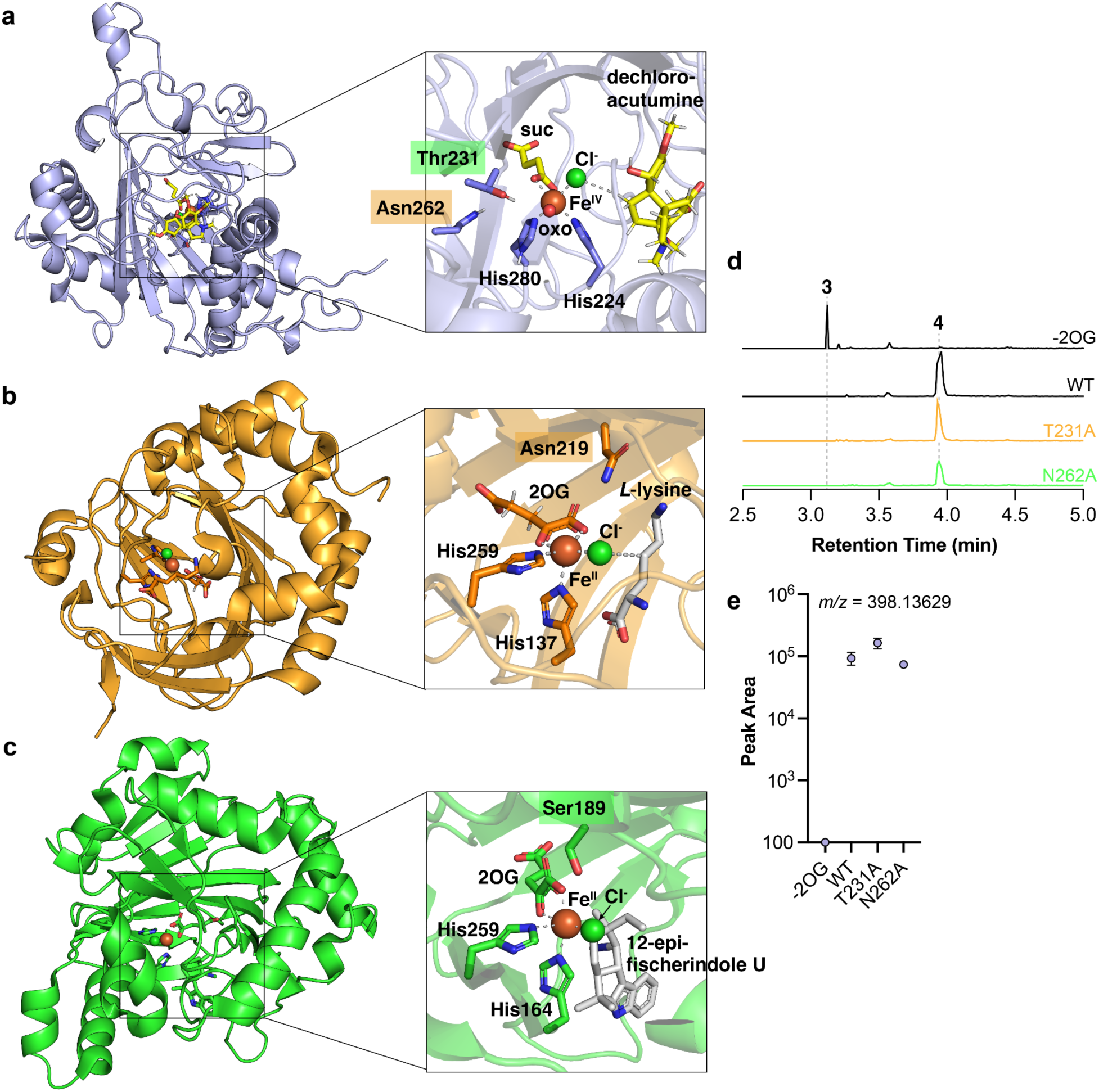
Structural and functional comparison between plant and bacterial 2ODHs. Structures of (**a**) plant 2ODH, *Mc*DAH (with the oxo in the equatorial position) compared with bacterial 2ODHs (**b**) *Sc*BesD and (**c**) *Hw*WelO5. Catalytically essential residues in *Sc*BesD (Asn219) and *Hw*WelO5 (Ser189) and their counterpart residues in *Mc*DAH are highlighted. (Abbreviation; suc: succinate, 2OG: 2-oxoglutarate) (**d**) Extracted ion chromatograms (EICs) of acutumine, 398.13629 *m/z* ; [M+H]^+^ of **4** for *Mc*DAH WT enzyme assay performed without and with 2-oxoglutarate (black), *Mc*DAH T231A mutant assay (orange), and *Mc*DAH N262A (green). (**e**) Integrated peak area of **4** from the chromatograms shown in **d**. All assays were performed in triplicates and the error bars represent standard error of the mean.

**Supplementary Fig. 27.**
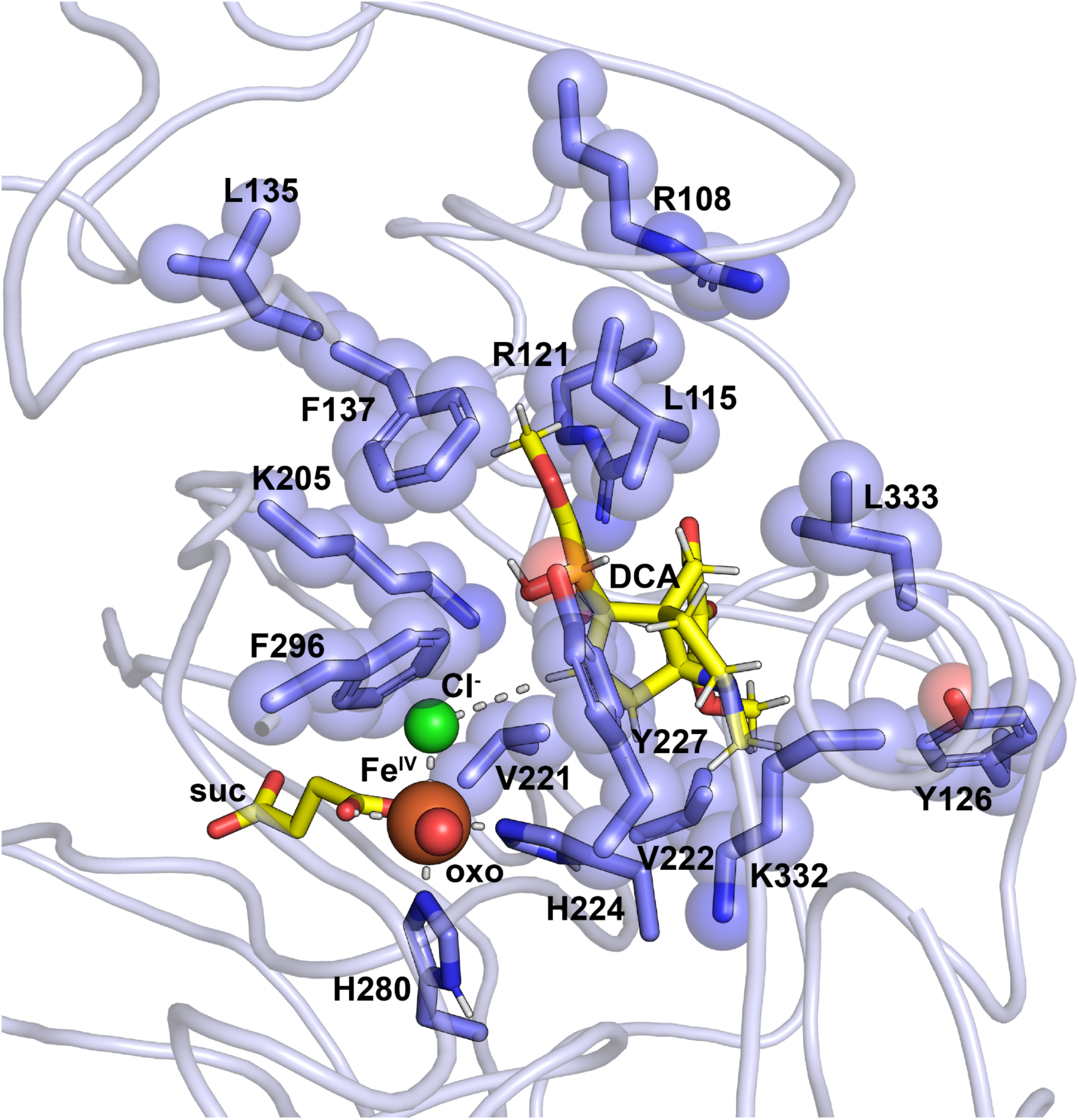
Active-site-lining residues around the substrate binding pocket in *Mc*DAH. Relevant residues chosen in this study for alanine scanning portrayed in equatorial-oxo conformation of *Mc*DAH model. (Abbreviation; suc: succinate, DCA: dechloroacutumine)

**Supplementary Fig. 28.**
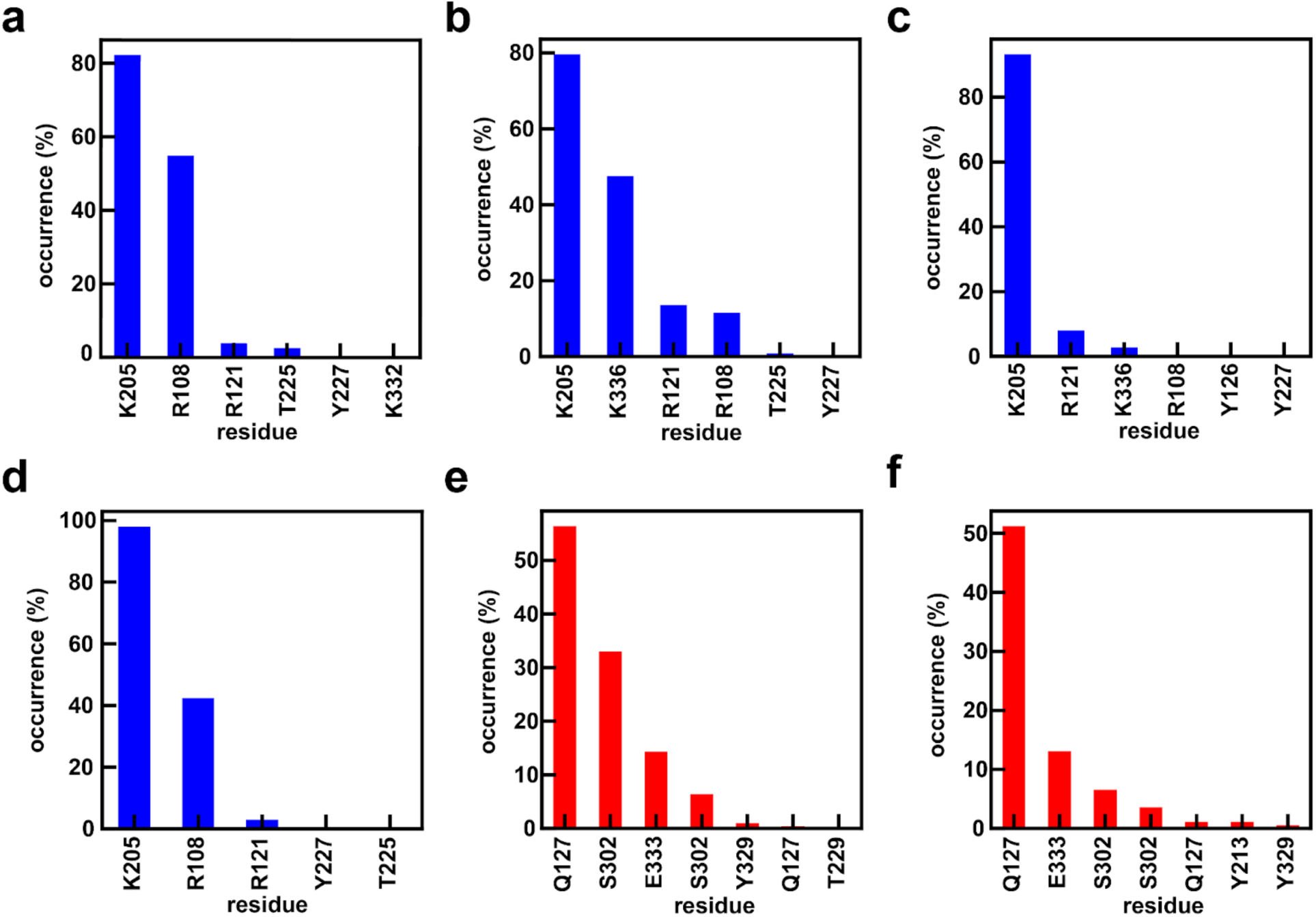
|. Comparison of the hydrogen bonding occurrence (%) summed over any atoms belonging to either interaction partner between DAH residues and dechloroacutumine (blue bars) in the following configurations: (**a**) axial-oxo unrestrained with succinate, (**b**) axial-oxo restrained obtuse with succinate, (**c**) equatorial-oxo unrestrained with succinate, and (**d**) equatorial-oxo restrained obtuse with succinate. Bars in each plot are ordered by decreasing frequency of hydrogen bonding occurrence. Comparison of hydrogen bonding occurrence (%) between FLS residues and dihydrokaempferol (red plots) for the following configurations: (**e**) unrestrained with succinate and (**f**) restrained acute with succinate. For the restrained simulations, harmonic restraints of 100 kcal/(mol·rad^2^) were employed for the angle between the oxo, the iron, and the hydrogen atom target. Harmonic restraints of 100 kcal/(mol·Å^2^) were used for the distance between iron and the hydrogen atom target. Hydrogen bonding occurrence is obtained based on the default CPPTraj geometric criteria and a modified distance cutoff of 3.2 Å.

**Supplementary Fig. 29.**
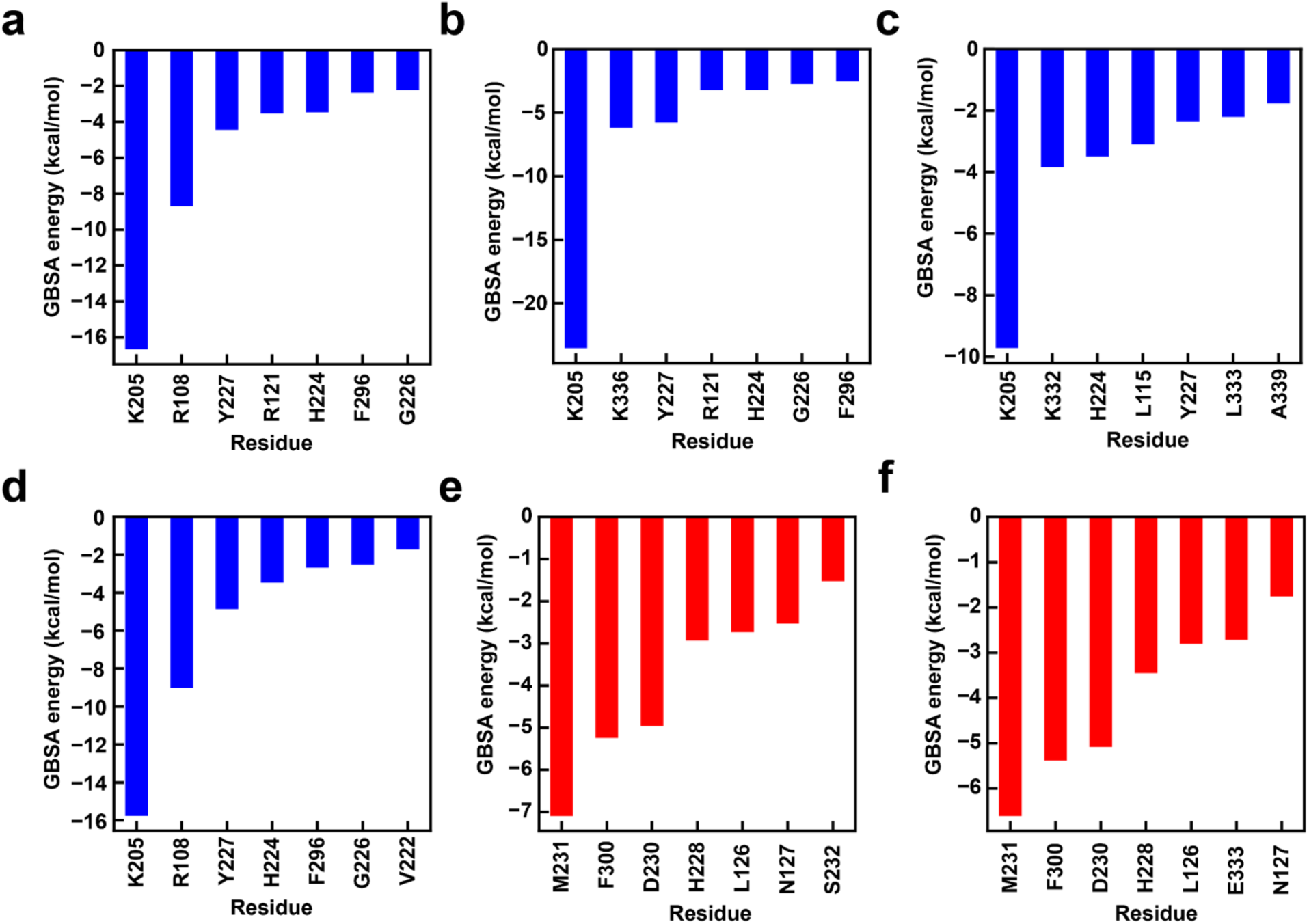
| **Comparison of the Generalized Born surface area (GBSA) analysis in kcal/mol.** Contributions to the non-covalent interactions are shown as the energetic sum of the interaction components, vDW, electrostatic, polar, and non-polar, between DAH residues and dechloroacutumine (blue bars) in the following configurations: (**a**) axial-oxo unrestrained with succinate, (**b**) axial-oxo restrained obtuse with succinate, (**c**) equatorial-oxo unrestrained with succinate, (**d**) equatorial-oxo restrained obtuse with succinate. Comparison of GBSA of the classical interactions between FLS residues and dihydrokaempferol (red bars) for the following configurations: (**e**) unrestrained with succinate and (**f**) restrained acute with succinate. GBSA was performed on 1000 snapshots from the primary cluster of DBSCAN-clustered restrained MD simulations. Snapshots were taken 50 ps frames apart as described previously.^1^ No entropy correction was applied, and the Generalized Born model with the Onufriev, Bashford, and Case (OBC) effective radii set 1 (igb=2) was employed.^2^

**Supplementary Fig. 30.**
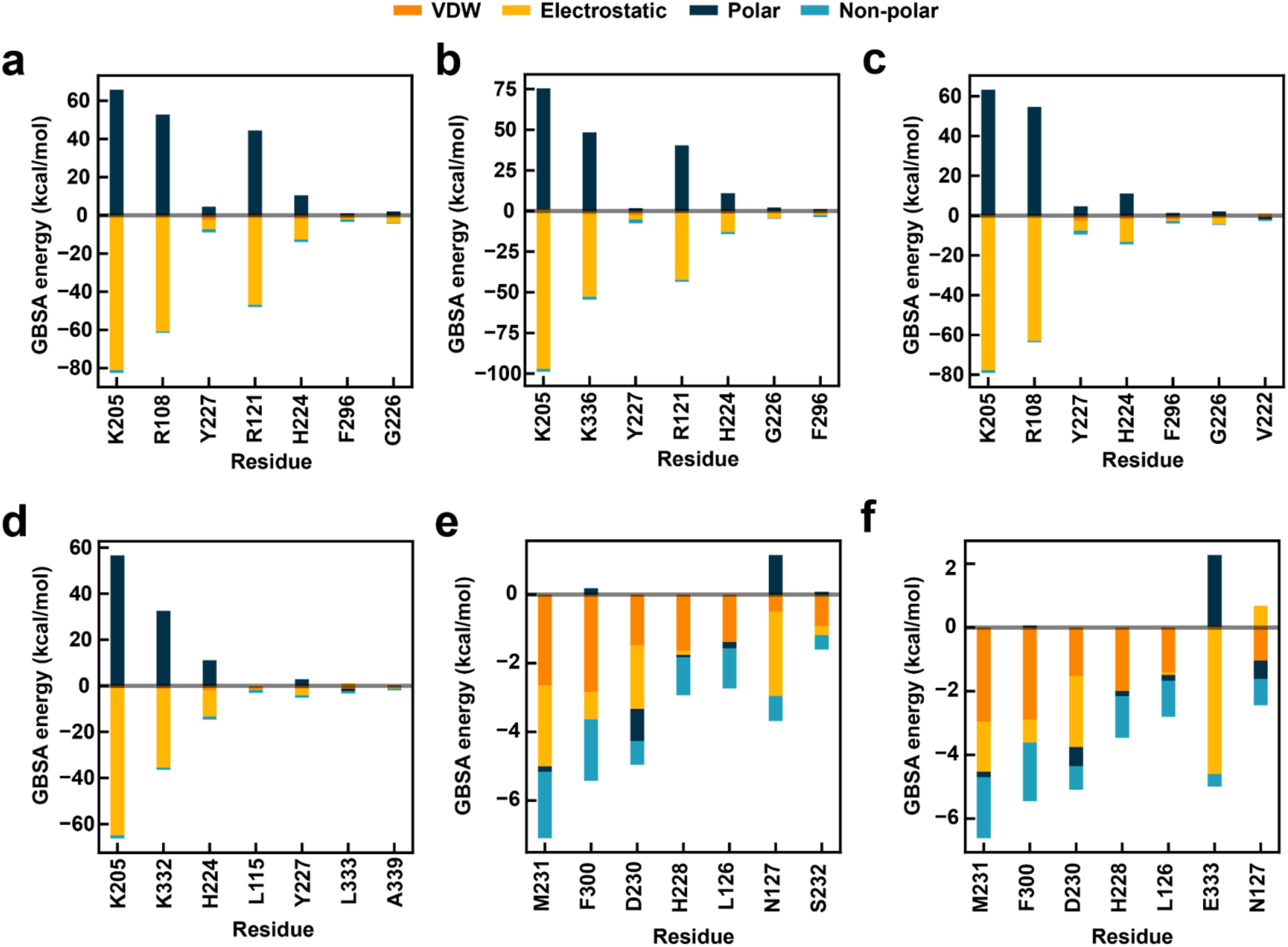
Comparison of the GBSA interaction energies (in kcal/mol). Contributions to the non-covalent interactions are broken up by VDW, electrostatic, polar, and non-polar interaction components for the classical interactions between DAH residues and dechloroacutamine in the following configurations: (**a**) axial-oxo unrestrained with succinate, (**b**) axial-oxo restrained obtuse with succinate, (**c**) equatorial-oxo unrestrained with succinate, (**d**) equatorial-oxo restrained obtuse with succinate. Comparison of the GBSA of the classical interactions between FLS residues and dihydrokaempferol for the following configurations: (**e**) unrestrained with succinate and (**f**) restrained acute with succinate. GBSA was performed on 1000 snapshots from the primary cluster of DBSCAN-clustered restrained MD simulations. Snapshots were taken 50 ps frames apart as described previously.^1^ No entropy correction was applied, and the Generalized Born model with the Onufriev, Bashford, and Case (OBC) effective radii set 1 (igb=2) was employed.^2^

**Supplementary Fig. 31.**
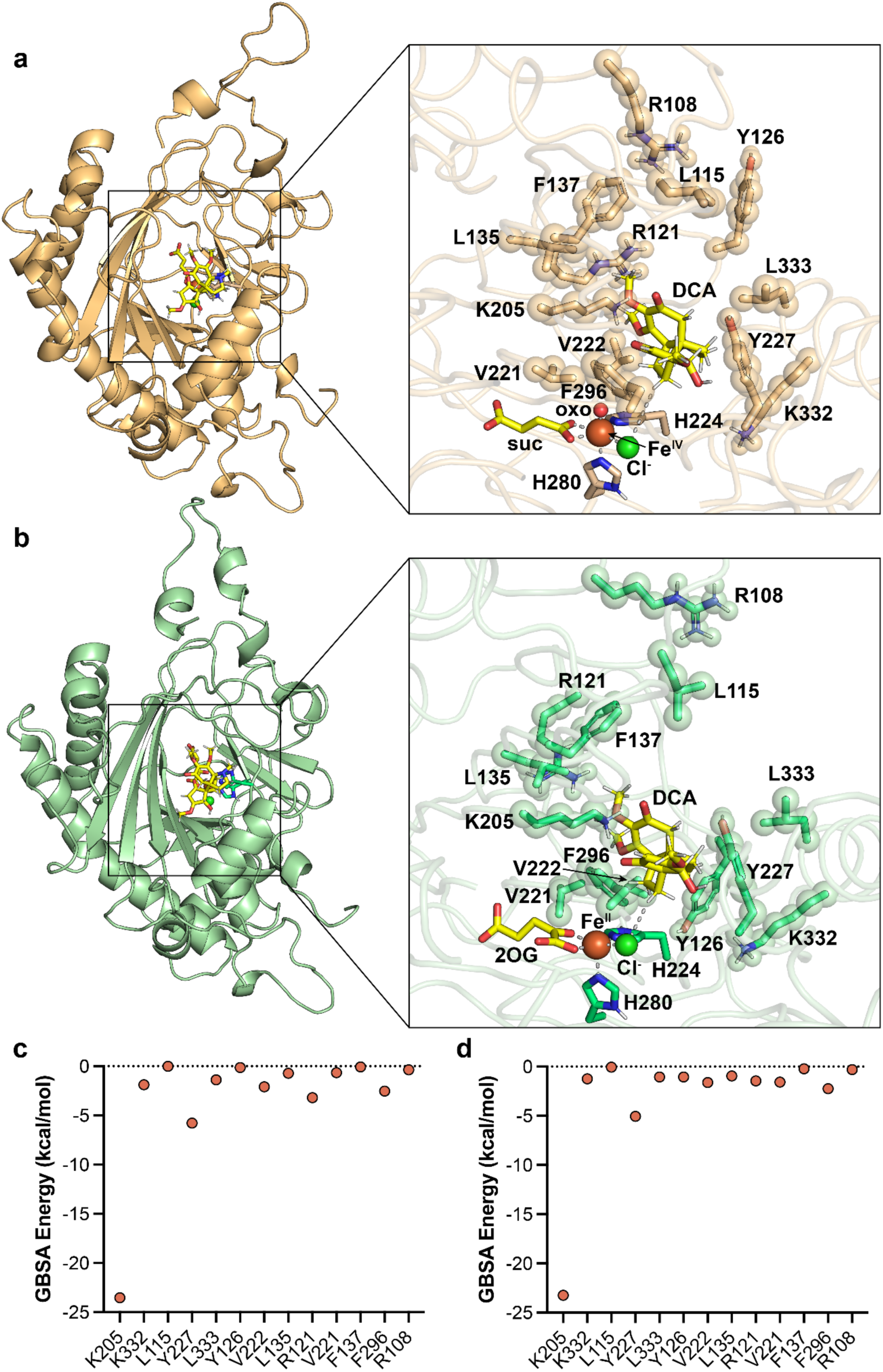
MM-GBSA hydrogen-bonding energy calculations from MD simulations of *Mc*DAH structural models. (**a**) *Mc*DAH structural model with the oxo in the axial position portraying relevant residues chosen in this study for alanine scanning. (**b**) 2OG-bound *Mc*DAH structural model portraying relevant residues chosen in this study for alanine scanning. (In both panels **a** and **b**– Abbreviation; suc: succinate, DCA: dechloroacutumine) (**c**) MM-GBSA hydrogen-bonding energy calculation for *Mc*DAH structural model with the oxo in the axial position. (**d**) MM-GBSA hydrogen-bonding energy calculation for *Mc*DAH structural model with 2OG bound. In both panels **c** and **d**, the x-axis is ordered by low-to-high GBSA energy calculation based on the *Mc*DAH structural model with the oxo in the equatorial position.

**Supplementary Fig. 32.**
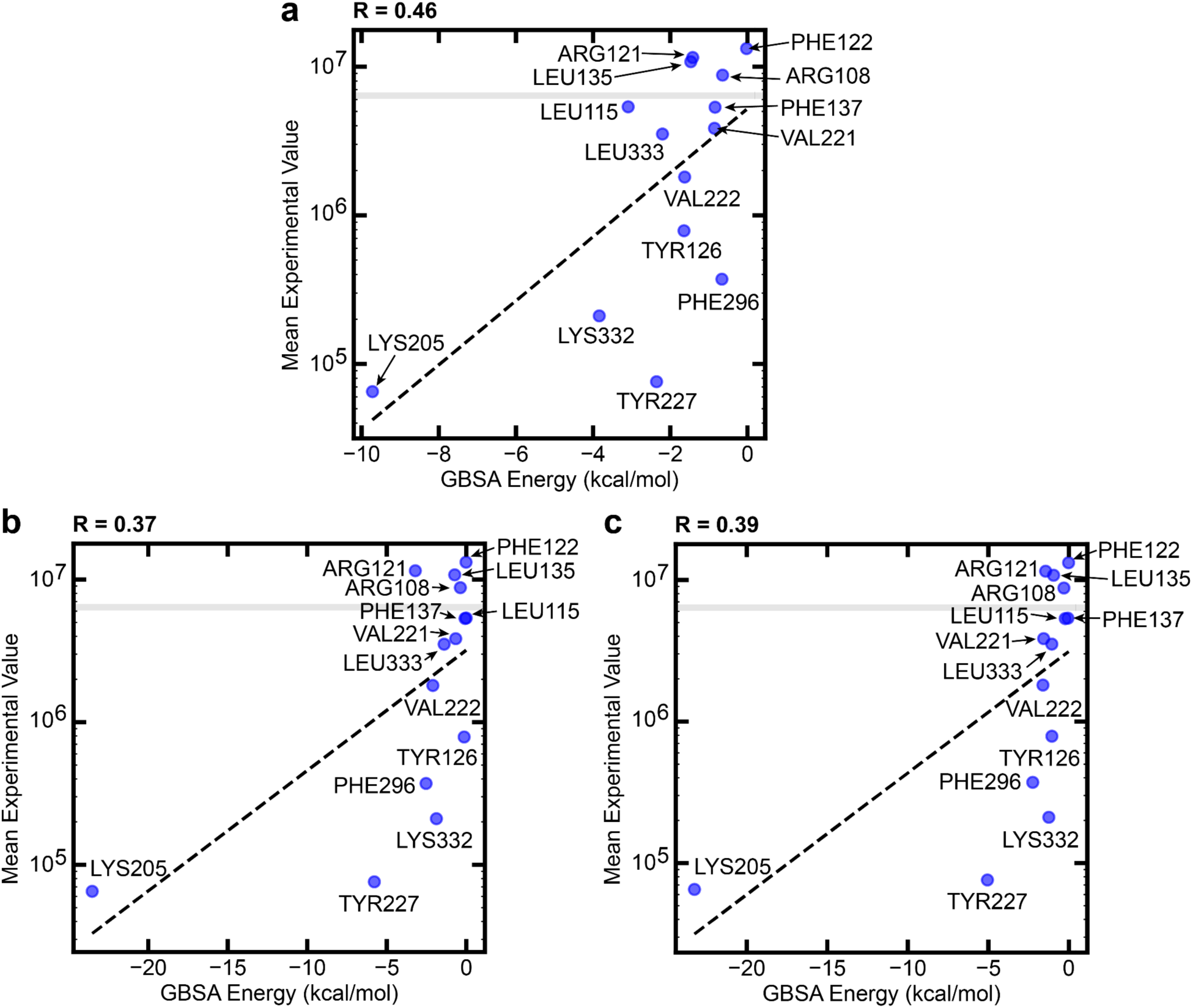
Parity plots comparing the MM-GBSA hydrogen-bonding energy calculations to *Mc*DAH alanine mutant assay data. (**a**) Comparison of the *Mc*DAH structural model with the oxo in the equatorial position with the corresponding alanine mutant experimental data. (**b**) Comparison of the *Mc*DAH structural model with the oxo in the axial position with the corresponding alanine mutant experimental data. (**c**) Comparison of the 2OG-bound *Mc*DAH structural model with the corresponding alanine mutant experimental data. For all panels, the y-axis is shown as log_10_-scale of LC-HRAM-MS peak areas of acutumine ([M+H]^+^ = 398.13629 *m/z*). The R-value represents Pearson’s correlation coefficient comparing the x-axis and y-axis values. Grey lines in each panel indicate the mean peak area of acutumine measured for WT *Mc*DAH.

**Supplementary Fig. 33.**
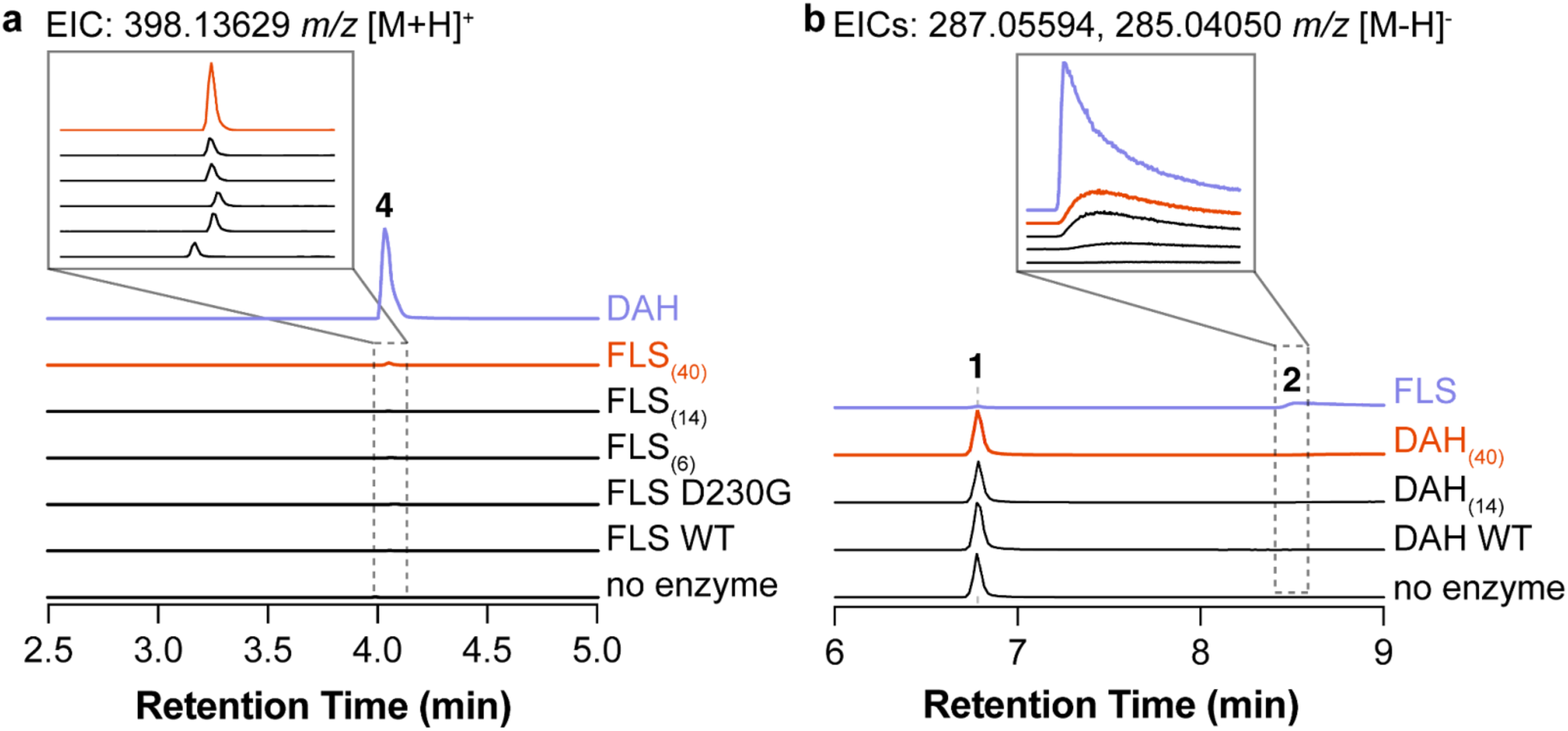
Qualitative LC-HRAM-MS chromatogram of *Mc*FLS mutants for minimal *Mc*DAH activity. (**a**) Extracted ion chromatograms (EICs) of acutumine, 398.13629 *m/z* ; [M+H]^+^ of **4**. Retention time shift is observed in no enzyme sample due to changes in the chromatography buffer and as it was run independently from other samples present in this figure. Trace amounts of acutumine is detected in the control samples due to its presence from dechloroacutumine substrate used in the assays. (**b**) Combined extracted ion chromatograms (EICs) of dihydrokaempferol, 287.05594 *m/z*; [M-H]^-^ of **1** and kaempferol, 285.04050 *m/z*; [M-H]^-^ of **2**. DAH_(14)_ enzyme has the β-sheet loop and substrate positioning loop regions swapped with their sequence alignment counterparts of *Mc*FLS. DAH_(40)_ enzyme has the β-sheet loop, substrate positioning loop, and C-terminal helical loop regions swapped with their counterparts of *Mc*FLS sequence.

**Supplementary Fig. 34.**
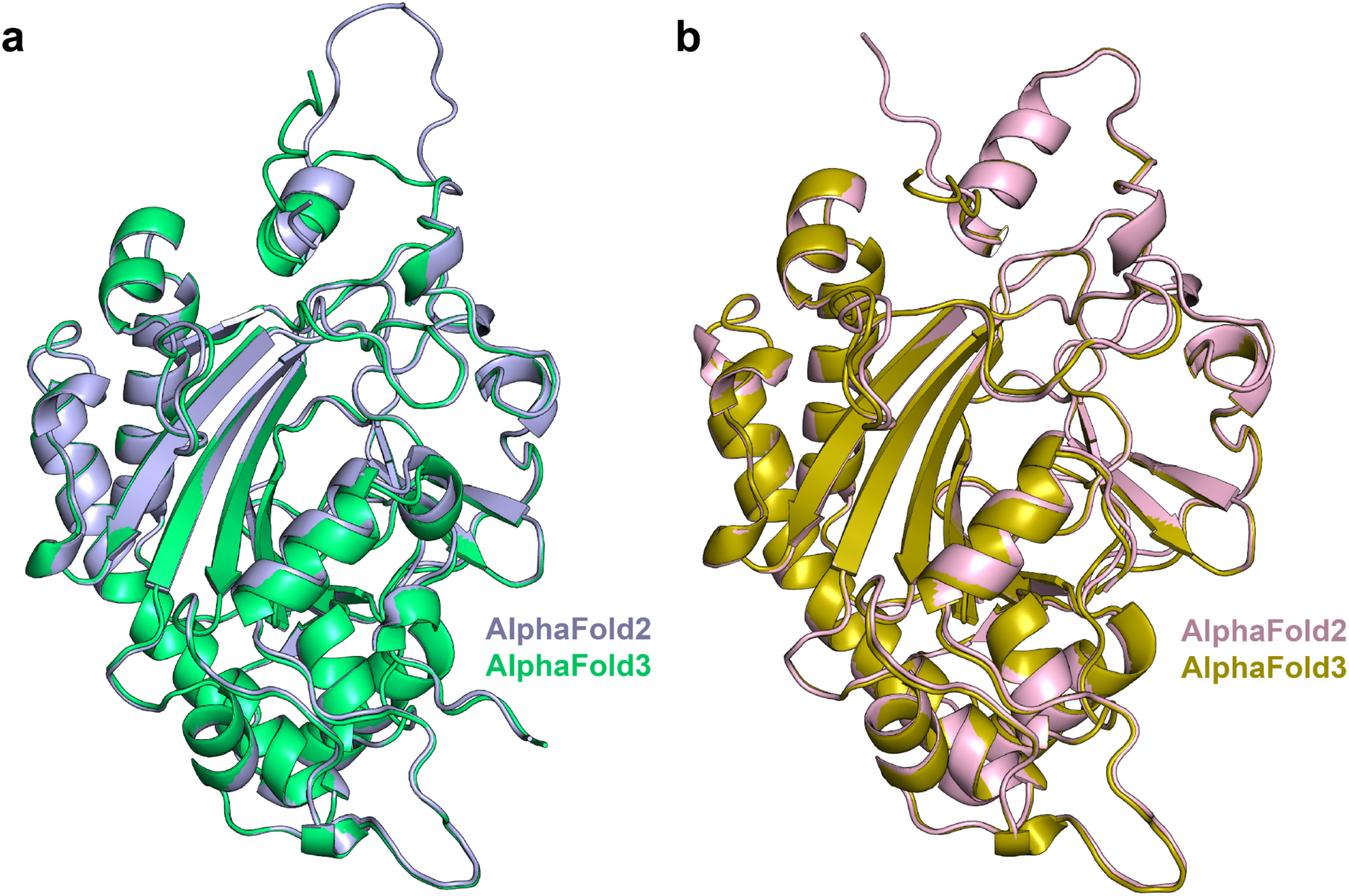
Structural comparison of AlphaFold2 and AlphaFold3 structural models. (**a**) DAH structural models from AlphaFold2 (light blue) and AlphaFold3 (green). Structural alignment resulted in an RMSD value of 0.295 for 2301 atoms.(**b**) FLS structural models from AlphaFold2 (pink) and AlphaFold3 (dark yellow). Structural alignment resulted in an RMSD value of 0.318 for 2266 atoms.

